# Alteration of cardiolipin-dependent mitochondrial coupling in muscle protects against obesity

**DOI:** 10.1101/715953

**Authors:** Alexandre Prola, Jordan Blondelle, Aymeline Vandestienne, Jérôme Piquereau, Raphaël GP Denis, Stéphane Guyot, Hadrien Chauvin, Arnaud Mourier, Martine Letheule, Marie Maurer, Céline Henry, Nahed Khadhraoui, Guillaume Courtin, Nicolas Blanchard-Gutton, Laurent Guillaud, Inès Barthélémy, Mélanie Gressette, Audrey Solgadi, Florent Dumont, Julien Castel, Julien Ternacle, Jean Demarquoy, Alexandra Malgoyre, Nathalie Koulmann, Geneviève Derumeaux, Marie-France Giraud, Stéphane Blot, Frédéric Joubert, Vladimir Veksler, Serge Luquet, Frédéric Relaix, Laurent Tiret, Fanny Pilot-Storck

## Abstract

The tubular shape of mitochondrial cristae depends upon a specific composition of the inner mitochondrial membrane, including cardiolipin that allows strong curvature and promotes optimal organization of ATP synthase. Here we identify *Hacd1,* which encodes an enzyme involved in very long chain fatty acid biosynthesis, as a key regulator of composition, structure and functional properties of mitochondrial membranes in muscle. In *Hacd1*-deficient mice, the reduced cardiolipin content was associated with dilation of cristae and caused defective phosphorylating respiration, characterized by absence of proton leak and oxidative stress.

The skeletal muscle-specific mitochondrial coupling defect produced a global elevation in basal energy expenditure with increased carbohydrate and lipid catabolism, despite decreased muscle mass and locomotor capacities. Mice were protected against diet-induced obesity despite reduced muscle activity, providing an *in vivo* proof of concept that reducing mitochondrial coupling efficiency in skeletal muscle might be an actionable mechanism in metabolic disease conditions.

## INTRODUCTION

Skeletal muscles play a pivotal role in metabolism by their capacity to produce the energy required to achieve contraction. They represent the major energy sink of the body (Rolfe and Brown, 1997), and muscle mitochondria use chemical substrates such as carbohydrates and fatty acids as fuel to generate the transmembrane proton gradient and phosphorylate ADP (Mitchell, 1961). Through this oxidative phosphorylation coupling, ATP is produced and promotes sustainable cross-bridge cycling in muscles. Continual ATP production is guaranteed by convergent signaling regulators such as the insulin and AMPK pathways, combined with a complex regulation of both the content and activity of mitochondria (Mouchiroud et al., 2014). In particular, the highly specialized and essential function of mitochondria relies on a very specific organization of its double membrane, reminiscent of its endosymbiotic origin (Zimorski et al., 2014). Notably, the inner mitochondrial membrane (IMM) displays a massive surface extension by folding into cristae, along which respiratory complexes and oligomers of ATP synthase concentrate in a stratified proximodistal pattern (Vafai and Mootha, 2012). Evidence has accumulated to highlight the importance of a specific lipid composition of the IMM to promote stability of the strongly curved structure of cristae, which is essential to their function. Accordingly, the IMM is enriched in cardiolipin (CL), a four-acyl chain phospholipid required to build tubular-shaped cristae (Paradies et al., 2014). In the striated, *i.e.* skeletal and cardiac, muscles characterized by very high oxidative capacities, remodeling of CL with C18 fatty acids catalyzed by Tafazzin proved to be essential for their normal function (Barth et al., 1983; Ikon and Ryan, 2017a). Fatty acids with ≥ C18 acyl chains are synthesized by the very long chain fatty acids (VLCFA) elongation cycle that involves four steps catalyzed by endoplasmic reticulum-resident enzymes (Denic and Weissman, 2007; Ikeda et al., 2008; Kihara, 2012). HACD (3-hydroxyacylCoA dehydratase) proteins catalyze the third step of this elongation cycle and four *HACD* paralogous genes are present in mammals, with specific expression patterns (Blondelle et al., 2015; Ikeda et al., 2008; Wang et al., 2004). Notably, the full-length isoform of *HACD1* (*HACD1*-*fl*), which encodes the catalytically active isoform of HACD1, is mainly expressed in striated muscles (Blondelle et al., 2015). We and others identified the essential role of *HACD1* in muscle homeostasis through the characterization of loss-of-function mutations in dogs, humans and mice, all leading to congenital myopathies characterized by reduced muscle mass and strength (Blondelle et al., 2015; Muhammad et al., 2013; Pelé et al., 2005; Toscano et al., 2017). We demonstrated that *HACD1* is required for efficient myoblast fusion during muscle postnatal development and as a consequence, HACD1-deficient dogs and mice display an early and uncompensated reduced muscle mass throughout their lifespan (Blondelle et al., 2015). However, integrated mechanisms of muscle weakness remain undeciphered. Fatty acids derived from the VLCFA elongation cycle contribute to membrane properties and are the main components of CL that are specifically enriched in mitochondria of striated muscles; hence, we hypothesized that *Hacd1* is involved in the constitution of a functional CL pool thereby contributing to the genetic regulation of their outstanding oxidative capacities.

Using HACD1-deficient mice and dogs, we provide here evidence that *Hacd1* is a major nuclear regulatory gene of mitochondrial membrane homeostasis and coupling efficiency in skeletal muscles. Our results further demonstrate that yielding a defective oxidative phosphorylation specifically in skeletal muscle mitochondria is an efficient mechanism for protecting mice against energetic overload conditions.

## RESULTS

### HACD1-deficient mice display increased glucose tolerance and insulin sensitivity

In mice and dogs, HACD1 deficiency (*Hacd1*-KO mice and CNM dogs) leads to reduced muscle mass and strength (Blondelle et al., 2015; Pelé et al., 2005). The hypotrophy arises during postnatal muscle development and remains stable thereafter, whereas the consequences of HACD1 deficiency on muscle function and metabolism remained undeciphered. We recorded spontaneous locomotor activity and speed in *Hacd1*-KO mice and showed they were reduced by a half and a third respectively when compared to their wild-type littermates (WT) (Figures 1A and S1A). A treadmill test was performed to more specifically assess their skeletal muscle functional capacities and the maximal aerobic speed of *Hacd1*-KO mice was significantly reduced compared to WT (Figure 1B). More precisely, we identified that - when challenged with submaximal aerobic exercise - *Hacd1-*KO mice ran a shorter maximal distance (−40%) during a shorter period (time to exhaustion was reduced by 37%), however their maximal aerobic speed was reduced by only 13,8% (Figures 1B, 1C and S1B). This demonstrated that *Hacd1-*KO mice have reduced endurance capacity reminiscent of the exercise intolerance and fatigue observed in CNM dogs from the onset of clinical signs (https://tinyurl.com/cnmdogs).

**Figure 1.**
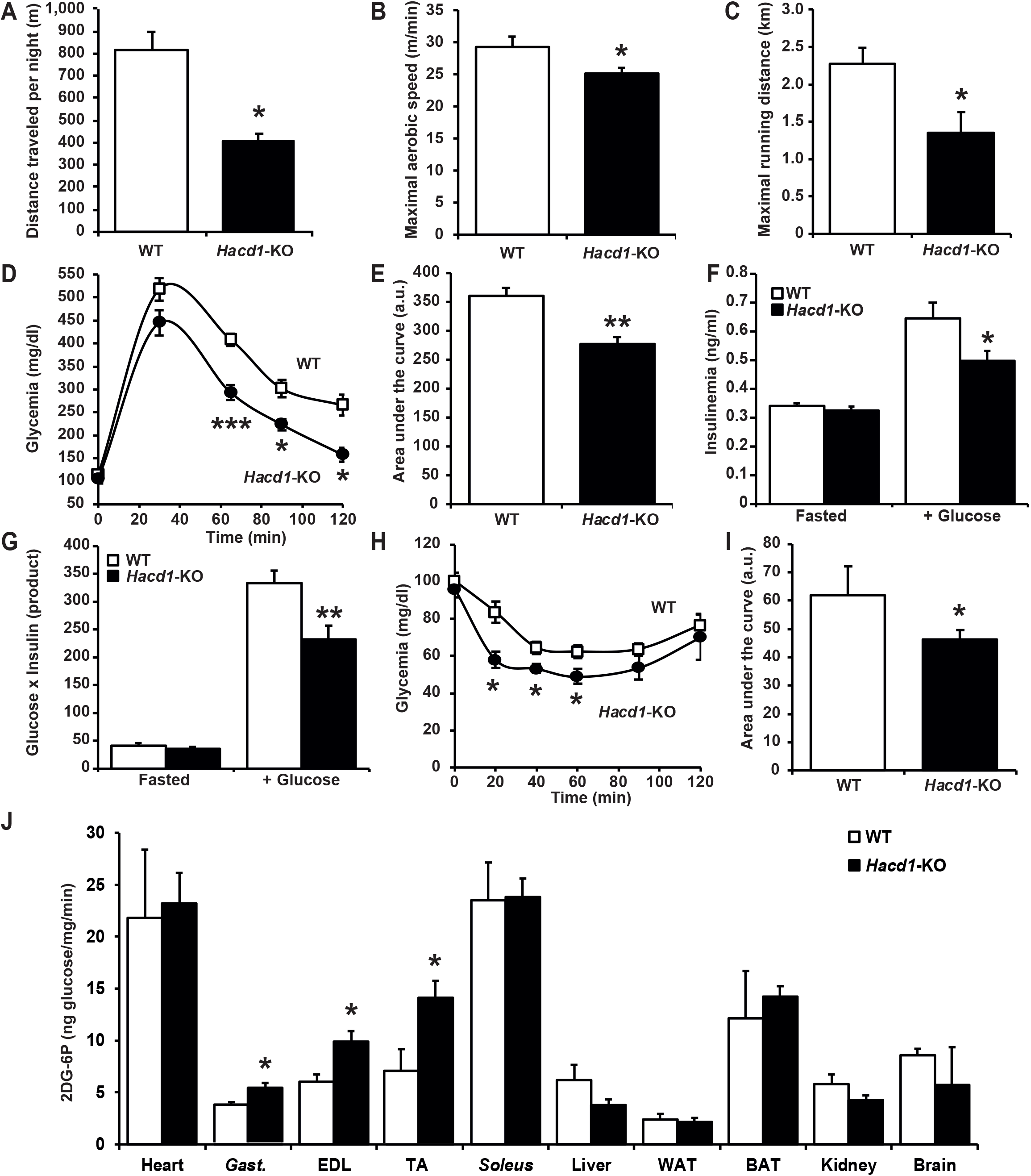
Increased glucose clearance in hypokinetic *Hacd1*-KO mice. **(A)** Distance traveled per night. **(B)** Maximal aerobic speed during a treadmill test. **(C)** Maximal running distance at 60% of maximal aerobic speed. **(D)** Glycemia measured after fasting and assessed during 120 min after an intraperitoneal glucose injection at T_0_ (glucose tolerance test). **(E)** Area under the glycemia curves (arbitrary units, a.u.) displayed in **(D)**. **(F)** Insulinemia measured after fasting (Fasted) and 30 min after an intraperitoneal glucose injection (+ Glucose). **(G)** Insulinemia x Glycemia product (ng/ml x mg/dl) in the two conditions displayed in **(F)**. **(H)** Glycemia measured after food deprivation and assessed during 120 min after an intraperitoneal insulin injection at T_0_ (insulin sensitivity test). **(I)** Area under the glycemia curves displayed in **(H)**. (**J**) [^14^C]-2-deoxy-D-glucose-6P content in heart, superficial *gastrocnemius* (*Gast*.), *extensor digitorum longus* (EDL), *tibialis anterior* (TA), *soleus*, liver, white adipose tissue (WAT), brown adipose tissue (BAT), kidney and brain in WT and *Hacd1*-KO mice, 120 min after [^14^C]-2-deoxy-D-glucose. *n* = 4 per group for **A**, *n* = 8 per group for **B-C**, *n* = 7 per group for **D-I**; *n* = 6 per group for **J**; error bars: ± s.e.m.; \**P* < 0.05, \*\**P* < 0.01 and \*\*\**P* < 0.001.

Skeletal muscle is the body’s largest sink for glucose in response to insulin (Srikanthan and Karlamangla, 2011); hence, we hypothesized that reduced muscle mass and activity in *Hacd1-*KO mice would impair their ability to produce an efficient response to a systemic glucose overload. Surprisingly, starting from comparable fasting blood glucose and insulin levels (Figure 1D and 1F), *Hacd1-*KO mice exhibited higher glucose clearance (Figures 1D and 1E) despite reduced insulinemia (Figure 1F and 1G) after an intraperitoneal glucose bolus. This indicated a higher insulin sensitivity, further evidenced by a more pronounced hypoglycemia after an insulin injection in *Hacd1-*KO mice, compared to WT (Figures 1H and 1I). The labelled [^14^C]-2-deoxy-D-glucose ([^14^C]-2-DG) was used as a tracer for glucose uptake; after intraperitoneal injection in *Hacd1-*KO and WT mice, we observed a selective accumulation of [^14^C]-2-DG-6P in *Hacd1*-KO superficial *gastrocnemius*, *extensor digitorum longus* and *tibialis anterior* muscles contrasting with comparable contents in heart, liver, white adipose tissue, brown adipose tissue, kidney and brain in *Hacd1*-KO and WT mice (Figure 1J). These findings pointed to a specific increase of glucose uptake in skeletal muscles of *Hacd1*-KO mice. Glucose uptake relies on both the insulin pathway and the glucose gradient across the sarcolemma, which is increased during exercise by the rise in catabolism (Richter and Hargreaves, 2013). After insulin injection, the phosphorylation level of insulin receptor β (INSRβ), insulin receptor substrate 1 (IRS1) and AKT, and the expression of GLUT4 were comparable in *gastrocnemius* muscle of WT and *Hacd1-*KO mice (Figures S1C and S1D), favoring the hypothesis of a higher glucose gradient in *Hacd1*-KO mice driven by glucose catabolism, rather than overactivation of the insulin-signaling pathway. Of note, [^14^C]-2-DG-6P that cannot enter the glycolysis pathway inhibits glucose hexokinase (Chen and Guéron, 1992), thus, accumulation of [^14^C]-2-DG-6P likely counteracted the phosphorylation-driven glucose gradient, yielding the normal blood clearance of [^14^C]-2-DG observed in *Hacd1*-KO mice (Figure S1E and S1F).

Routine staining of transverse sections of HACD1-deficient muscles revealed no glycogen accumulation (data not shown), supporting a facilitated entry of glucose into catabolic pathways. Markers of anaerobic glycolysis (such as lactate dehydrogenase and glycero-3-phosphate dehydrogenase activities) and expression levels of the dynamically regulated *Hk2, Pdk4* and *Glut4* genes, were unaffected or reduced (Figures S1G-S1J). In contrast, markers of oxidative activity such as the succinate dehydrogenase (SDH) and NADH dehydrogenase activities were increased in muscles from both *Hacd1*-KO mice and CNM dogs (Figures S1I and S1J), revealing a shift of myofibers towards a more oxidative metabolism. In CNM dogs and in human patients, this metabolic shift has been also associated with a shift towards type I slow-twitch, oxidative fibers (Muhammad et al., 2013; Tiret et al., 2003; Toscano et al., 2017). In mice, however, the functional metabolic switch induced no modification in the expression pattern of myosin heavy chain isoforms in a highly oxidative (*soleus*), glycolytic (superficial *gastrocnemius*) or mixed (*tibialis anterior*) muscle (Figures S1K-S1O).

### HACD1 deficiency increases whole body energy expenditure

To assess additional systemic consequences of the identified muscular metabolic shift in steady state conditions, we monitored mice fed with a normal diet (ND) over nine weeks. Because of their acquired muscle hypotrophy, *Hacd1-*KO mice had a constantly reduced body mass when compared with their WT littermates (Figure 2A), but strikingly, they displayed an increased calories intake per gram of body mass (Figures 2B, 2C, S2A and S2B). To summarize, feed efficiency (*i.e.* body mass gain per ingested kcal) was markedly reduced in *Hacd1-*KO mice (−46%; Figure 2D) suggesting an elevated energy expenditure at the expense of body mass gain. This was confirmed by indirect calorimetry that revealed an increased energy expenditure relative to the lean body mass in *Hacd1-*KO mice both on normal diet (Figure 2E) and upon a short-term high-fat diet (HFD) (Figure 2F). ANOVA analysis including all animals independent of diet manipulation also revealed an increase in energy expenditure in *Hacd1*-KO mice (Figure S2C). Interestingly, calculated energy expenditure at rest - an estimation of resting metabolism - under HFD was higher in *Hacd1-*KO mice compared to WT (Figures 2G, S2D and S2E), pointing towards a better immediate adaptation to HFD, and a potential switch towards enhanced fatty acid oxidation. Indeed, under ND in which lipids represent only 12% of energy, the decreased respiratory exchange ratio (RER) during the nocturnal activity period revealed a preference for lipid substrates in *Hacd1-*KO mice, confirmed by an increased calculated fatty acid oxidation (Figures 2H-2K, S2F and S2G). Significant contributors to energy expenditure are locomotor activity and brown adipose tissue (BAT)-dependent thermogenesis; along with a reduced locomotor activity (Figure 1A), *Hacd1-*KO mice had lighter white and brown adipose tissues, expressed normal levels of transcripts encoding the UCP1 thermogenic protein and had unchanged body temperature (Table S5 and Figures S2H-S2K), excluding locomotor activity and thermogenesis as causative mechanisms for the increased energy expenditure that more likely resulted from a higher basal metabolism.

**Figure 2.**
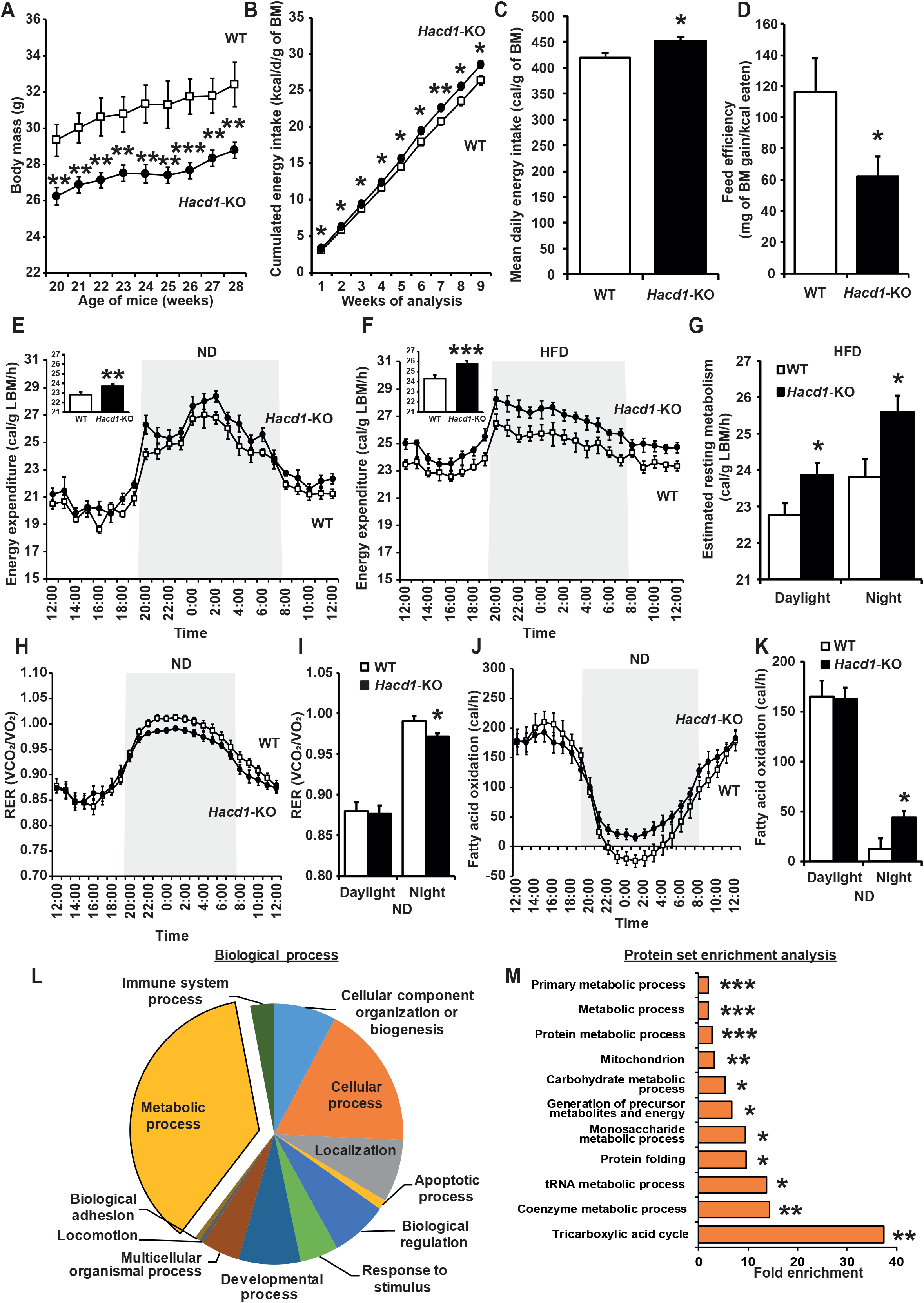
Increased energy expenditure in *Hacd1*-KO mice. **(A)** Values over a 9-wk assessment period of the body mass (BM) under normal diet, from 20 weeks of age. **(B** and **C)** Cumulated **(B)** and mean daily **(C)** energy intake over the 9-wk assessment period. **(D)** Feed efficiency during the 9-wk assessment period. **(E)** Circadian energy expenditure measured by indirect calorimetry under normal diet (ND); the active period of night is shaded. Mean hourly energy expenditure during the assessment period is represented as histogram. **(F)** Circadian energy expenditure measured by indirect calorimetry under high-fat diet (HFD); the active period of night is shaded. Mean hourly energy expenditure during the assessment period is represented as histogram. **(G)** Estimated resting metabolism during the assessment period of **(F)** during daylight and night. **(H)** Respiratory exchange ratio (VCO_2_/VO_2_) under ND measured by indirect calorimetry; the active period of night is shaded. **(I)** Respiratory exchange ratio during the assessment period of **(H)** during daylight and night. **(J)** Circadian fatty acid oxidation under ND measured by indirect calorimetry; the active period of night is shaded. **(K)** Fatty acid oxidation during the assessment period of **(J)** during daylight and night. **(L and M)** Gene ontology analysis and PANTHER protein set statistical enrichment analysis performed on the panel of the 72 proteins that are upregulated in the *tibialis anterior* muscle of *Hacd1*-KO mice, compared to WT mice. *n* = 8 per group for **A-D**, *n* = 6 per group for **E-K**, and *n* = 3 per group for **L** and **M**; error bars: ± s.e.m.; \**P* < 0.05, \*\**P* < 0.01 and \*\*\**P* < 0.001.

To globally identify major molecular pathways involved in this elevated energy expenditure via an unbiased approach we performed a non-targeted large-scale proteomic analysis of *tibialis anterior* muscles from WT and *Hacd1-*KO mice. A panel of 96 proteins upregulated in *Hacd1-*KO mice was identified and further classified using the PANTHER classification system (Mi et al., 2013). The Gene Ontology biological process and molecular function involving the highest percentage of upregulated proteins were “metabolic process” (37%) and “catalytic activity” (54%), respectively (Figures 2L, S2L and Table S1). A statistical overrepresentation test confirmed that proteins involved in metabolism were globally enriched by a factor > 2 (*P* = 7.63 x 10^-10^) - most notably those of the mitochondrial tricarboxylic acid cycle, which were enriched by a factor of 37.5 (*P* = 0.0143) (Figure 2M).

### HACD1 deficiency leads to increased mass and altered function of muscle mitochondria

Altogether, these data highlighted a major, yet undescribed role of *Hacd1* in muscle metabolism and mitochondrial function. To confirm the activated catabolism of glucose and lipids is due to a mitochondrial impairment in muscles of HACD1-deficient animals, we assessed the expression of mitochondrial markers. Elevated activity of cytochrome *c* oxidase (COX) was observed in the *gastrocnemius* and *soleus* muscles of *Hacd1-*KO mice, along with an elevated activity of citrate synthase (CS) in the highly glycolytic *gastrocnemius* muscle (Figures 3A and 3B). This increase in mitochondrial mass was confirmed through the increased expression of CS, VDAC and ATP5A proteins in the *gastrocnemius* muscle (Figures S3A and S3B). Stimulation of mitochondrial biogenesis likely contributed to the observed increase in mitochondrial mass since the *Ppargc1a (PGC-1alpha)*, *Ppargc1b (PGC-1beta)*, *Tfam* and *Nrf-1* transcription factors that are major promoters of mitochondrial biogenesis were upregulated in the *gastrocnemius* muscle of *Hacd1-*KO mice along with their direct mitochondrial target *CoxI* (Figures 3C). Increased CS activity was also found in the *biceps femoris* of 5-mo-old CNM dogs (Figure S3C), *i.e.* at the onset of clinical signs of the myopathy (Tiret et al., 2003). These datasets pinpointed an increased mitochondrial mass as an early and conserved feature of HACD1 deficiency in skeletal muscles.

**Figure 3.**
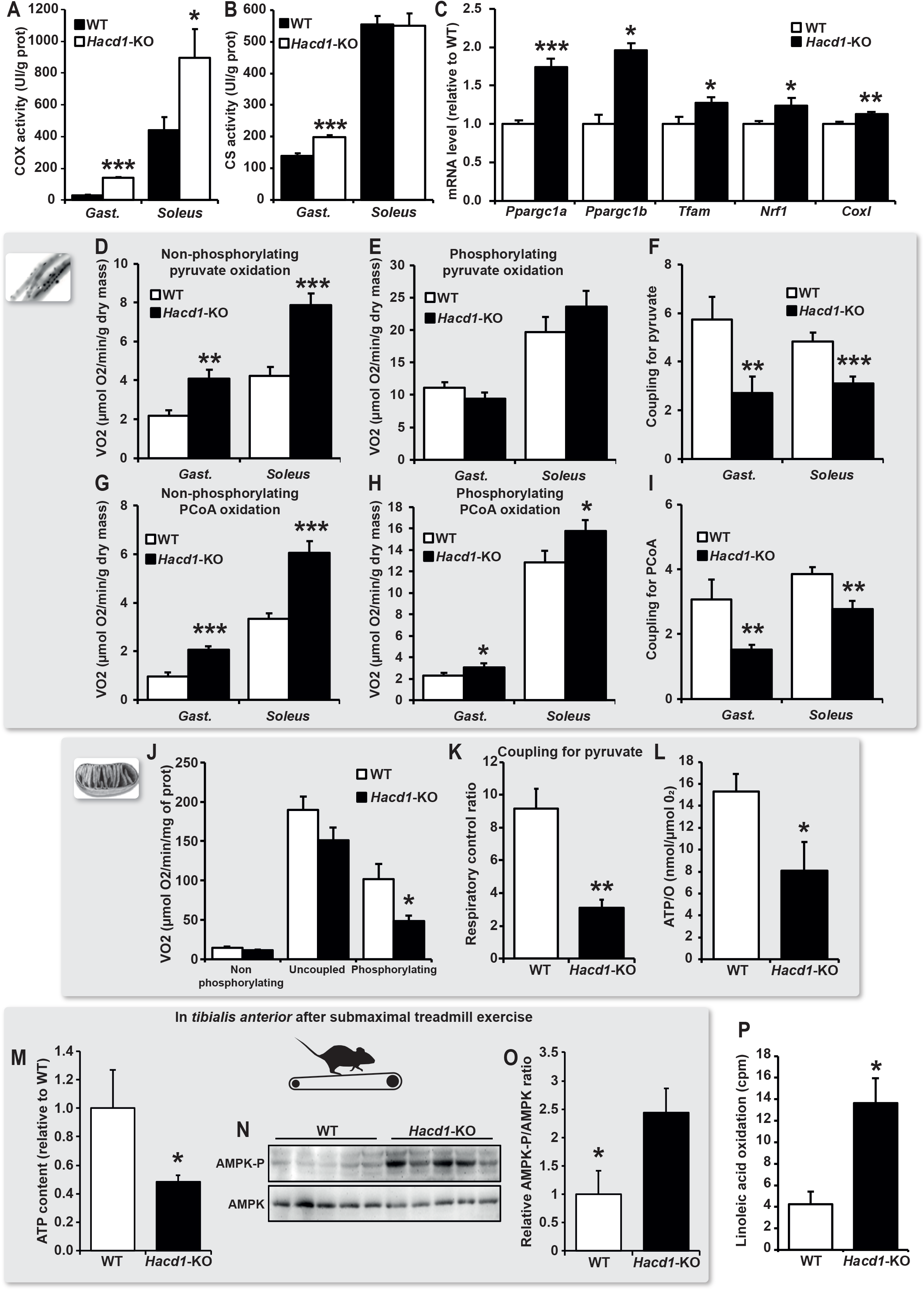
Impaired mitochondrial function in skeletal muscles of *Hacd1*-KO mice. **(A)** Cytochrome *c* oxidase (COX or Complex IV) activity in superficial *gastrocnemius* and *soleus* muscles. **(B)** Citrate synthase (CS) activity, a marker of the mitochondrial mass, in superficial *gastrocnemius* and *soleus* muscles. **(C)** *Ppargc1a, Ppargc1b, Tfam, Nrf1* and *CoxI* mRNA expression normalized by the geometrical mean of three independent reference genes in superficial *gastrocnemius* muscle. **(D** and **E)** Non-phosphorylating **(D)** and phosphorylating **(E)** oxidation rate in the presence of pyruvate in permeabilized myofibers freshly isolated from superficial *gastrocnemius* (*Gast*.) and *soleus* muscles. **(F)** Mitochondrial coupling for pyruvate (ratio of phosphorylating to non-phosphorylating oxidation rates of pyruvate from **D** and **E**). **(G** and **H)** Non-phosphorylating **(G)** and phosphorylating **(H)** oxidation rate in the presence of Palmitoyl-Coenzyme A (PCoA) in permeabilized myofibers freshly isolated from superficial *gastrocnemius* (*Gast*.) and *soleus* muscles. **(I)** Mitochondrial coupling for PCoA (ratio of phosphorylating to non-phosphorylating oxidation rates of PCoA from **G** and **H**). **(J)** Oxidation rate of freshly isolated mitochondria from *tibialis anterior* muscle (containing both glycolytic and oxidative fibers) of WT and *Hacd1*-KO mice in the presence of pyruvate plus ADP (Phosphorylating), oligomycin (Non-phosphorylating) and FCCP (Uncoupled). **(K)** Mitochondrial coupling ratio for pyruvate (Respiratory Control Ratio (RCR; State 3/State 4)) of isolated mitochondria from *tibialis anterior* muscle of WT and *Hacd1*-KO mice. **(L)** ATP/O ratio calculated from simultaneous recording of O_2_ consumption and ATP production on isolated mitochondria from *tibialis anterior* muscle of WT and *Hacd1*-KO mice. **(M)** ATP content after submaximal exercise on treadmill in *tibialis anterior* muscle of *Hacd1*-KO mice, compared to WT mice, set to 1.0. **(N-O)** Representative immunoblots **(N)** and quantification **(O)** of phospho-AMPK and AMPK in *tibialis anterior* muscle after submaximal exercise on treadmill. **(P)** ^14^C-labelled linoleic acid consumption rates in isolated *gastrocnemius* muscle. *n* = 6 per group for **A-I** and **N-O**, *n* = 4 per group for **J**-**L** and **P,** *n* = 5 per group for **M,**; error bars: ± s.e.m.; \**P* < 0.05, \*\**P* < 0.01 and \*\*\**P* < 0.001.

Function of the total mitochondrial population was then assessed in both the highly glycolytic *gastrocnemius* and the highly oxidative *soleus* muscles of *Hacd1-*KO mice, by evaluating non-phosphorylating respiration in the absence of ADP through the measurement of oxygen consumption in permeabilized muscle fibers. In both muscles, an increase in the non-phosphorylating respiration capacity was noticed in *Hacd1-*KO mice, irrespective of the substrate used (Figures 3D, 3G and Table S2) and corroborating the increased mitochondrial mass (Figure 3B). However, once normalized to the mitochondrial protein content, respiration of purified mitochondria under non-phosphorylating conditions (in the absence of ADP or uncoupled by FCCP) was normal in the presence of pyruvate (Figure 3J). This revealed that mitochondria in HACD1-deficient muscles retained normal capacities of non-phosphorylating respiration. At the same time, phosphorylating respiration of *Hacd1*-KO mitochondria in the presence of ADP was markedly lower (Figure 3J), leading to a diminished quality of respiration represented by the ratio of phosphorylating to non-phosphorylating respiration (−66%; Figure 3K). At the level of muscle fibers, which are characterized by the previously identified increase in mitochondrial mass, global phosphorylating respiration was unchanged when using pyruvate as a substrate or slightly increased when using a lipid substrate (Figures 3E and 3H), also yielding a decrease of 28 to 50% in the respiratory coupling depending on muscle and substrates (Figures 3F and 3I).

Accordingly, the ATP production flux by muscle mitochondria was reduced (Figure S3D), yielding a diminished ATP/O ratio (Figure 3L) that indicates a poorly efficient mitochondrial respiration. Remarkably, this was accompanied by neither detectable modification of the redox potential in both state 3 and state 4 (Figure S3E) nor a difference in the generation of ΔΨm in state 4 (Figure S3F) - demonstrating that successive steps of NADH production by dehydrogenases, NADH consumption by the respiratory chain and establishment of the proton gradient by the respiratory chain, respectively, were all unaffected. The consumption of ΔΨm in state 3 was also normal (Figure S3F), in contrast with models harboring a specific ATP synthase deficiency (Mayr et al., 2010). In addition, the preserved non-phosphorylating respiration indicated that no proton leak occurred (Figure 3J).

In summary, HACD1 deficiency resulted in a marked decrease of the mitochondrial oxidation-to-ATP production (OXPHOS) coupling in skeletal muscles with no modifications in proton gradient or function of the respiratory chain, and regardless of the substrate used (Figures 3F, 3I, 3K and 3L). This functional deficiency led to a significant deficit of muscle ATP content, despite increased mitochondrial mass, in *Hacd1*-KO mice challenged with an intense exercise (Figure 3M). As expected from a deficit in energy production, activating phosphorylation of the AMP kinase (AMPK) - known as an energy sensor and a promoter of mitochondrial biogenesis - was increased in this context (Figure 3N and 3O). The preference of *Hacd1*-KO mice for lipid catabolism, previously evidenced at the systemic level by respiratory exchange ratio (RER) and fatty acid oxidation measurement (Figures 2H-K), was confirmed at the tissue level by a significantly increased phosphorylating respiration of muscle fibers using two distinct lipid substrates (Figures 3H and S3G), and at the organ level by a massive increase in fatty acid β-oxidation (Figures 3P and S3H). This was associated with an upregulation of key actors of lipid oxidation and their main regulators *Ppara* and *Ppard*, concomitantly with a downregulation of *de novo* fatty acid synthesis regulators (Figures S3J). Finally, muscles of *Hacd1-*KO mice displayed a similar relative rate of oxidation of palmitoylCoA (C16) versus octanoate (C8) (Figure S3I), which, in addition to unchanged levels of transcripts coding carnitine palmitoyl transferases (*Cpt*) and fatty acid translocase (*Cd36*) (Figure S3J), supported a global increase in fatty acid catabolism rather than a privileged mitochondrial translocation of long chain fatty acids. Remarkably, the defective mitochondrial coupling induced by HACD1 deficiency was not associated with oxidative stress, as evidenced by normal ratio of aconitase/fumarase activities, expression of anti-oxidant genes and content of peroxidated lipids or carbonylated proteins (Figures S4A-S4E).

Given the expression of *Hacd1-fl* in cardiac muscle (Blondelle et al., 2015), we next evaluated whether the heart could additionally contribute to the systemic metabolic phenotype described in *Hacd1*-KO mice. In contrast to our observations in skeletal myofibers, we observed no difference in respiration capacities of HACD1-deficient cardiac myofibers (Figures S3K-M), which was in accordance with a normal cardiac function in *Hacd1-*KO mice evaluated with a comprehensive panel of echocardiographic parameters (Table S3). To evaluate whether functional HACD redundancy could specifically protect the heart from deleterious consequences of HACD1 deficiency, we quantified in WT the expression of *Hacd1* paralogs and found *Hacd2, Hacd3* and *Hacd4* more expressed in the heart, compared to the skeletal muscle, along with a relatively decreased expression of *Hacd1* (Figure S3N). In addition, a complex gene expression signature reflecting mitochondrial biogenesis, fatty acid ß-oxidation and lipogenesis was specifically identified in the skeletal muscle of *Hacd1*-KO mice, whereas their expression was unaffected in the heart, as well as in liver, white and brown adipose tissues of *Hacd1-*KO mice (Figure S3J and S3O) in which markers of mitochondrial mass were detected at normal levels (Figures S3P and S3Q). Altogether, these data strongly suggest that the systemic phenotype of increased energy expenditure and lipid catabolism of *Hacd1-*KO mice can be attributed to defective mitochondrial coupling in skeletal muscle.

### Altered domain organization and lipid composition of muscle mitochondrial membranes in HACD1-deficient mice

In accordance with a preserved mitochondrial proton gradient upon HACD1-deficiency (Figure S3G), the expression of *Ucp1*, *Ucp2* and *Ucp3,* which encode heat-producing uncoupling proteins localized within the IMM, was normal in muscles and other metabolically active organs (Figures S4F-S4I). The significant reduction in phosphorylating respiration measured in isolated mitochondria is therefore not linked to protein uncoupling or impaired function of the respiratory chain, but likely results from an intrinsic membrane defect. To investigate the putative role of membrane properties, we first assessed assembly of the ATP synthase on a native gel and observed a normal pattern in *Hacd1*-KO mice (Figure 4A). We further evaluated mitochondrial translocase activity by measuring the Km for ADP in permeabilized fibers from superficial *gastrocnemius* and *soleus* muscles, and similarly detected no difference between WT and *Hacd1*-KO mice (Figure S4J). This suggested that the defective coupling mechanism involved other, less obvious, actors presumably involved in proton translocation to the ATP synthase. Mitochondrial respiration efficiency relies on a highly specific organization and phospholipid composition of the IMM (Lewis and McElhaney, 2009; Paradies et al., 2014). We investigated properties of the IMM using TMA-DPH, this fluorescent probe is captured by and stored in the membrane allowing dynamic assessment of the organization of surrounding lipids by polarized excitation (Klausner et al., 1980; Shrivastava et al., 2016; Stubbs et al., 1995). In particular, the TMA-DPH fluorescence lifetime decay, which more specifically reflects membrane molecular organization, differed significantly for the IMM of *Hacd1*-KO mice when compared to WT. The most prominent difference was found for the longest lifetime components that correspond to the most polar domains (Figure 4B, Table S4 and Listing S1) (Karnovsky et al., 1982; Stubbs et al., 1995). By contrast, no change was observed for the outer mitochondrial membrane (Figure 4C and Table S4). These results revealed that *Hacd1* contributes to establishing a functional molecular organization of the IMM, likely including polar lipids. HACD1 is involved in the synthesis of VLCFA, hence we quantified the full panel of individual fatty acids contained in muscle mitochondria phospholipids (Figure S5A). The smallest fatty acids produced by the VLCFA elongation complex are C18 species (Ohno et al., 2010; Sawai et al., 2017). Accordingly, the amount of C18-C26 fatty acids was significantly reduced in mitochondria isolated from *Hacd1*-KO skeletal muscles with a corresponding increase in shorter C10-C17 fatty acids (Figures S5B), yielding a significant reduction in C18-C26 to C10-C17 ratio (Figure 4D). Contents in n-3 and n-6 polyunsaturated fatty acids were also decreased in *Hacd1-*KO mice (Figures S5C-S5F). In parallel, a global comparative analysis of phospholipids revealed that mitochondria isolated from *Hacd1-*KO muscles contained only 54% of the total phospholipids quantified in WT mitochondrial membranes when normalized to mitochondrial protein content (Figure 4E). This massive reduction in the phospholipid to protein ratio resulted from a combined reduction in the absolute content of sphingomyelin (SM), phosphatidylethanolamine (PE), lysophosphatidylethanolamine (LPE), phosphatidylserine (PS), phosphatidylinositol (PI) and cardiolipin (CL) species (Figure 4F), the two latter being the most severely reduced (PI, - 45%; CL, - 49%). Relative to the total phospholipid content, the two anionic CL and PI were also reduced, in association with an increase in the neutral phosphatidylcholine (PC) (Figure 4G). Of note, acyl chain composition of CL species was unchanged in *Hacd1-*KO mice (Figure S6), showing that the Tafazzin-dependent step of CL remodeling was preserved in this context. In contrast, and in accordance with the absence of a deleterious cardiac phenotype in *Hacd1-*KO mice (Table S3), phospholipid composition of mitochondria isolated from the cardiac muscle was unaltered in *Hacd1-*KO mice (Figure S5G and S5H). Altogether, these data strongly suggested that the altered lipid composition of skeletal muscle mitochondria in HACD1-deficient mice was the primary cause of their dysfunction and that, in addition to modifying fatty acid chain length, HACD1 deficiency yielded a more complex, muscle specific alteration in mitochondrial membrane phospholipid composition.

**Figure 4.**
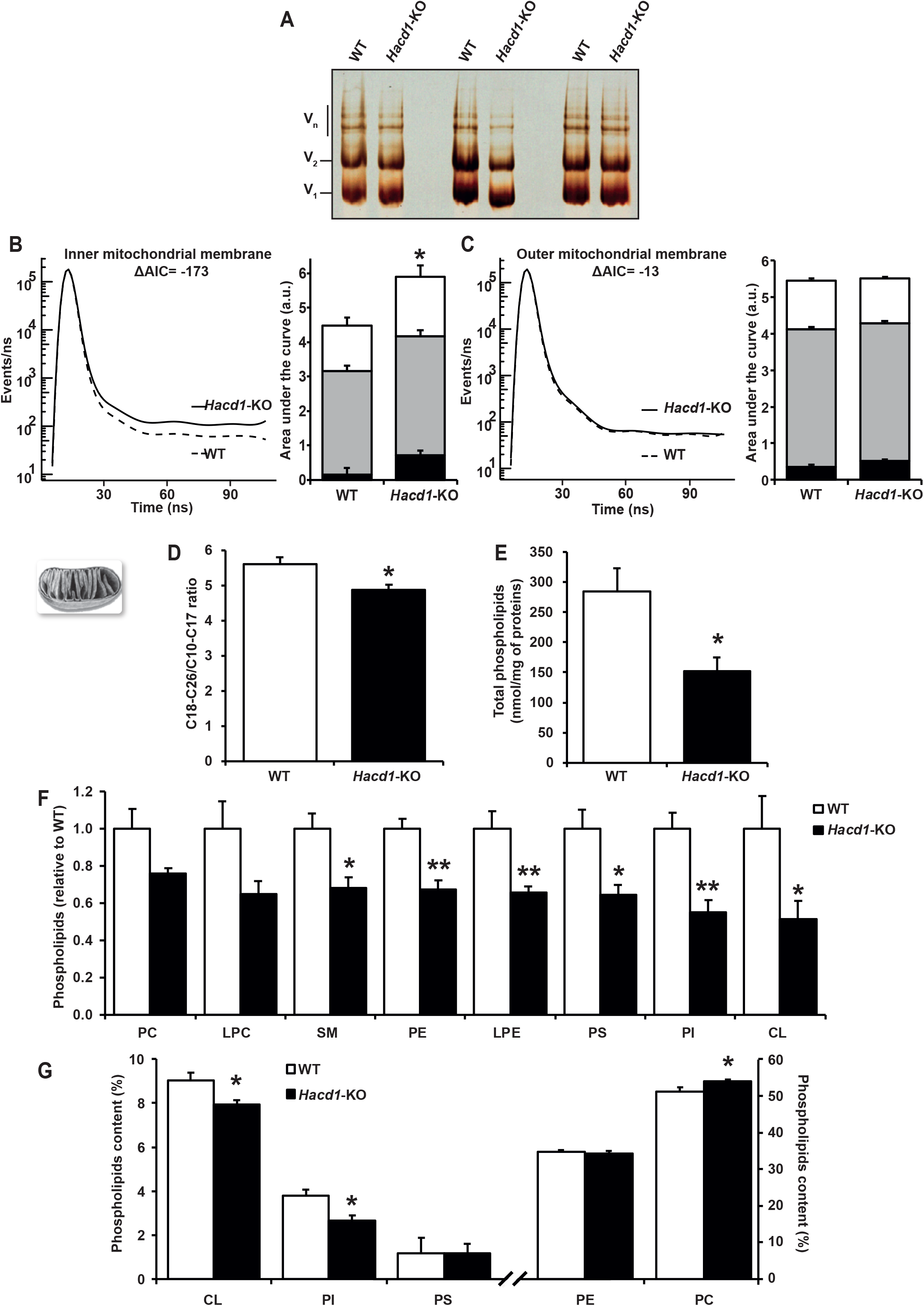
Impaired lipid composition of mitochondrial membranes in skeletal muscles of *Hacd1*-KO mice. **(A)** In-gel activity of mitochondrial ATP synthase complexes, isolated from the *tibialis anterior* muscle. **(B** and **C)** TMA-DPH lifetime decay (curves) and mixing proportions (histograms) in inner **(B)** and outer **(C)** mitochondrial membranes, measured on mitoplasts and intact isolated mitochondria from the *tibialis anterior* muscle, respectively. **(D)** Ratio of C18-26 to C10-17 total phospholipid fatty acids of mitochondria isolated from the *tibialis anterior* muscle. **(E)** Total phospholipid content of mitochondria isolated from the *tibialis anterior* muscle, normalized to the mitochondrial protein content. **(F)** Phospholipid species content of mitochondria isolated from the *tibialis anterior* muscle of *Hacd1*-KO mice, normalized to the mitochondrial protein content and compared to those of WT mice, set to 1.0. **(G)** Relative phospholipid species content of mitochondria isolated from the *tibialis anterior* muscle, normalized to the total content of mitochondrial phospholipids. PC = phosphatidylcholine, LPC = lysophosphatidylcholine, SM = sphingomyeline, PE = phosphatidylethanolamine, LPE = lysophosphatidylethanolamine, PS = phosphatidylserine, PI = phosphatidylinositol and CL = cardiolipins; *n* = 4 per group for **A-C,** *n* = 5 per group for **D-G**; ΔAIC = difference in the Akaike’s Information Criterion (full analysis in Tables **S4**); error bars: ± s.e.m.; \**P* < 0.05 and \*\**P* < 0.01.

### Altered cristae and cardiolipin-dependent OXPHOS efficiency

CL are enriched in the IMM and are essential to cristae organization (Paradies et al., 2014). The decrease in CL content associated with altered physical properties of the IMM upon HACD1-deficiency prompted us to further investigate the ultrastructural organization of *Hacd1*-KO muscle mitochondria. We first validated the quality of our samples by observing mitochondria with a preserved ultrastructure in muscles of WT mice, including quantifying size variation and shape of normal cristae (Figures 5A-C and S7A-C). In *Hacd1-*KO mice, we noticed exaggerated dilation of cristae tips (> 15 nm) in 23 and 30% of mitochondria from *gastrocnemius* and *soleus* muscles, compared to 0.7 and 1.3 % in WT, respectively (Figures 5A, 5B, S7A and S7B), and occasionally observed some mitochondria including both normal and dilated cristae. Precise quantification of cristae ultrastructure confirmed their significant thickening (Figures 5C and S7C).

**Figure 5.**
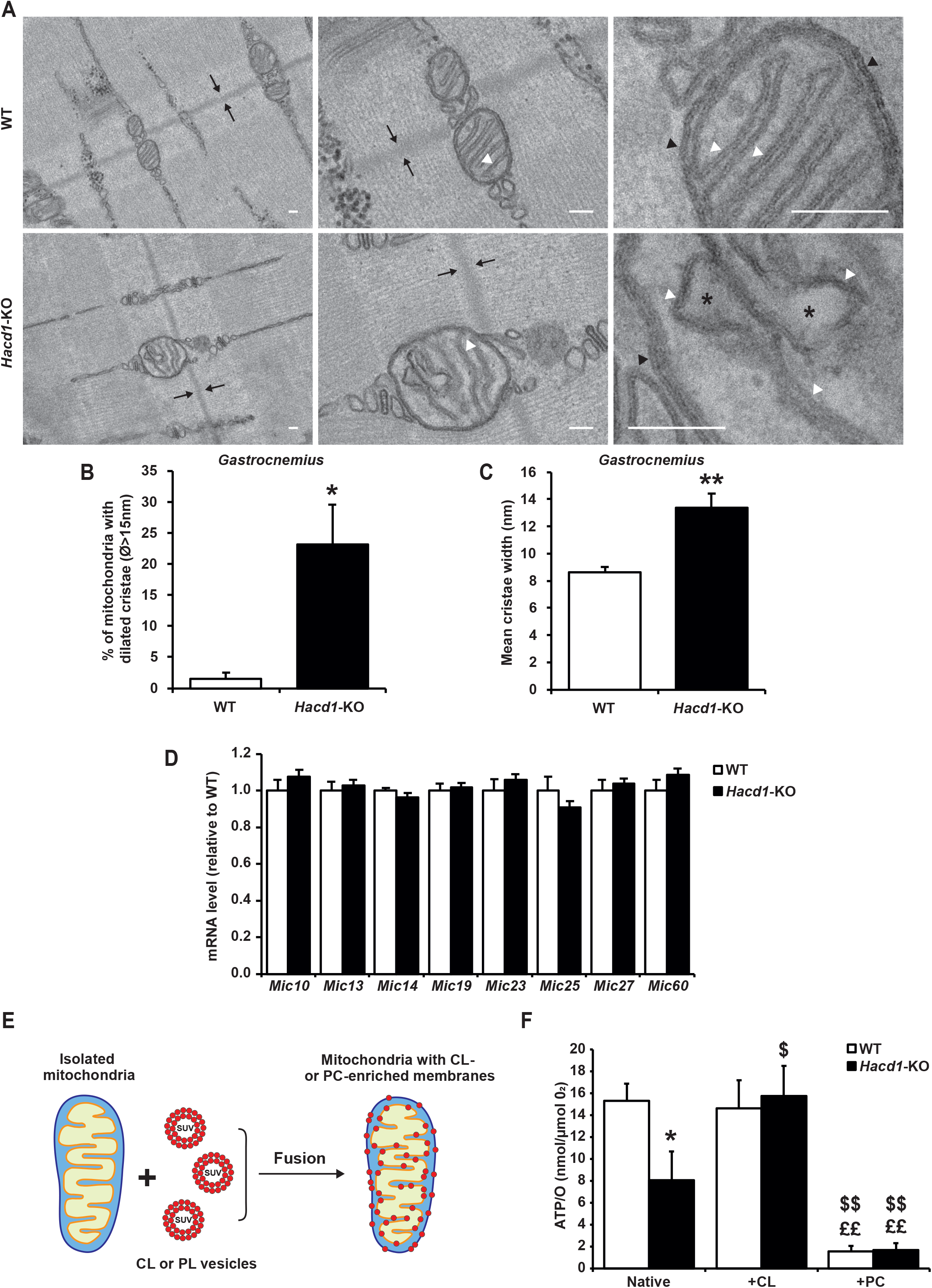
Abnormal cristae in mitochondria of *Hacd1*-KO mice muscle and functional rescue of coupling by cardiolipin. **(A)** Transmission electron microscopy of longitudinal sections of myofibers from the superficial *gastrocnemius* muscle of wild-type (WT; upper panel) and *Hacd1*-KO (lower panel) mice. Images are from datasets taken at a low (x 2,500; left panel), intermediate (x 10,000; middle panel) and high (x 30,000; right panel) magnification. The Z-line is delimited by arrows. Cristae are regular tubular-shaped invaginations (white arrow heads) of the inner mitochondrial membrane, unambiguously identified on the same plane than the outer mitochondrial membrane (black arrow heads). Excessive dilation of cristae tips (asterisk) is frequently observed in *Hacd1*-KO myofibers. **(B** and **C)** Morphometric quantification of mitochondria containing cristae with excessively dilated tips (maximal diameter >15nm, in percentage) in the superficial *gastrocnemius* muscle **(B)** and the mean of the maximal diameter of cristae **(C)**. **(D)** mRNA expression of major genes encoding the MICOS complex, normalized by the geometrical mean of three independent reference genes, in the superficial *gastrocnemius* muscle. (**E**) Diagram depicting the experimental steps allowing to enrich mitochondrial membranes with tagged phospholipids. (**F**) ATP/O ratio calculated from simultaneous recording of O_2_ consumption and ATP production of native, cardiolipin-enriched (+CL) and phosphatidylcholine-enriched (+PC) mitochondria isolated from *tibialis anterior* muscle of WT and *Hacd1*-KO mice. Scale bars in **A**: 100 nm; *n* = 4 per group for **A-C**, *n* = 8 per group for **D,** *n* = 4 per group for **E-F**; error bars: ± s.e.m.; \**P* < 0.05 and \*\**P* < 0.01 vs WT, $ = *P* ≤ 0.05 vs Native, $$ = *P* ≤ 0.01 vs Native, ££ = *P* ≤ 0.01 vs +CL.

In addition to CL, two major protein complexes contribute to organizing the functional IMM architecture: dimers of ATP synthase keep tight membrane curvature along the cristae and the MICOS complex is involved in shaping cristae at their junction with the inner boundary membrane (Habersetzer et al., 2013). A defect in ATP synthase oligomerization was excluded (Figure 4A) and in parallel, we found unaltered contents of mRNA transcribed from genes encoding the MICOS complex (Figure 5D). Similarly, we observed no visible modification in the assembly of respiratory supercomplexes (data not shown).

Taken together these results demonstrated major ultrastructural cristae modifications upon *Hacd1* deficiency; we excluded alterations in ATP synthase and MICOS oligomers, therefore this is more likely attributable to intrinsic modifications of lipid composition of the IMM.

To confirm the *Hacd1*-dependent regulatory role of membrane lipid composition (including CL) on muscle mitochondrial coupling, we designed an *in vitro* assay to enrich mitochondrial membranes with specific phospholipids taking advantage of their capacity to fuse with small unilamellar vesicle composed of CL or PC (CL/PC-SUV) (Figure 5E). We validated phospholipid incorporation within the IMM using fluorescent CL and PC lipids (Figure S7D). CL enrichment was also quantified using a Acridine Orange 10-Nonyl Bromide (NAO) probe and we detected an increase of 43% and 33% of NAO signal in both mitochondria and mitoplasts respectively (Figures S7E and S7F).

Then, we simultaneously measured O_2_ consumption and ATP production of mitochondria following specific PL enrichment. A decrease in O_2_ consumption and ATP production was observed with both CL- and PC-SUV in the two WT and *Hacd1-* KO conditions (Figure S7G-H), likely because PL enrichment non-selectively dilutes respiratory chain complexes within the membrane (Schneider et al., 1980). However, the ATP/O ratio that reflects coupling efficiency of ATP synthase activity with the respiratory chain function was fully rescued when muscle mitochondria from *Hacd1*-KO mice were specifically enriched with CL, whereas it was dramatically decreased by PC in both *Hacd1*-KO and WT mice (Figure 5F). The functional rescue triggered by CL indicated a key role of *Hacd1* in the precise regulation of IMM lipid composition, in particular CL content, crucial for respiratory coupling efficiency in muscle.

### HACD1-deficient mice are protected against HFD-induced obesity

*Hacd1*-deficient mice had disorganized and less efficient muscle mitochondria, yielding a lower ATP production and hence compensatory mechanisms such as increased mitochondrial mass and substrate consumption – in particular fatty acids. To evaluate whether energy of the diet would have been a limiting factor in the efficiency of these adaptive mechanisms, we challenged *Hacd1*-KO mice by feeding them with HFD for nine weeks. As previously noticed when fed with a ND, HFD-fed *Hacd1-*KO mice had an increased cumulative food intake when normalized to their body mass (Figures 6A, S8A and S8B). However, from the onset of the HFD we observed a regular, striking reduction of their relative body mass gain by more than half over this period compared to WT (Figures 6B and 6C). Fecal lipid loss was unchanged in *Hacd1*-KO mice (Figure S8C), indicating that the massive decrease in feed efficiency (−66%, Figure S8D) was due to enhanced catabolism. As in ND conditions, HFD-fed *Hacd1-*KO mice had no detectable elevation of their body temperature (Figure S8E). The BAT as well as the gonadal, retroperitoneal and mesenteric white fat pads of *Hacd1-*KO mice were reduced (Figures 6D-6F, S8F, S8G and Table S5), leading to a global reduction of 43% in their adiposity index compared to WT (Figure 6G). At the cellular level, we observed a significant reduction in the size of adipocytes (−46% of the surface area, Figure 6H). In parallel, *Hacd1*-KO mice also displayed resistance to liver and muscle steatosis (Figures 6I and 6J). At the biochemical level, serum concentrations in cholesterol and free fatty acids were similar in *Hacd1-*KO and WT mice whereas triglyceridemia was reduced in *Hacd1*-KO mice (Figures S8H-J). Finally, after nine weeks of HFD, *Hacd1-*KO challenged to a glucose overload retained their higher glucose tolerance (Figures 6K and 6L) and sensitivity to insulin (Figure S8K), as previously noticed under a ND. Altogether, these results highlighted a striking mechanism whereby increased energy expenditure and lipid catabolism of *Hacd1-*KO mice due to defective mitochondrial coupling in skeletal muscle protected against obesity.

**Figure 6.**
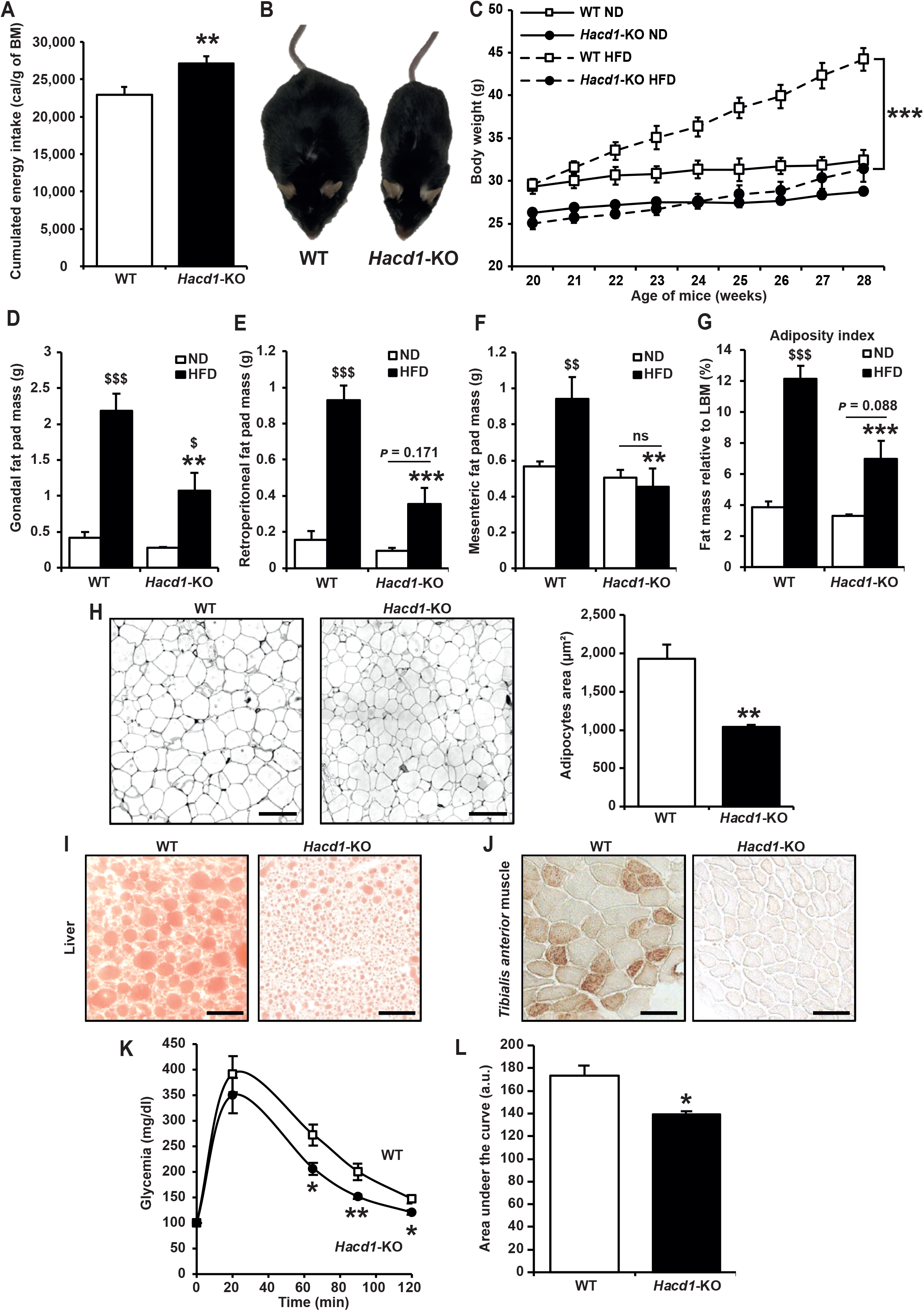
*Hacd1* deficiency results in protection against high fat diet-induced obesity and insulin resistance. **(A)** Cumulated energy intake during the 9-wk period of high-fat diet (HFD). **(B)** Morphology of wild-type (WT) and *Hacd1*-KO mice after 9 wk of HFD. **(C)** Body mass weekly evolution of mice fed during 9 wk with HFD, compared to age-matched mice fed with normal diet (ND; also presented in Figure 2A). **(D-F)** Mass of gonadal **(D)**, retroperitoneal **(E)** and mesenteric **(F)** fat pads after 9 wk of HFD. **(G)** Total body fat percentage (adiposity index) after 9 wk of ND or HFD, expressed as a percentage of the eviscerated body mass (lean body mass, LBM). **(H)** Hematoxylin&Eosin-stained transverse sections of gonadal fat pads after 9 wk of HFD, highlighting the contour of adipocytes, and morphometric quantification of adipocytes surface. **(I** and **J)** Oil-Red-O-stained transverse sections of liver **(I)** and *tibialis anterior* muscle **(J)** after 9 wk of HFD. **(K)** Glycemia measured after fasting and assessed during 120 min after an intraperitoneal glucose injection at T_0_ in mice fed during 9 wk with HFD. **(L)** Area under the glycemia curves displayed in **(K)**. Scale bars represent 50 µm in **H**, **I** and **J**; *n* = 8 per group for **A-G**, *n* = 4 per group for **H-J**, *n* = 7 per group for **K** and **L**; error bars, ± s.e.m.; \**P* < 0.05, \*\**P* < 0.01 and \*\*\**P* < 0.001 *versus* respective WT values. $*P* < 0.05 and $$$*P* < 0.001 *versus* respective ND values.

## DISCUSSION

Focusing on the essential metabolic role of skeletal muscles in maintaining homeostasis we conducted in depth analyses of mice and dogs deficient for HACD1, which is highly expressed in striated muscles. We have unraveled a novel, skeletal muscle-specific mechanism whereby a pathway regulating the lipid composition of mitochondrial membranes impacts cristae shape, finely-tuned capacities of coupled oxidative phosphorylation and, consequently, adaptive optimization of systemic energy expenditure to cope with the high ATP demand in muscles.

Despite accumulated data supporting a crucial role of VLCFA in cell homeostasis of organs such as the skin, eye, liver or testis (Bhandari et al., 2016; Matsuzaka et al., 2007; Sassa et al., 2013), little is known regarding their specific role in muscles, or about mechanisms whereby cells control VLCFA contents (Ikeda et al., 2008; Kobayashi and Nagiec, 2003; Zimmermann et al., 2017). Spontaneous or induced animal models of HACD1 deficiency have revealed unanticipated effects. Contrarily to our initial hypotheses, reduction of locomotor activity and muscle mass in *Hacd1*-KO mice was associated with increased glucose tolerance and insulin sensitivity, and with an unexpected raise in energy expenditure at the whole-body level. We identified the source of this energy leak in an insufficient phosphorylating respiration in skeletal muscles. Consequently, the resulting low-energy status triggers activation of the AMPK sensor, which likely activates compensatory mechanisms of increased mitochondrial biogenesis and fatty acid oxidation, yet is insufficient to overcome the energy deficit. Altogether, these findings highlight novel roles of HACD1 and VLCFA in regulating muscle and systemic metabolism to meet skeletal muscle energy needs.

The molecular mechanism leading to lowered phosphorylating capacity of mitochondria from HACD1-deficient muscles does not rely on classical uncoupling mechanisms leading to proton leak; rather, it involves a prominent alteration in the lipid composition of mitochondrial membranes with subsequent modifications of their physical properties, including a remodeled organization of the IMM. More precisely, mitochondrial phospholipid content dropped dramatically in *Hacd1*-KO mice and was associated with a decrease in PI and CL relative to the other phospholipids. Remarkably CL is a four acylchain phospholipid nearly restricted to the IMM where its specific accumulation is proposed to favor strong curvatures allowing tubulation of cristae (Paradies et al., 2014). Using CL-lacking models, CL has also been implicated in assembly of respiratory supercomplexes and ATP synthase dimers (Acehan et al., 2011; Zhang et al., 2005), two processes that were not altered in *Hacd1*-KO mice. Notwithstanding the confirmed prominent role of CL in shaping strong curvature of cristae, our data indicate that assembly of molecular complexes allowing ATP production can be achieved even with a reduced content of CL. This unveils the existence of distinct CL content thresholds to achieve complementary roles of CL in ATP production. The two last steps of CL synthesis occur within mitochondria, first CL synthase catalyses binding of CDP-diacylglycerol to phosphatidylglycerol to form CL, and then Tafazzin remodels the acylchain composition in the heart and skeletal muscles, which results in an enrichment of linoleoyl chains (C18:2, ω-6) in CL (Oemer et al., 2018). The functional importance of the latter reaction is illustrated by Barth syndrome, characterized by a life-threatening infantile dilated cardiomyopathy, which develops associated with Tafazzin deficiency (Barth et al., 1983; Ikon and Ryan, 2017a). Notably, acylchain repartition among CL species was unchanged in HACD1-deficient muscle mitochondria, indicating that the last Tafazzin-dependent step of CL maturation is unaffected. Of note, alteration in lipid composition of mitochondrial membranes was specific to skeletal muscles, and in particular phospholipid composition of cardiac mitochondria was unchanged in *Hacd1*-KO mice, in agreement with a normal cardiac function. In these mice, HACD activity in the heart is presumably achieved by the redundant paralogous *Hacd* genes, highly expressed in the heart (Figure S3P; (Ikeda et al., 2008; Wang et al., 2004). In contrast with the experimentally-induced respiratory uncoupling through proton leak (Jastroch et al., 2010), the defective mitochondrial coupling in HACD1-deficient skeletal muscles produced no oxidative stress at the cellular level. This is consistent with our longitudinal study over 20 years demonstrating that HACD1-deficient dogs under medical care have a normal life span, similarly to *Hacd1*-KO mice, and indicates that human patients could derive benefit from innovative treatments that manipulate this pathway.

The mechanism whereby a specific composition and structure of the IMM dynamically optimizes respiratory coupling is still a subject of active investigation. Careful analysis of the respiratory parameters observed in *Hacd1*-KO mice revealed normal activity of the respiratory chain (*i.e.* of complexes I to IV). We also measured normal V_0_ (*i.e.* O_2_ consumption without ADP), which indicates normal, basal back leakage of protons through other IMM ion channels than the ATP synthase. To emphasize that the reduced coupling efficiency observed here results from a mechanism independent of proton leak that yields classical uncoupling, and to fit with international guidelines (https://tinyurl.com/mitopedia), we referred to our finding as a mitochondrial dyscoupling. Upon HACD1-deficiency, molecularly modified membranes hamper efficient coupling between proton pumping and ATP synthesis by the ATP synthase, which we found normally oligomerized. A specific movement of protons along the polar heads of phospholipids, which is faster than in the aqueous phase, has been demonstrated (Prats et al., 1986; Teissié et al., 1985). In a comparative approach including archae, bacteria, plants and animals, it has been proposed as a general model that once protons exit a proton pump, a von Grotthuß mechanism mediated by anionic lipid headgroups might transfer them from their entry portals on another species of proton pump, providing a membrane surface proton transport circuitry that has been named the “hyducton” (Yoshinaga et al., 2016). Due to its anionic charge and abundance, CL is proposed as a highly efficient proton trap within the IMM, supplying at high speed protons to the ATP synthase and allowing it to reach maximal efficiency before any major change in pH occurs (Haines and Dencher, 2002). By documenting an absolute 2-fold drop in PL and a relative decrease in the two anionic CL and PI, combined with a low respiratory efficiency despite normal activity of the respiratory chain and normal assembly of ATP synthase, our *in vivo* data provide a strong physiological support to the hyducton model. Our *in vitro* data further demonstrated that providing additional CL to the IMM fully rescued the dyscoupling phenotype of muscle mitochondria from *Hacd1*-KO mice. Together these results provide compelling evidence that in addition to their role on cristae formation, CL plays a direct, crucial molecular role on mitochondrial coupling (Figure 7A).

**Figure 7.**
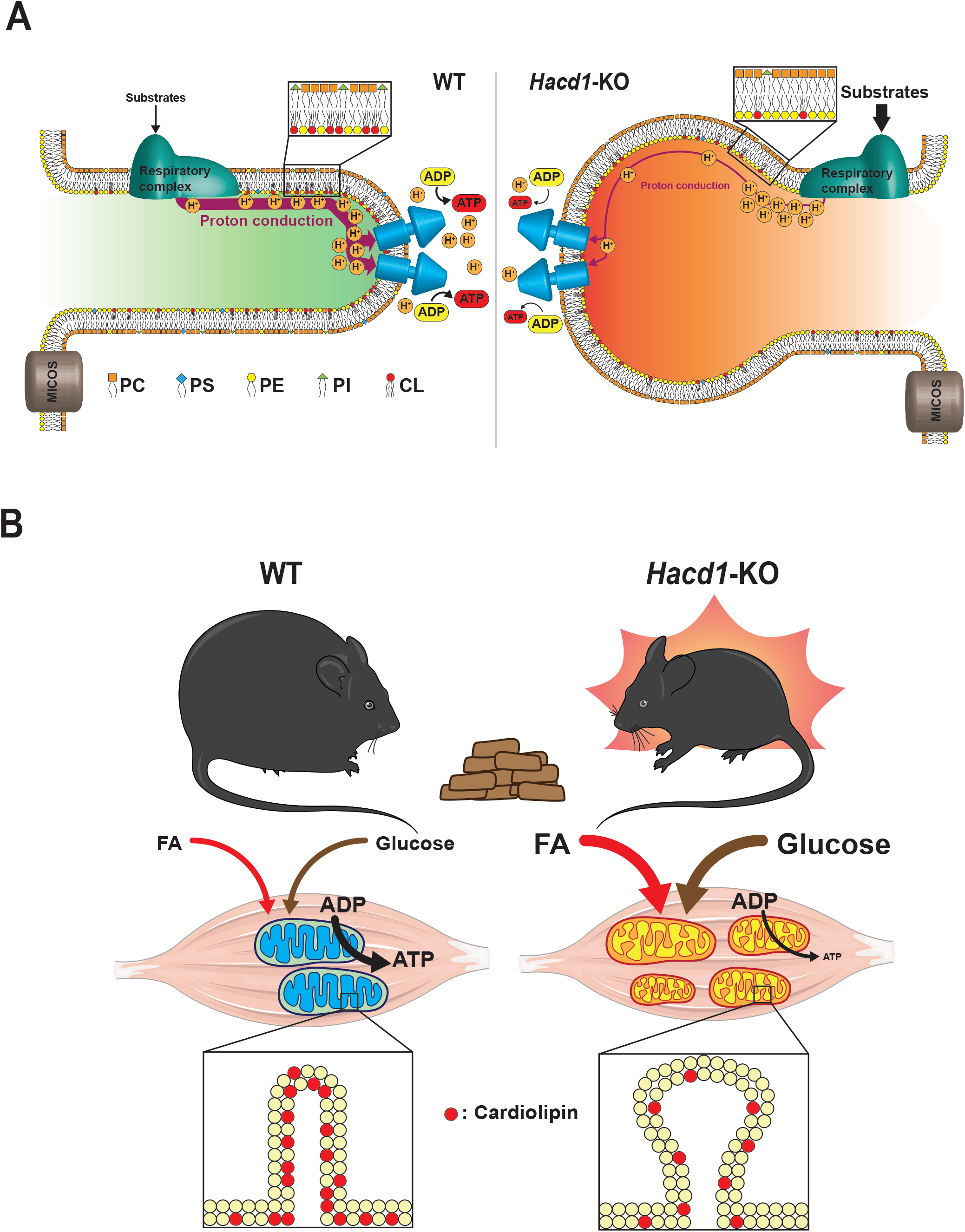
Graphical summary of ambivalent consequences of HACD1 deficiency. (**A**) Model of the molecular mechanism underlying the mitochondrial dyscoupling. In WT conditions, anionic lipids such as cardiolipin included in the inner leaflet of the inner mitochondrial membrane contribute to OXPHOS efficiency by promoting the transfer of protons from the respiratory chain to ATP synthase oligomers. In *Hacd1*-KO mice, the decreased content of anionic lipids is deleterious to the proton flux, hence impairing ATP production. (**B**) The muscle-specific mitochondrial dyscoupling resulting from HACD1 deficiency increases energy expenditure in mice fed with a high fat diet, protecting them from obesity and collateral molecular damages or vital functions that have so far been identified as limiting factors in the proposed treatments.

Importance of mitochondrial membranes lipid composition *per se* in the functional structure of cristae is exemplified in HACD1-deficient muscles by unaltered levels of all the presently known proteins acting in cristae shaping and function, such as the ATP synthase, respiratory supercomplexes and the Micos complex. CL is known to play an essential role in cristae biogenesis (Ikon and Ryan, 2017b; Khalifat et al., 2008) and through cristae remodeling, may also trigger the dynamic adaptation of mitochondria, with reversible modifications in cristae structure observed to meet variable metabolic needs of the cell. Indeed low ADP conditions prompt cristae to organize into a so called orthodox structure, whilst upon high ADP availability, they rather organize into a condensed structure, characterized by a small amount of matrix and abundant cristae (Hackenbrock, 1966) that support optimal ATP production (Lizana et al., 2008; Mannella, 2006; Song et al., 2013). HACD1-deficient mitochondria should facilitate further investigation targeted towards deciphering causal relationships linking cristae shaping and respiratory efficiency.

Human societies are facing an unprecedented epidemic of obesity, which is accompanied by hepatic steatosis, accumulated abdominal fat and reduced insulin sensitivity, altogether leading to a clinical entity called the metabolic syndrome (Eckel et al., 2005). Prevention or treatment of obesity-related disorders include low energy diet and regular exercise, yet the required level of energy restriction is hard to cope with in the long-term, exercise compliance is extremely low in obese people and often the beneficial effects are cancelled by the fact that appetite is unfortunately stimulated by exercise (Kraschnewski et al., 2010; Westerterp, 2010). In the context of this vicious circle, an appealing strategy to combat obesity and restore insulin sensitivity has been to associate an increase in lipid oxidation with a controlled reduction of cellular energy efficiency by favoring mitochondrial uncoupling (Boström et al., 2012; Colman, 2007; Tseng et al., 2010). Dinitrophenol (DNP), a potent mitochondrial uncoupling compound, was successfully used during three decades for weight loss, and then withdrawn in 1938 because of its toxicity mostly in highly ATP-dependent tissues including the heart (Grundlingh et al., 2011). Among promising strategies, is a controlled-release formulation for DNP that specifically targets the liver (Perry et al., 2015). Alternatively, proteins such as UCP1, a physiological uncoupler in BAT, represent interesting targets to increase energy dissipation (Kim and Plutzky, 2016). Here we provide evidence that the skeletal muscle-specific genetic modulation of mitochondrial membrane composition is efficient in protecting mice against a diet-induced obesity, with an improved insulin sensitivity, an absence of oxidative stress and a fully preserved cardiac function (Figure 7B). Importantly, these benefits occur in mice displaying a reduction in their skeletal muscle mass associated with a two-fold reduction in their locomotor activity, a situation mimicking the deleterious clinical condition of most obese patients. Our data thus represent an *in vivo* proof of concept that promoting dyscoupling in skeletal muscle mitochondria can confer a protection against obesity, even in patients with reduced muscle mass and disabled from exercising.

Together, these findings reveal that *Hacd1* and VLCFA exert coupling control in skeletal muscle to promote a high phosphorylating respiration, in particular through specific content and properties of mitochondrial anionic phospholipids, which likely contribute to optimizing proton bioenergetics.

## SUPPLEMENTAL INFORMATION

Supplemental Information including eight figures, six tables and one listing can be found with this article online.

## AUTHOR CONTRIBUTIONS

Conceptualization, A.P., J.B. and F.P-S; Methodology, A.P., S.G., A.S., N.Ko., G.D., M-F.G., S.B., S.L. and F.P-S; Formal Analysis, A.P., J.B., A.V., J.P., R.D., S.G., A.M., M.L., M.M., C.H., N.Ka., G.C., N.B-G., L.G., I.B., M.G., A.S., J.C., J.T., J.D., A.M., G.D. and F.P-S; Data Curation, A.P., J.B., J.P., R.D., S.G., H.C., A.M., C.H., A.S., F.D., J.D., A.M., N.Ko., G.D., M-F.G., V.V., S.L. and F.P-S; Figures Conception, A.P., J.B., R.D., H.C., A.M., L.T. and F.P-S; Writing – Original Draft, A.P., L.T. and F.P-S; Writing – Review & Editing, all authors; Funding Acquisition, S.B., F.R., L.T. and F.P-S; Resources, J.P., R.D., S.G., H.C., A.M., I.B., J.D., A.M., N.Ko., M-F.G., S.B., F.J., V.V., S.L., F.R., L.T. and F.P-S; Supervision, A.P., F.R., L.T. and F.P-S.

## ACKNOWLEDGMENTS

The authors are grateful to Camille Laisne, Diana Gelperowic and the UETM team for taking care of mice and dogs; the Knock-Out Mouse Program (the “KOMP”) for initially providing the ES cell lines carrying the knocked out allele of *Hacd1*; Jean-Paul Pais de Barros and the LAP platform (Dijon, France) for lipidomic analyses; Garance Delahaye, Marie-Perle Sellier and Camille Poncet for technical help; Renée Ventura-Clapier for insightful discussions; Gemma Walmsley for correcting the manuscript. We acknowledge the technical platform of metabolism of the Unit “Biologie Fonctionnelle et Adaptative” (University Paris Diderot, Sorbonne Paris Cité, BFA, UMR 8251 CNRS, Paris, France) for metabolic analysis and the animal core facility “Buffon” of the Université Paris Diderot Paris 7/Institut Jacques Monod, Paris for animal husbandry. We acknowledge Sylvère Durand and the technical platform of metabolomics of the Institut Gustave Roussy (LS2M plateform, IGR, Villejuif, France) for metabolic analysis. N.Ka.’s PhD fellowship is financially supported by the French Fondation pour la Recherche Médicale (FRM). The work was primarily supported by the Agence Nationale de la Recherche (ANR-12-JSV1-0005), the Association Française contre les Myopathies (AFM 16143 and Translamuscle), the CNM Project and the French Institut national de la santé et de la recherche médicale (Inserm). The authors declare that they have no conflict of interest.

## SUPPLEMENTAL TABLE LEGENDS

**Supplemental Table S1 | Proteomic dataset and Panther analysis.** Related to Figure 2.

**Supplemental Listing S1 | Final non-linear mixed effects models of the fluorescence decay of TMA-DPH on mitoplasts and whole mitochondria**. Related to Figure 4.

Specifics, given in the *R statistical language* for the **nlme** library, of the non-linear mixed effects models A, B1, B2 and C. More details can be found at <u>github.com/hchauvin/prola2019</u> (DOI: 10.5281/zenodo.1228112).

**Supplemental Table S2 | Mitochondrial respiration rates.** Related to Figure 3.

Analyses were performed in muscle fibers from 4-mo-old male mice and in cardiac fibers from 8-mo-old male mice. Complex activity and substrate utilization unit is µmolO_2_/min/g of dry mass of fibers or per mg of protein for isolated mitochondria. Respiration rates in the presence of saturating pyruvate and ADP (Phosphorylating, State 3) and after complete ADP consumption (State 4). RCR: Respiratory Control Ratio. Respiration rates in the presence of oligomycin (Non-phosphorylating) and FCCP (Uncoupled). At the end of the experiment, cytochrome *c* was added to the medium (Cyt-C stimulation) to check that the stimulation of respiration did not excess 10%, confirming the integrity of the outer mitochondrial membrane. Simultaneous measurement of respiration rates and ATP production in the presence of saturating pyruvate and ADP (State 3) for ATP/O determination. Values are given as mean ± s.e.m.. Number in parentheses is the number of analyzed animals. Value for each animal was the mean of 1-3 independent analyses. For statistical analysis, unpaired t-test was used. ACR: Acceptor Control Ratio.

**Supplemental Table S3 | Echocardiographic parameters of WT and *Hacd1*-KO mice**. Related to Figure S3.

Cardiac function of 7-mo-old mice fed with normal diet was analyzed by echocardiography. EF: Ejection Fraction; FS: Fractional shortening; HR: Heart rate; LVd, LVs: Left Ventricle diameter in diastole and systole, respectively; IVSd, IVSs: interventricular septum thickness in diastole and systole, respectively; PWd, PWs: posterior wall thickness in diastole and systole, respectively. Strain rate was measured in unit/s (ant.) and (post.). Results are represented as mean ± s.e.m.. *n* = 8 per group.

**Supplemental Table S4 | Non-linear mixed-effects models of the fluorescence decay of TMA-DPH**. Related to Figure 4.

Values are given as fixed effects ± s.e for the variables defined in models A, B1, B2 and C (see *Materials and Methods*). More specifically, *B*_1_, *B*_2_ and *B*_3_ are scale coefficients for the three exponentials and *θ*_1_, *θ*_1_ and *θ*_1_ are monotonous parametrizations of characteristic times. For models B1, B2 and C, the variables are regressed against the mice phenotype (WT or *Hacd1*-KO). The contrast matrix for this regression is given in the *Materials and Methods*. *P* < 0.1; \**P* < 0.05; \*\**P* < 0.01; \*\*\**P* < 0.001.

**Supplemental Table S5 | Anatomical parameters of WT and *Hacd1*-KO mice after normal or high fat diet**. Related to Figure 6.

Anatomical parameters measured on 7-month-old mice after 9 wk of normal (ND) or high fat diet (HFD). BM: body mass; initial and final: before and after the 9-wk period of ND or HFD, respectively; BM gain: difference between final and initial body mass; HM: heart mass; TA: *tibialis anterior* mass; GWAT, RWAT and MWAT: gonadal, retroperitoneal and mesenteric white adipose tissue mass, respectively; BAT: brown adipose tissue mass. Results are represented as mean ± s.e.m.. *n* = 9. \**P* < 0.05; \*\**P* < 0.01; \*\*\**P* < 0.001 *versus* respective WT values. $*P* < 0.05; $$*P* < 0.01; $$$*P* < 0.001 *versus* respective ND values.

**Supplemental Table S6 | Sequence of qPCR primers used in this study.** Related to Methods.

**Supplemental Figure 1 | Increased oxidative activity of muscles in hypokinetic *Hacd1*-KO mice.** Related to Figure 1.

**(A)** Mean speed of spontaneous nocturnal locomotor activity. **(B)** Time to exhaustion during endurance treadmill test (60% of maximal aerobic speed). **(C and D)** Representative immunoblots **(C)** and quantification **(D)** of major members of the insulin signaling pathway, normalized to the expression of ß-Actin, measured in superficial *gastrocnemius* muscles of fasted mice, 20 min after intraperitoneal injection of insulin. **(E)** Non-metabolizable [^14^C]-2-deoxy-D-glucose tolerance test. **(F)** Area under the curve of [^14^C]-2-deoxy-D-glucose clearance displayed in **(E)**. **(G)** Lactate dehydrogenase activity in *soleus* and superficial *gastrocnemius* (Gast.) muscles. **(H)** *Hk2, Pdk4* and *Glut4* mRNA expression normalized by geometrical mean of three independent reference genes in superficial *gastrocnemius* muscle of WT and *Hacd1*-KO mice. **(I-J)** Glycerol-3-phosphate dehydrogenase (GPDH), succinate dehydrogenase (SDH) and NADH dehydrogenase (NADH tetrazolium reductase reaction, NADH-TR) activity on *tibialis anterior* muscle sections of WT (upper panel) and *Hacd1*-KO (lower panel) mice **(I)** and on *biceps femoris* muscle sections of control (CTL; *HACD1^CNM/+^*; upper panel) and *HACD1*-mutant dogs (*CNM/CNM; HACD1^CNM/CNM^*; lower panel) **(J)**. **(K-M)** Myosin Heavy Chain (MHC) isoform distribution in superficial *gastrocnemius* **(K),** *tibialis anterior* **(L)** and *soleus* **(M)** muscles. **(N)** Quantitative analysis of fiber type distribution on *tibialis anterior* section detected by immunofluorescence. **(O)** *MyHC-I, MyHC-IIa*, *MyHCIIb* and *MyHC*-IIx mRNA expression normalized by the geometrical mean of three independent reference genes in *tibialis anterior* muscle. *n* = 4 per group for **A,** *n* = 8 per group for **B**, *n* = 6 per group for **C-H,** *n* = 3 per group for **I-J,** *n* = 9 per group for **K-M,** *n* = 3 per group for **N,** *n* = 6 per group for **O.** Scale bar in **I-J**: 200µm. Error bars, ± s.e.m.; \**P* < 0.05.

**Supplemental Figure 2 | Increased energy expenditure and reduced brown adipose tissue mass in *Hacd1*-KO mice.** Related to Figure 2.

**(A)** Main daily food intake per mouse over a 9-wk assessment period of normal diet (ND). **(B)** Main daily food intake per gram of body mass during the 9-wk assessment period of ND. **(C)** Analysis of variance of energy expenditure, independently of the type of diet. **(D)** Estimated resting metabolism during the assessment period of Figure 2E (ND) during daylight and night. **(E)** Analysis of variance of estimated resting metabolism, independently of the type of diet. **(F-G)** Respiratory exchange ratio **(F)** and fatty acid oxidation **(G)** under high-fat diet during daylight and night. **(H)** Rectal body temperature at rest and in active state. **(I and J)** BAT mass **(I)** and BAT mass relative to body mass **(J). (K)** *Ucp1* mRNA expression normalized by geometrical mean of three independent reference genes in the brown adipose tissue (BAT). **(L)** Gene ontology analysis performed with PANTHER software of the 72 upregulated proteins in *Hacd1*-KO mice *tibialis anterior* muscle compared to WT. *n* = 8 per group for **A-B** and **H-J**, *n* = 6 per group for **C-G and K**. *n* = 3 per group for **L**. Error bars, ± s.e.m.; \**P* < 0.05; \*\**P* < 0.01 *versus* WT. £*P* < 0.05 *versus* mice at rest.

**Supplemental Figure 3 | HACD1 deficiency impairs mitochondrial function in skeletal muscles.** Related to Figure 3.

**(A-B)** Representative immunoblots **(A)** and quantification **(B)** of citrate synthase (CS), ATP5A and VDAC normalized to β-actin in superficial *gastrocnemius* muscle. **(C)** Citrate synthase (CS) activity, a marker of mitochondrial mass, in *biceps femoris* muscle of control (CTL; *HACD1^CNM/+^*) and *HACD1*-mutant dogs (*CNM/CNM; HACD1^CNM/CNM^*). **(D)** ATP production measured on isolated mitochondria from *tibialis anterior* muscle of WT and *Hacd1*-KO mice. **(E)** NAD(P)H/NAD(P)+ ratio measured on isolated mitochondria from *tibialis anterior* muscle of WT and *Hacd1*-KO mice in state 4 (no ADP) and state 3 (2 mM ADP). **(F)** Δψm measured on isolated mitochondria from *tibialis anterior* muscle of WT and *Hacd1*-KO mice in state 4 (no ADP) and state 3 (2 mM ADP). **(G)** Phosphorylating respiration rates in the presence of octanoate in permeabilized myofibers freshly isolated from superficial *gastrocnemius* (Gast.) and *soleus* muscles. **(H)** β-hydroxyacyl-CoA dehydrogenase (HADHA) activity in isolated superficial *gastrocnemius* muscle. **(I)** Ratio (expressed in %) of phosphorylating respiration rates in the presence of PCoA versus octanoate from Figures 3H and S3G. **(J)** mRNA expression in superficial *gastrocnemius* muscle of a panel of genes involved in lipid metabolism, normalized by the geometrical mean of three independent reference genes. **(K** and **L)** Non-phosphorylating **(K)** and phosphorylating **(L)** respiration rates of saponin-permeabilized cardiac fibers of WT and *Hacd1*-KO mice in the presence of pyruvate. **(M)** Mitochondrial coupling of saponin-permeabilized cardiac fibers of WT and *Hacd1*-KO mice the presence of pyruvate (Acceptor Control Ratio (ACR) of phosphorylating to non-phosphorylating oxidation rates of pyruvate from **K** and **L**). **(N)** *Hacd1*-*FL*, *Hacd2*, *Hacd3* and *Hacd4* mRNA expression normalized to the geometrical mean of three independent reference genes in the superficial *gastrocnemius* muscle and heart. *Hacd1*-*FL* encodes the catalytically active isoform for HACD1. **(O)** mRNA expression of a panel of genes involved in mitochondrial biogenesis, lipid oxidation and lipogenesis, normalized by geometrical mean of three independent reference genes, in superficial *gastrocnemius* muscle (Gast.), heart, liver, gonadal white adipose tissue (WAT) and brown adipose tissue (BAT) of WT and *Hacd1*-KO mice.**(P-Q)** Representative immunoblots **(P)** and quantification **(Q)** of CS and VDAC, normalized to the expression of ß-Actin in heart, liver, white adipose tissue (WAT) and brown adipose tissue (BAT). *n* = 6 per group for **A-B** and **G-S**, *n* = 3 CTL and *n* = 4 *CNM/CNM* for **C,** *n* = 4 per group for **D-F**. Error bars, ± s.e.m.; \**P* < 0.05; \*\**P* < 0.01; \*\*\**P* < 0.001.

**Supplemental Figure 4 | Defective mitochondrial coupling in *Hacd1*-KO mice muscle is not associated with oxidative stress or *Ucp1*-*3* induction.** Related to Figure 3.

**(A)** Aconitase and fumarase enzymatic activities and ratio of aconitase to fumarase activity in isolated mitochondria from *tibialis anterior* muscle. **(B)** 9-hydroxyoctadecadienoic acid (9-HODE) and 13-hydroxyoctadecadienoic acid (13-HODE) quantification in isolated mitochondria from *tibialis anterior* muscle. **(C and D)** Quantitative bloting of protein carbonylation in superficial *gastrocnemius* muscle. **(E)** *Sod2*, *Gpx1*, *Pex11a*, *Pex19* and *Catalase* mRNA expression normalized by geometrical mean of three independent reference genes in superficial *gastrocnemius* muscle. **(F-H)** *Ucp1* **(F)**, *Ucp2* **(G)** and *Ucp3* **(H)** mRNA expression normalized by geometrical mean of three independent reference genes in superficial *gastrocnemius* muscle (Gast.), heart, liver, gonadal white adipose tissue (WAT) and brown adipose tissue (BAT) of *Hacd1*-KO mice, compared to WT mice set to 1.0. **(I)** Quantitative immunobloting of UCP3 in superficial *gastrocnemius* muscle. (**J**) Apparent K_m_ of oxygen consumption for ADP on permeabilized fibers from superficial *gastrocnemius* (*Gast*.) and *soleus* muscles in WT and *Hacd1*-KO mice. *n* = 6 per group for **A,** *n* = 5 per group for **B-D** *n* = 6 per group for **E-J**. Error bars, ± s.e.m.; \**P* < 0.05.

**Supplemental Figure 5 | Lipid and fatty acids content in mitochondria from *Hacd1*-KO mice muscle.** Related to Figure 4.

**(A)** Total fatty acid methyl ester (FAME) composition of mitochondria isolated from *tibialis anterior* muscle of *Hacd1*-KO mice, compared to WT mice, set to 1.0. **(B)** Fatty acids repartition from **(A)**. **(C** and **D)** Absolute **(C)** and relative **(D)** quantification of saturated (SFA), monounsaturated (MUFA) and polyunsaturated (PUFA) fatty acids from **(A)**. Absolute quantification is normalized to WT, set to 1.0. **(E** and **F)** Absolute **(E)** and relative **(F)** quantification of omega-3 (n-3), omega-6 (n-6), omega-7 (n-7) and omega-9 (n-9) fatty acids from **(A)**. Absolute quantification is normalized to WT, set to 1.0. **(G)** Total phospholipid content of mitochondria isolated from the heart, normalized to the mitochondrial protein content and compared to those of WT mice, set to 1.0. **(H)** Relative phospholipid species content of mitochondria isolated from the heart, normalized to the total content of mitochondrial phospholipids. CL = cardiolipins, PI = phosphatidylinositol, PS = phosphatidylserine, SM = sphingomyeline, PE = phosphatidylethanolamine, PC = phosphatidylcholine; *n* = 5 per group for **A-F**. *n* = 6 per group for **G-H**. Error bars, ± s.e.m. \**P* < 0.05. \*\**P* < 0.01.

**Supplemental Figure 6 | Cardiolipins composition in mitochondria from *Hacd1*-KO mice muscle.** Related to Figure 4.

**(A-D)** Relative cardiolipins composition of mitochondria isolated from *tibialis anterior* muscle of *Hacd1*-KO mice, compared to WT mice. Each cardiolipin species is expressed as a percentage of total cardiolipin content and separated as very low (≤0.1%, **A**), low (0.1%< x ≤0.3%, **B**), middle (0.3%< x ≤2%, **C**) and high (>2%, **D**) abundance. *n* = 5 per group for **A-D**; error bars: ± s.e.m.; multiple pair wises tests using Holm-Bonferroni method revealed no significant differences.

**Supplemental Figure 7 | Abnormal cristae in mitochondria of *Hacd1-KO* mice muscle and mitochondrial enrichment by phospholipids**. Related to Figure 5.

**(A)** Transmission electron microscopy of longitudinal sections of myofibers from *soleus* muscle of WT (upper panel) and *Hacd1*-KO (lower panel) mice. Images are from datasets taken at a low (x 2,500; left panel), intermediate (x 10,000; middle panel) and high (x 30,000; right panel) magnification. The Z-line is delimited by arrows. Cristae are regular tubular-shaped invaginations (white arrow heads) of the inner mitochondrial membrane, unambiguously identified on the same plane than the outer mitochondrial membrane (black arrow heads). Excessive dilation of cristae tips (asterisk) is frequently observed in *Hacd1*-KO myofibers. **(B** and **C)** Morphometric quantification in *soleus* muscle of the percentage of mitochondria containing cristae with excessively dilated tips (maximal diameter >15 nm) **(B)** and the mean maximal diameter of the tubular segment of cristae **(C**). **(D)** Representative images of mitoplasts made from native mitochondria or after fusion of mitochondria with TopFluor-Cardiolipin (CL) vesicles and stained with Mitotracker Red. Scale bar: 1µm. **(E-F)** Quantification of cardiolipin content with Nonyl-Aacridine Orange (NAO) on native mitochondria or after fusion with CL vesicles **(E)** and on mitoplasts isolated from the corresponding mitochondria **(F)**. **(G-H)** Oxidation rate (**G)** and ATP production **(H)** measured on native isolated mitochondria from *tibialis anterior* muscle or after fusion with cardiolipin (+CL) or phosphatidylcholine (+PC) vesicles. Scale bars in **A**: 100 nm. *n* = 4 per group for **A-H**. Error bars, ± s.e.m.; \**P* < 0.05; \*\*\**P* < 0.001 vs WT, $ = p ≤ 0.05 vs Native, $$ = p ≤ 0.01, $$$ = p ≤ 0.01 vs Native, £ = p ≤ 0.05, ££ = p ≤ 0.01, £££ = p ≤ 0.001 vs +CL.

**Supplemental Figure 8 | *Hacd1* deficiency results in protection against diet-induced obesity and insulin resistance.** Related to Figure 6.

**(A** and **B)** Main daily food intake during the 9-wk period of HFD per mouse **(A)** or per gram of body mass (BM) **(B)**. **(C)** Feed efficiency during the 9-wk period of HFD. **(D)** Fecal lipid excretion after 8 wk of HFD. **(E)** Rectal body temperature at rest and in active state after 9 wk of HFD. **(F** and **G)** Brown adipose tissue (BAT) mass **(F)** and BAT mass relative to body mass **(G)** after 9 wk of ND or HFD. **(H-J)** Serum cholesterol **(H)**, free fatty acids **(I)** and triglyceride **(J)** level after 9 wk of HFD. **(K)** Glycemia measured after food deprivation and assessed during 120 min after an intraperitoneal insulin injection at T_0_ (insulin sensitivity test). *n* = 8 per groupe for **A-K.** Error bars, ± s.e.m.; \**P* < 0.05; \*\**P* < 0.01 *versus* WT; £*P* < 0.05 *versus* mice at rest. $$*P* < 0.01; $$$*P* < 0.001 *versus* normal diet-fed mice.

## MATERIAL AND METHODS

### KEY RESSOURCES TABLE

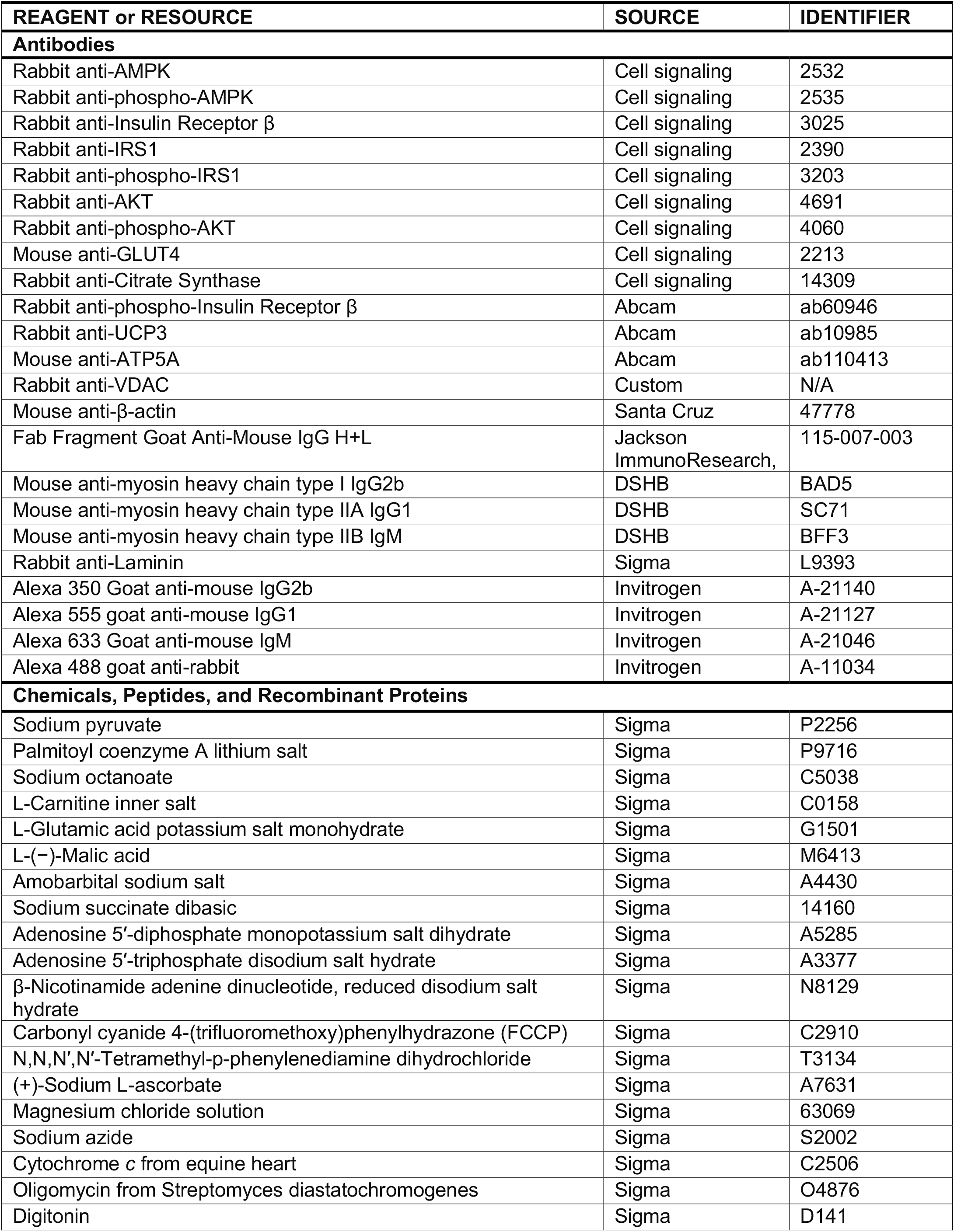
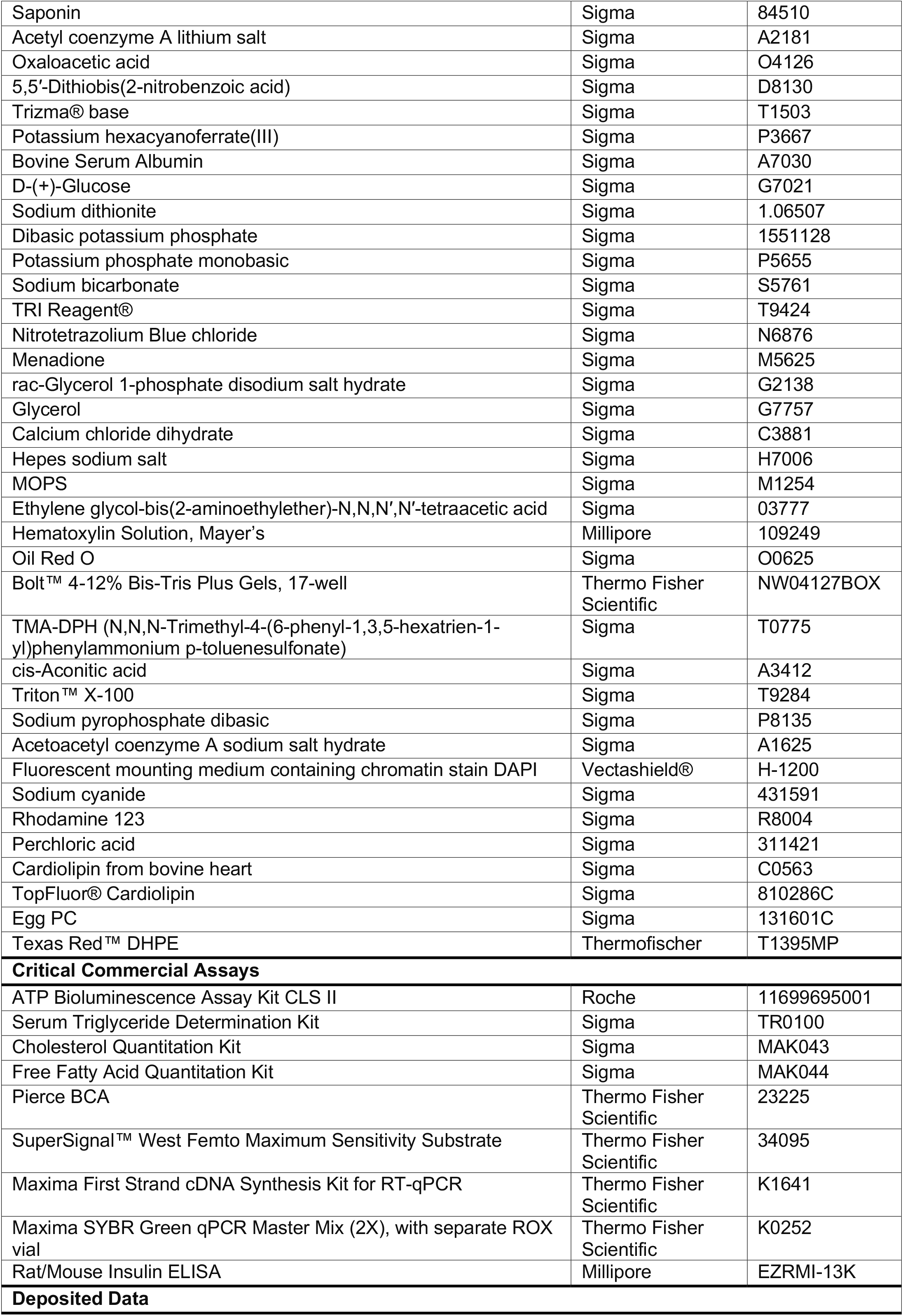
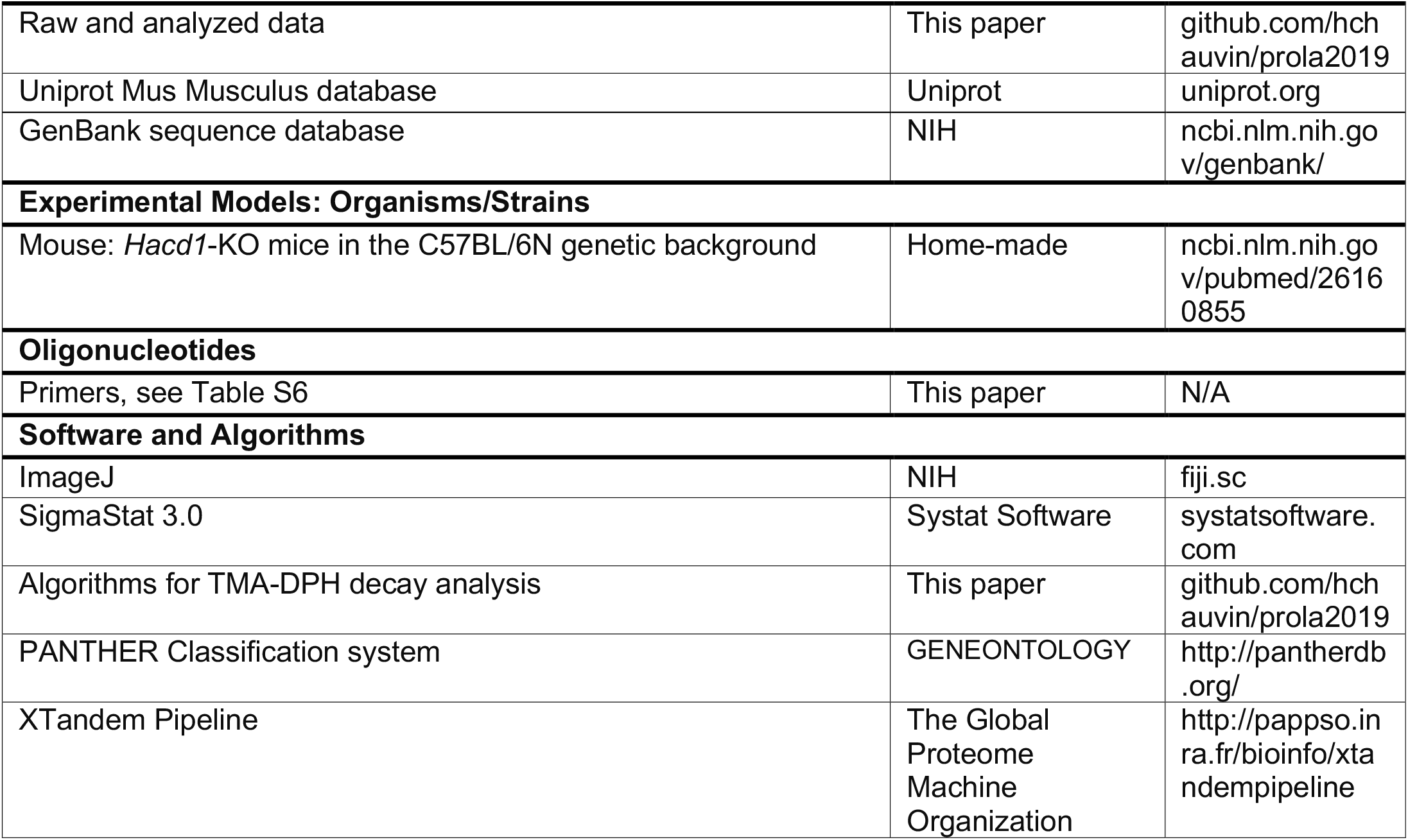

### CONTACT FOR REAGENTS AND RESSOURCES SHARING

Requests for information on methods and reagents should be directed to the Lead Contact, Fanny Pilot-Storck.

### EXPERIMENTAL MODEL AND SUBJECT DETAILS

#### Mice

Experiments on mice were approved by the Anses/EnvA/Upec Ethics Committee (C2EA – 16; approval number 11/11/15-2, 20/12/12-16 and 17-030) and all care and manipulations were performed in accordance with national and European legislation on animal experimentation. The whole study was carried on male mice that were housed in stainless steel cages containing environmental enrichment, in rooms maintained at 22 ± 1 °C with an air change rate of 12 vol/h and lit from 7am to 7pm. Food and water were given *ad libitum* unless otherwise stated. *Hacd1*-knockout (*Hacd1*-KO) mice were generated as previously described (Blondelle et al. 2015). Food was either a normal diet (ND, 3188 kcal/kg from carbohydrates (64%), proteins (24%) and fat (12%); maintenance diet for mice #C1324, Altronim) or a high-fat diet (HFD, 5241 kcal/kg from fat (60%), carbohydrates (23%) and protein (17%); Diet-Induced Obesity-Diet #C1090, Altronim). Kcal% was related to metabolizable energy (calculated values). Mice assessed for body mass gain under ND or HFD were housed individually and mice body mass and food intake were measured each week. Food efficiency was determined by dividing the body mass gain (mg) by the total calorie intake (kcal), as previously described in Butler et al. (2001). Mice were euthanized by cervical dislocation. The fast-twitch glycolytic superficial *gastrocnemius*, fast-twitch oxidative glycolytic (mixed) *tibialis anterior* and the slow-twitch *soleus* muscle were used to provide a representative panel of muscles with different myofiber composition. In the whole study, brown adipose tissue (BAT) refers to interscapular BAT.

#### Dogs

Experiments on dogs were approved by the Anses/EnvA/Upec Ethics Committee (C2EA – 16; approval number 20/12/12-18) and all care and manipulations were performed in accordance with national and European legislation on animal experimentation. Experiments in the present article were performed from thawed biopsies stored at −80°C and obtained in the context of a previous study (Walmsley et al., 2017), hence no additional experiment was performed on them.

## METHODS DETAILS

### Spontaneous locomotor activity

Spontaneous locomotor activity was measured over a 12 h night period using the Activmeter system (Bioseb). During the experiment, mice (4-to-5-mo-old) were individually caged with food and water provided *ad libitum*. The system allowed to quantify movements of the animal (distance, duration, slow *versus* fast movements, average speed of the animal when it is in motion) and the time and duration of immobility. After a 24 h-period of acclimation, locomotor activity was recorded during four consecutive nights.

### Treadmill test

The running capacity was evaluated on a treadmill (Tecmachine, Medical Development) at a 4% slope. All mice (4-to-5-mo-old) were initially acclimatized to the treadmill for 5 days (10 min/day, 12 m/min). The maximal aerobic speed was determined by challenging mice to an exercise of progressively increased intensity. Speed was increased every 2 min and the speed of the last completed running step before exhaustion was considered as the individual maximal aerobic speed. Endurance capacity was determined by challenging mice to a submaximal exercise intensity (60% of the maximal aerobic speed) until exhaustion.

### Glucose tolerance test

The glucose tolerance test was performed in mice (7-mo-old) fasted overnight for 16h prior to an intraperitoneal injection of a 40% glucose in normal saline solution at a dose of 3 g of glucose per kg of body mass for ND-fed mice or 1 g/kg glucose for HFD-fed mice. Blood was sampled from the tail vein just before the injection (T_0_) and then after 30, 60, 90 and 120 min in order to assay blood glucose concentration (Glycemia; Contour XT glucometer; Bayer).

### Insulin sensitivity test

The insulin sensitivity test was performed in mice (7-mo-old) fasted 4 h prior to an intraperitoneal injection of 0.1 UI/ml of insulin (Actrapid) in normal saline solution at a dose of 0.8 UI of insulin per kg of body mass for ND-fed mice and 1.2 UI/kg for HFD-fed mice. Blood was sampled from the tail vein just before the injection (T_0_) and then after 20, 40, 60, 90 and 120 min in order to assay blood glucose concentration (Glycemia; Contour XT glucometer; Bayer).

### Glucose tolerance test with 2-deoxyglucose

Animals (4-mo-old) were food-restricted for 4 h and a bolus of [^14^C]-2-deoxy-D-glucose ([^14^C]-2-DG) (32 µCi, Perkin Elmer) was injected concomitantly with a glucose injection (3 g/kg body mass). Blood samples (10 µl) were collected from the tail at 0, 5, 10, 20, 30, 40, 60 and 120 min.

Each blood sample was placed in a tube containing 50 µl of ZnSO4. After mixing by pipetting, 50 µl of Ba(OH)_2_ were added. The tube was then vortexed and centrifuged (2 min, 10000 rpm) and 50 µl of the supernatant were recovered and placed in a counting vial for evaporation overnight. After evaporation, the dry pellet was taken up with 500 μl of distilled water and the volume completed with scintillant up to 4.5 ml and vortexed again. Vials were analyzed on a TRI-CARB 2100TR liquid scintillation counter (PerkinElmer).

### 2-deoxyglucose 6-phosphate accumulation

Mice (4-mo-old) were euthanized 120 min after [^14^C]-2-DG injection, tissues were collected, weighted and then digested in NaOH 1 N 1 hr at 60 °C. The homogenate was precipitated in PCA 6% for 2-DG plus 2-DG-6P quantification or in Ba(OH)2 and ZnSO4 (0.3 N) for 2-DG quantification. Samples were then vortexed and centrifuged (2 min, 13000 g) and 400 µl of the supernatant were recovered and placed in a counting vial for evaporation overnight. After evaporation, the dry pellet was taken up with 500 μl of distilled water and the volume completed with scintillant up to 4.5 ml and vortexed again. Vials were analyzed on a TRI-CARB 2100TR liquid scintillation counter (PerkinElmer). The obtained value corresponding to 2-DG and 2-GD-6P content was then corrected by the value of 2-DG quantification and then normalized to tissue weight.

### Indirect calorimetry study

Mice (4-mo-old) were individually housed 14 days before and presented with training bottle and acclimated to the chambers for 48 h before experimental measurements. For the experiment, they were fed 7 days with normal diet and then 7 days with high-fat diet. They were analyzed for whole energy expenditure, oxygen consumption and carbon dioxide production, RQ (VCO_2_/VO_2_), food intake and locomotor activity (Beambreaks/h) using a calorimetric device (Labmaster, TSE Systems GmbH). Data were recorded every 15 min for each animal during the entire experiment. Data analysis was performed on excel XP using extracted raw value of VO_2_ consumed, VCO_2_ production (express in ml/h), and energy expenditure (kcal/h). Subsequently, each value was normalized to the whole lean body mass extracted from the EchoMRI (Whole Body Composition Analysers, EchoMRI, Houston, USA) analysis as previously described (Joly-Amado et al., 2012). No practical method of estimation of the basal metabolism is available presently (Even et al., 1994). However, an estimation could be made from windows of time during which animals should have no access to food, their previous intake had been far enough in time to discard the thermic effect of food and the spontaneous activity had been the lowest possible within the past 30 min. Data points of energy expenditure were considered as the best estimation of basal metabolism when the spontaneous activity was lower, within the last 30 min, than 5% of the highest daily value and when no intake was recorded within the last 60 min. Fatty acid oxidation was calculated from the following equation: fat ox (kcal/h) = energy expenditure (kcal/h) x (1-RER/0,3) according to (Bruss et al., 2010).

#### Data Analysis

The results are expressed as mean ± s.e.m.. Variance equality was analyzed by F test (Microsoft Excel), and comparisons between groups were carried out using Analysis of variance with food (ND or HFD) or phase (day or night) and genotype of mice (WT or *Hacd1*-KO mice) and their interactions as factors followed by post hoc Tukey test (Minitab 16, Paris France). Data of calorimetrics presented represent mean of at least 96 h measurement. A *P*-value < 0.05 (*) was considered statistically significant.

### Echocardiography

Mice (7-mo-old) were trained to be grasped because transthoracic echocardiography (TTE) was performed in non-sedated mice in order to avoid any cardiac depressor effect of anesthetic agents. Typical heart rates at recording were above 600 bpm. Mice were carefully caught by the left hand and placed in supine position. Images were acquired from a parasternal position at the level of the papillary muscles using a 13-MHz linear-array transducer with a digital ultrasound system (Vivid 7, GE Medical System, Horton, Norway). Left ventricular (LV) diameters, anterior and posterior wall thicknesses were serially obtained from M-mode acquisition. As we expected a homogenous function and planned to compare sequential echocardiography, we used M-mode technic to assess both LV volumes, mass, and ejection fraction. Relative LV wall thickness (RWT) was defined as the sum of septal and posterior wall thickness over LV end-diastolic diameter, and LV mass was determined using the uncorrected cube assumption formula [LV mass = (AWTdþ LVEDDþ PWTd)^3^ - (LVEDD)^3^]. Diastolic function was not assessed by echocardiography because of the heart rate above 600 bpm precluding the analysis of transmitral flow. Therefore, we assessed relaxation by dP/dtmin during *in vivo* hemodynamic analysis. Peak systolic values of radial SR in the anterior and posterior wall were obtained using Tissue Doppler Imaging (TDI) as previously described (Derumeaux et al., 2008). TDI loops were acquired from the same parasternal view with a careful alignment with the radial component of the deformation (Ferferieva et al., 2013) at a mean frame rate of 514 fps and a depth of 1 cm. The Nyquist velocity limit was set at 12 cm/s. Radial SR analysis was performed offline using the EchoPac Software (GE Medical Systems) by a single observer (G.D.) blinded to the genotype of the animals. Peak systolic of radial SR was computed from a region of interest positioned in the mid-anterior wall and was measured over an axial distance of 0.6 mm. The temporal smoothing filters were turned off for all measurements. Because slight respiratory variations exist, we averaged peak systolic of radial SR on eight consecutive cardiac cycles. The intra-observer variability of radial SR was assessed (G.D.) using the same acquisition and same method at 24-h intervals (3.5 ± 3.4% [3.3–3.7]).

### Rectal temperature

Rectal temperature of WT and *Hacd1*-KO mice (7-mo-old) was measured using a dedicated system (Bio-Thermo-Cis thermometer equipped with a Bio-Bret-3 rectal probe for mice; Bioseb) at rest (2pm) and during activity (7pm), in ND and after 9 wk of HFD.

### Extraction of total RNA, RT-PCR and RT-qPCR analyses

Samples from 4-mo-old mice were snap-frozen in liquid nitrogen and stored at - 80 °C. Total RNAs were isolated using TRIzol reagent (Sigma) according to the manufacturer’s protocol. Purity of RNAs was assessed by a ratio of absorbance at 260 nm and 230 nm >1.7. RNA quality was checked on agarose gel. One microgram of RNA was used for reverse transcription with the Maxima First Strand cDNA Synthesis Kit for RT-qPCR (Fermentas). cDNA were amplified using the Maxima SYBR Green qPCR Master Mix (2X; Fermentas). qPCR reactions were performed on a Roche Light Cycler system (Roche). All PCR and qPCR products were examined qualitatively on agarose gels. All presented RT-qPCR results were normalized by the geometrical mean of three independent reference genes (*Ywhaz*, *Polr2a*, *Rplpo*), as previously described (Vandesompele et al., 2002). Sequences of primers are listed in Table S6.

### Western blot experiments

For the analysis of Insulin receptor β, phosphor-IRS1, IRS1, phospho-AKT, AKT and GLUT4 protein content, mice (4-to-5-mo-old) fasted for 4 h were euthanized 20 min after intraperitoneal injection of insulin (0.8 UI per kg of body mass).

Frozen tissues from 4-to-7-mo-old mice were lysed in RIPA lysis buffer (50 mM Tris– HCl, pH 8, 150 mM NaCl, 1% Triton, 1 mM EDTA, 0.1% SDS, 0.5% deoxycholic acid), supplemented with a cocktail of protease and phosphatase inhibitors (Pierce), using a Precellys homogenizer (Bertin). Protein concentration was assessed using the bicinchoninic acid assay (Pierce). Protein extracts (20 μg) from superficial *gastrocnemius* muscles of 6-mo-old WT and *Hacd1*-KO mice were loaded on Bolt™ 4-12% Bis-Tris gels (Invitrogen), separated for 22 min at 200 V and subsequently transferred to PVDF membranes using transfer stack and iblot2 system for 7 min at 20 V (Invitrogen). Thereafter, blots were blocked for 60 min in 1.25% gelatin in Tris Buffer Saline-Tween at room temperature, followed by incubation with different primary antibodies (Insulin receptor β, phosphor-IRS1, IRS1, phospho-AKT, AKT, GLUT4, phospho-AMPK, AMPK and Citrate Synthase from Cell signaling; phospho-Insulin receptor β and ATP5A from Abcam; VDAC (custom-made, kind gift from Dr. C. Lemaire, INSERM U1180), overnight at 4 °C. After washing, membranes were incubated with the appropriate secondary antibodies for 60 min at room temperature and revealed using the West Femto chemiluminescent substrate (Pierce). Light emission was recorded using a chemiluminescent detection system (G-Box, Syngene) and quantified by the ImageJ software (v1.47, NIH). Protein content was normalized to β-actin (Antibody from Santa Cruz).

### Proteomic analyses

Muscle lysates of *tibialis anterior* from 4-mo-old mice were loaded on a gel and each lane of gel was cut and washed for 15 min with an acetonitrile (ACN) / 100 mM ammonium bicarbonate mixture (1:1). Digestion was performed in 50 mM ammonium bicarbonate pH 8.0 and the quantity of modified trypsin (Promega, sequencing grade) was 0.1 µg per sample. Digestion was achieved for 6 h at 37 °C. The supernatant was conserved. Peptides were extracted by 5% formic acid in water/ACN (v/v). Supernatant and extract tryptic peptides were dried and resuspended in 20 µl of 0.1% (v/v) trifluoroacetic acid (TFA). HPLC was performed on an Ultimate 3000 LC system (Dionex). A 4 µL sample was loaded at 20 µl/min on a precolumn cartridge (stationary phase: C18 PepMap 100, 5 µm; column: 300 µm i.d., 5 mm; Dionex) and desalted with 0.08% TFA and 2% ACN. After 4 min, the precolumn cartridge was connected to the separating PepMap C18 column (stationary phase: C18 PepMap 100, 3 µm; column: 75 µm i.d., 150 mm; Dionex). Buffers were 0.1% HCOOH, 2% ACN (A) and 0.1% HCOOH and 80% ACN (B). The peptide separation was achieved with a linear gradient from 0 to 36% B for 18 min at 300 nl/min. Including the regeneration step at 100% B and the equilibration step at 100% A, one run took 50 min. Eluted peptides were analyzed on-line with a LTQ-Orbitrap mass spectrometer (Thermo Electron) using a nanoelectrospray interface. Ionization (1.35 kV ionization potential) was performed with liquid junction and a capillary probe (10 µm i.d.; New Objective). Peptide ions were analyzed using Xcalibur 2.07 with the following data-dependent acquisition steps (1): full MS scan in Orbitrap (mass-to-charge ratio (m/z) 300 to 1600, profile mode) and (2) MS/MS in linear Trap (qz = 0.25, activation time = 30 ms, and collision energy = 35%; centroid mode). Steps 2 was repeated for the four major ions detected in step 1. Dynamic exclusion time was set to 60 s. A database search was performed with XTandem (X! tandem CYCLONE version 2010.12.01.1, http://www.thegpm.org/TANDEM/). Enzymatic cleavage was declared as a trypsin digestion with one possible miss cleavage. Cys carboxyamidomethylation and Met oxidation were set to static and possible modifications, respectively. Precursor mass and fragment mass tolerance were 10 ppm and 0.5 Da, respectively. A refinement search was added with similar parameters except that semi-trypsic peptide and possible N-ter proteins acetylation were searched. Few databases were used: the Mus musculus database (49728 entries, version March 2011 from Uniprot), contaminant database (trypsin, keratins…). Only peptides with a E value smaller than 0.1 were reported. Identified proteins were filtered and grouped using XTandem Pipeline (version 3.1.4) (http://pappso.inra.fr/bioinfo/xtandempipeline/) according to: (1) a minimum of two different peptides was required with a E value smaller than 0.05, (2) a protein E value (calculated as the product of unique peptide E values) smaller than 10^-4^. To take redundancy into account, proteins with at least one peptide in common were grouped. This allowed to group proteins of similar function. Within each group, proteins with at least one specific peptide relatively to other members of the group were reported as sub-groups. Gene Ontology analysis and protein set enrichment statistical analyses were performed using the PANTHER (Protein ANnotation THrough Evolutionary Relationship) classification system (www.pantherdb.org; (Mi et al., 2013)).

### ATP quantification

Muscle ATP content was quantified using the CLSII kit (Sigma). Mice (4-to-5-mo-old) were euthanized immediately after a submaximal treadmill exercise of 40 min at 19 m/min and 4% slope, and *tibialis anterior* muscle were removed and snap-frozen in liquid nitrogen. Frozen tissues were weighted, minced with scissors and homogenized in the lysis buffer provided in the kit using a dounce homogenizer. Homogenates were centrifuged at 15,000 g for 10 min and supernatants diluted 50:50 in the dilution buffer provided by the kit. ATP quantitation was performed according to the manufacturer’s instructions. For each sample, ATP concentration was normalized to the total protein content assessed with the bicinchoninic acid assay (Pierce).

### Muscle fibers respiration

The mitochondrial respiration was studied *in vitro* in saponin-skinned fibers as previously described (Kuznetsov et al., 2008). Briefly, fibers from 4-mo-old mice were separated under a binocular microscope in solution S (see below) on ice and then permeabilized in solution S containing 50 µg/ml of saponin for 30 min at 4 °C. After being placed 10 min in solution R (see below) to wash out adenine nucleotides and creatine phosphate, skinned separated fibers were transferred in a 3 ml water-jacketed oxygraphic cell (Strathkelvin Instruments) equipped with a Clark electrode, as previously described (Kuznetsov et al., 2008), under continuous stirring. Solution S contained: 2.77 mM CaK_2_EGTA, 7.23 mM K_2_EGTA, 20 mM taurine, 0.5 mM DTT, 20 mM imidazole, 50 mM potassium-methane sulfonate, 5.7 mM Na_2_ATP, 15 mM creatine-phosphate (final solution: pH 7.1, 160 mM ionic strength, 100 nM free Ca^2+^, 1 mM free Mg^2+^). Solution R contained 2.77 mM CaK_2_EGTA, 7.23 mM K_2_EGTA, 20 mM taurine, 0.5 mM DTT, 20 mM imidazole, 3 mM phosphate, 90 mM potassium-methane sulfonate, 10 mM sodium-methane sulfonate and 2 mg/ml fatty acid-free bovine serum albumin, (final solution: pH 7.1, 160 mM ionic strength, 100 nM free Ca^2+^, 1 mM free Mg^2+^). After the experiments, fibers were harvested and dried, and respiration rates expressed as µmoles of O_2_ per min and per g of dry mass.

#### Measurement of the maximal muscular oxidative capacities

After the determination of basal respiration rate measured at 22 °C with R solution plus 1 mM pyruvate and 4 mM malate as mitochondrial substrates (V0, non-phosphorylating rate), fibers were exposed to an increasing concentration of ADP to determine the apparent Km for ADP and maximal mitochondrial respiration (at saturating concentration of ADP (2 mM) Vmax, phosphorylating rate). The ratio of Vmax/V0 represented the degree of coupling between oxidation and phosphorylation of ADP for pyruvate.

#### Measurement of the respiratory chain complexes

While Vmax was recorded, electron flow went through complexes I, III, and IV. Then, four min after this Vmax measurement, complex II was stimulated with succinate (15 mM) (complexes I, II, III, IV). Complex I was then blocked with amytal (2 mM). In these conditions, mitochondrial respiration was evaluated by complexes II, III, and IV.

#### Measurement of mitochondrial substrate utilization

Experiments were started in solution R plus 4 mM malate, 2 mM carnitine and 0.1 mM palmitoyl-coenzyme A (PCoA). After the determination of basal respiration rate V0, the maximal fiber respiration rate of PCoA was measured in the presence of a saturating ADP concentration (2 mM). Then substrates were sequentially added every three to four min as follows: 0.1 mM octanoate, 1 mM pyruvate and 15 mM succinate. The ratio PCoA/V0 corresponded to the degree of coupling between oxidation and phosphorylation for this substrate.

### Isolation of skeletal muscles mitochondria and respiration measurement

*Tibialis anterior* muscles were quickly collected after euthanasia of mice (8-mo-old) in ice-cold PBS (Thermo Fisher Scientific) supplemented with 10 mM EDTA; muscle samples were then minced with scissors, rinsed and incubated in 5 ml of ice-cold PBS supplemented with 10 mM EDTA and 0.05% trypsin for 30 min, followed by a centrifugation at 200 g for 5 min. Pellets were suspended in Isolation Buffer 1 (67 mM sucrose, 50 mM KCl, 10 mM EDTA, 0.2% BSA and 50 mM Tris-HCl, pH 7.4) and homogenized with a dounce homogenizer (Sigma). Mitochondria were purified by a differential centrifugation at 700 g for 10 min, and mitochondria-containing supernatants were then centrifuged at 8,000 g for 10 min. The crude mitochondrial pellet was resuspended in an appropriate volume of Isolation Buffer 2 (250 mM sucrose, 3 mM EGTA and 10 mM Tris-HCl, pH 7.4). Protein concentration was assessed using the bicinchoninic acid assay (Pierce). The mitochondrial oxygen consumption flux was measured as previously described (Mourier et al., 2014) at 37 °C using 500 µg of crude mitochondria proteins diluted in 3 ml of mitochondrial respiration buffer (250 mM sucrose, 20 µM EGTA, 2 mM KH_2_PO_4_, 1 mM MgCl_2_ and 10 mM Tris-HCl, pH 7.4) in an oxygraphic cell (Strathkelvin Instruments). The oxygen consumption rate was measured using 4 mM malate plus either 1 mM pyruvate and or 1 mM pyruvate and 15 mM succinate. Oxygen consumption was assessed in the phosphorylating state with 4 mM of ADP or in the non-phosphorylating state by adding 2.5 µg/ml of oligomycin. Respiration was uncoupled by successive addition of FCCP up to 5 µM to reach maximal respiration.

### Measurement of ATP synthesis flux

Isolated mitochondria (65 µg/ml) from 4-mo-old mice were resuspended in the mitochondrial respiration buffer (see above). After addition of ADP (4 mM), pyruvate (10 mM), glutamate (5 mM) and malate (5 mM), the oxygen consumption and ATP synthesis rates were both measured. Aliquots were collected every 60 s and precipitated in 7% HClO_4_/25 mM EDTA, centrifuged at 16,000 g for 5 min and then neutralized with KOH 2 M, MOPS 0.3 M. The ATP content in these samples was determined using the CLSII kit (Sigma). In a parallel experiment, oligomycin (2.5 μg/ml) was added to the mitochondrial suspension to determine the non-oxidative ATP synthesis rate.

### Membrane potential measurement

The transmembrane potential variations (ΔΨ) was estimated in isolated mitochondria from 4-mo-old mice from the fluorescence quenching of the lipophilic cationic dye rhodamine 123 using a Hitachi F7000 fluorimeter. Isolated mitochondria (65 µg protein/ml) were incubated in the Mitochondrial Buffer (250 mM sucrose, 20 µM EGTA, 2 mM KH_2_PO_4_, 1 mM MgCl_2_ and 10 mM Tris-HCl, pH 7.4) containing pyruvate (10 mM), glutamate (5 mM), malate (5 mM), 0.66 µM of rhodamine 123 (Sigma) and thermostated at 37 °C. When added, ADP was 1 mM and oligomycin was 2.5 µg/ml. The rhodamine 123 fluorescence signal at each steady state (F) was recorded using an excitation wavelength of 485 nm and fluorescence emission was continuously detected at 500 nm. At the end of each experiment, the maximum fluorescence signal (Fmax) was monitored after complete de-energization of the mitochondria following addition of FCCP (6 µM). Then, the Fmax-F/Fmax difference at each steady state was calculated.

### NAD(P)H/NAD(P)+ Redox measurement

Isolated mitochondria (100 µg protein/ml) from 4-mo-old mice were incubated in the Mitochondrial Buffer (250 mM sucrose, 20 µM EGTA, 2 mM KH_2_PO_4_, 1 mM MgCl_2_ and 10 mM Tris-HCl, pH 7.4) thermostated at 37 °C. When added, pyruvate was 10 mM, glutamate 5 mM, malate 5 mM, ADP 4 mM and cyanide 4 mM. The NAD(P)H autofluorescence signal at each steady state (F) was recorded using an excitation wavelength of 340 nm and fluorescence emission was continuously detected at 450 nm. At the end of each experiment, the maximum fluorescence signal (Fmax) was monitored after complete reduction within the NAD(P)H/NAD(P)+ couple following addition of cyanide. The fluorescence signal recorded in the presence of mitochondria alone without respiratory substrate was used as the 0% reduction state for the NAD(P)H/NAD(P)+ couple. NAD(P)H/NAD(P)+ ratio was expressed as a percentage of reduction according to the following formula: % Reduction = (F-F0%)/(F100%-F0%)*100.

### Mitochondrial isolation from heart

Mice (8-mo-old) were euthanized and hearts were quickly collected in ice-cold PBS (Thermo Fisher Scientific) supplemented with 10 mM EDTA; heart samples were then transferred into Isolation Buffer (310 mM sucrose, 20 mM Tris–HCl, 1 mM EGTA, pH 7.2), minced with scissors and homogenized with a dounce homogenizer (Sigma). Mitochondria were purified by a differential centrifugation at 1,200 g for 10 min, and mitochondria-containing supernatants were then centrifuged at 12,000 g for 10 min. The crude mitochondrial pellet was resuspended in an appropriate volume of Isolation Buffer. Protein concentration was assessed using the bicinchoninic acid assay (Pierce).

### ATP synthase oligomerization

Three independent preparations of mitochondria isolated from the *tibialis anterior* muscle of 6-mo-old WT and *Hacd1*-KO mice were resuspended at a protein concentration of 10 mg/mL in extraction buffer (150 mM potassium acetate, 15 % (w/v) glycerol, 2mM ε-amino caproic acid, 30 mM HEPES pH 7.4) in the presence of protease inhibitors (Complete, Roche) and 1.5% digitonin (w/v). After an incubation of 30 min at 4°C, samples were centrifuged at 25,000 g for 30 min at 4°C. 10 µL of supernatant were mixed with 1 µL of 0.0125% (w/v) SERVA blue G and loaded on 3-12% Bis-Tris native gel (Novex, Life technologies). Anode buffer was prepared with a 20-fold dilution of Novex Running Buffer, and light blue cathode buffer was obtained by a 200-fold dilution of Novex cathode additive in anode buffer. Gels were run at 150V for 120 minutes. ATPase activities were revealed in gel as described in (Grandier-Vazeille, Xavier and Guérin, 1996).

### Measurement of Fatty Acid Oxidation

*Gastrocnemius* muscles from 12-mo-old mice were incubated in 25 ml Erlenmeyer flasks containing Krebs-Henseleit bicarbonate medium, pH 7.4. Flasks were maintained at 37 °C in an oxygenated atmosphere (O_2_/CO_2_ 95/5). For fatty acid oxidation determination, 1 pCi of ^14^C-linoleate was added to the medium and the flask was hermetically sealed with rubber caps with centrally-inserted plastic wells. After 20-30 min of incubation, 0.4 ml of 40% (w/v) HClO_4_ were added through the rubber cap and 0.4 ml of 1 M solution of hyamine hydroxide were deposed in the center of the well. The flasks were then shaked at RT for 1 h. The radioactivity present in the center wells (corresponding to the formed CO_2_) was determined and the flask content was harvested and centrifuged at 700 g for 15 min. The resulting supernatant was neutralized with KOH (5 M) and total ^14^C-labeled acid-soluble products were counted. Total fatty acid oxidation corresponded to the sum of the catabolism of labeled linoleate into CO_2_ and acid-soluble products.

### Mitoplast preparation

Freshly isolated mitochondrial pellets were incubated in 1 ml of H300 Mitoplast Buffer (sucrose 70 mM, mannitol 220 mM, Hepes 2 mM, pH 7.4) supplemented with 0.05% BSA and 0.12 µg of digitonin per µg of mitochondrial protein, for 15 min at 4 °C, with gentle shaking. Mitochondria were centrifugated at 8,000 g for 5 min and pellets were incubated in 1 ml of H40 Mitoplast Buffer (H300 buffer diluted 7.5 X in distilled water) for 15 min at 4 °C, with gentle shaking. Suspension was then homogenized with a dounce homogenizer (Sigma) and centrifuged at 8,000 g for 5 min. Pellets were resuspended in 1 ml of H40 buffer and centrifuged at 8,000 g for 5 min. Protein concentration was assessed using the bicinchoninic acid assay (Pierce).

### Physical properties of mitochondrial membranes (TMA-DPH decay-fluorescence)

1-(4-Trimethylammoniophenyl)-6-phenyl-1,3,5-hexatriene p-toluenesulfonate (TMA-DPH, Sigma Aldrich) was used to monitor physical properties of mitochondrial membranes of 7-to-8-mo-old mice. TMA-DPH is composed of a cationic substitute (TMA) that anchors at the polar heads of the membrane, while allowing the fluorescent hydrophobic probe DPH to be located in nonpolar regions. TMA-DPH labeled the outer mitochondrial membrane from non-permeabilized whole mitochondria, or the inner mitochondrial membrane from mitoplasts. To limit fluorescent noise, whole mitochondria and mitoplasts were resuspended in the Hypotonic Buffer (sucrose 9 mM, mannitol 29 mM, Hepes 0.3 mM, pH 7.4) as described hereafter.

Experiments were repeated five times using independent samples, each from different mice. Decay-fluorescence measurements were performed on each sample at 37 °C after a measurement of the instrumental response of the spectrofluorimeter used to measure fluorescence-decay (“prompt”). Thirty µl of sample (DO_600_ = 0.055) diluted in 2.97 ml of Hypotonic Buffer were introduced into spectroscopic quartz cuvette with an optical path length of one cm (VWR International).

Fluorescence-decay was measured by the time-correlated single-photon counting (TCSPC) method using a Horiba-Fluoromax-4® spectrofluorimeter (Horiba) equipped with a 370-nm laser diode (NanoLED C2, Horiba) as the source of excitation. Fluorescence decays were measured in TCSPC setup (Deltahub, Horiba). The Instrument Response Function (IRF) was about 160 ps (measured at 370 nm using the hypotonic buffer). Emission and excitation wavelength of TMA-DPH were respectively fixed at 370 nm and 431 ± 1.1 nm. Each decay curve corresponded to 10,000 counts.

### Quantitative investigation of fluorescence decay of TMA-DPH

#### Curve smoothing

For illustration purposes, we smoothed the decay for the representative experiments using the generalized additive model (GAM) of R statistical package “mgcv” with a Gaussian link and a *P*-spline smooth term. This smoothing was not used for the subsequent modelling steps below.

#### Finite exponential mixture model

We modelled the fluorescence decay obtained from the lipidomic analyses of whole mitochondria and mitoplasts using finite exponential mixtures (Klausner et al., 1980; Stubbs et al., 1995). Those mixtures have been proven to be identifiable (Barndorff-Nielsen, 1965). We accounted for the artefacts of the measuring apparatus by convolving the mixtures with the instrumental response (or “prompt”). The model was comprised of 2*N* parameters, with *N* the number of exponentials (each exponential has a scale and a characteristic time). We did not find necessary to account for neither a potential time shift nor for background noise with an additional constant, as those additions did not improve the fit (data not shown). More specifically, let *P*(*i*) be the “prompt” (instrumental response) and *Y*(*i*) the “decay” (experimental response) for channel *i*. The times of measurement are *δi*, with *δ* the time interval between channels. The decay *G(t)* at time *t* is given by a mixture of exponentials with characteristic times τ*_k_*and scaling parameters *C_k_* (ℋ is the Heaviside function):

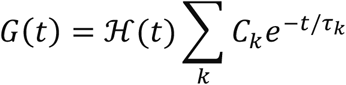

Convolving with the Prompt response *P*(*i*) gives the model, with *B_k_* = *δC_k_* scaling parameters:

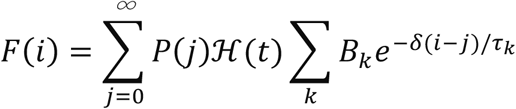

For the optimization procedures, we reparametrized the *τ_k_* to *θ_k_* = log *τ_k_*.

#### Non-linear mixed-effects models

To compare mixing proportions and characteristic times of decay, we used various non-linear mixed-effects models (see Listing S1): firstly, a baseline nonlinear mixed-effects model for either the outer or the inner membrane that did not discriminate WT and *Hacd1*-KO (model A); secondly, two models where either the mixing proportions or the characteristic times of decay alone could vary between WT and *Hacd1*-KO (respectively, models B1 and B2); finally, one model where both the mixing proportions and characteristic times were allowed to differ (model C).

More specifically, the mice phenotype (WT or *Hacd1*-KO) was encoded in a categorical factor with the following contrast matrix:

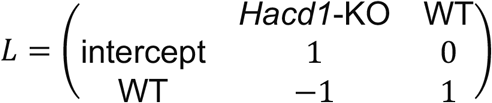

For instance, the mean θ_1_ for population *Hacd1-KO* is the intercept, and the mean θ_1_ for population WT is (intercept) + (WT).

#### Model fitting

We fitted the model through either a weighted nonlinear least-square method at the experiment level or nonlinear mixed-effects models (NLME) with potential linear regressors for the fixed parameters (Davidian and Giltinan, 2003; Pinheiro and Bates, 2000).

#### Statistical analysis

For model comparison, we performed an analysis of variance (ANOVA), privileging the reading of the Akaike’s Information Criterion over that of the Bayesian Information Criterion and the *P*-value of the log-likelihood-ratio (Burnham and Anderson, 2002).

#### Workflow

Firstly, we determined the number of exponentials to use by comparing the AIC between potential models and whether models converged when fitted through a Levenberg-Marquardt nonlinear least square method. Then, we determined the appropriate random-effects structure and heteroscedasticity correction for NLME modelling by assessing convergence, AIC, within-experiment and between-experiment heteroscedasticity for various combinations. Lastly, we added a linear dependence of (some of) the fixed effects to a dummy variable WT/*Hacd1*-KO and assessed the statistical significance of this addition with an analysis of variance (ANOVA).

#### Software implementation

The model is implemented in C++ using the Armadillo library for linear algebra (Sanderson and Curtin, 2016) and uses Armadillo’s Fast Fourier Transform to perform the discrete convolution with the instrumental response. Both the optimization and analysis were made in the R statistical language with package “nlme” (José C. Pinheiro and Douglas M. Bates, 2000). All the code is made available at <u>github.com/hchauvin/prola2019</u> with the appropriate tests and reproducible workflow (in addition, the releases are archived at 10.5281/zenodo.1228112).

### Lipidomic analyses

Mice (4-mo-old) were euthanized, and mitochondria were isolated from TA muscles as described above and immediately frozen in liquid nitrogen and protected from oxidation by a layer of argon gas, and stored at −80 °C until analysis. Quantitation of fatty acids and glycerophospholipids contents was performed as previously described for myoblasts (Blondelle et al., 2017). Phospholipid species representation was performed as previously described (Moulin et al., 2015). Briefly, total phospholipid amount was determined with Corona CAD detection by calculating the concentration of each PL class (in µg/ml) compared to calibration ranges of commercial standards. PL species intensities were determined by taking the mean relative intensity of m/z ions over the chromatographic peak of each PL class. Cardiolipin species identification was performed as described (Pulfer and Murphy, 2003) using high-resolution mass detection in full scan mode and MS2/MS3 fragmentation in data dependent mode. MSn fragmentation has not always been performed by the instrument due to very small amount of some ions. When no MSn data is available, CLs are mentioned as, e.g., CL (74:10). Otherwise, CL are mentioned as, e.g., CL (18:2-18:2-18:2-18:2). Sn1, sn2, sn3 and sn4 position of the acyl chains were not determined. Statistical analysis was performed using multiple pair wises tests according to the Holm-Bonferroni method.

### Mitochondrial phospholipids enrichment

Isolated mitochondria (500 µg) from 4-mo-old mice were incubated in Fusion Buffer (220 mM Mannitol, 70 mM Sucrose, 2 mM HEPES, 10 mM KH_2_PO_4_, 5 mM MgCl_2_, 1 mM EGTA, 10 mM Glutamate, 2 mM Malate, 10 mM Pyruvate and 2.5 mM ADP, pH 6.5) for 20 min at 30 °C under constant stirring agitation in the presence of 15 nmol small unilamellar vesicles (SUV), prepared as previously described (Mendez and Banerjee, 2017) and using natural or fluorescent phospholipids (Avanti Polar Lipids). After fusion, mitochondria were layered on a sucrose gradient (0.6 M) and centrifuged 10 min at 10,000g at 4 °C to remove SUV. Pellet was then washed in Mitochondrial Buffer 2 (250 mM sucrose, 3 mM EGTA and 10 mM Tris-HCl, pH 7.4).

### NAO quantification

Isolated mitochondria or mitoplasts (100 µg of proteins) from 4-mo-old mice were fixed in 2% PFA for 15 min, centrifuged at 10,000 g for 10 min and rinsed with PBS. The pellet was then resuspended and incubated 10 min at RT in 100 µl of PBS supplemented with 35 µM of Acridine Orange 10-Nonyl Bromide (NAO). Mitochondria or mitoplasts were then centrifuged at 10,000 g for 10 min, pellets were resuspended in 100 µl of PBS and fluorescence (λ excitation: 495 nm, λ emission: 620 nm) was acquired in 96-well plates with a spectrofluorimeter (TECAN infinite M200, TECAN, Austria).

### Serum chemistry

Serum insulin levels were determined in mice (7-mo-old) fasted overnight for 16h prior to an intraperitoneal injection of a normal saline solution containing 40% glucose, at a dose of 3 g of glucose per kg of body mass. Blood was collected before (Fasted) and 30 min after glucose injection (+Glucose), let at RT for 20 min for coagulation and centrifuged at 1000 g for 10 min. Serum insulin concentration was determined using the rat/mouse insulin ELISA kit (Millipore) according to the manufacturer’s protocol. Serum free fatty acids, triglycerides and cholesterol were assessed on mice fasted overnight for 16h by using NEFA assay kit, Triglycerides assay kit and Cholesterol assay kit, respectively (Sigma).

### Enzymatic assay

Complete enzymatic extractions from small pieces of frozen tissues from 4-mo-old mice or 5-mo-old dogs were obtained in an ice-cold buffer (one ml for 50 mg of tissue) containing 5 mM Hepes, pH 8.7, 1 mM EGTA, 1 mM DTT and 0.1% Triton X-100, using a Precellys homogenizer (Bertin), by 2 cycles of 8 sec at 6,500 rpm. Protein concentration was assessed using the bicinchoninic acid assay (Pierce). Total activity of cytochrome *c* oxidase (COX), citrate synthase (CS), β-hydroxyacyl-CoA dehydrogenase (HADHA) and lactate dehydrogenase (LDH) were assayed at 30 °C, pH 7.5, using standard spectrophotometric assays (Caffin et al., 2013). Total activities of aconitase and fumarase were assayed from mitochondrial extracts at 30 °C, pH 7.5, using standard spectrophotometric assays (Piquereau et al., 2012).

### Protein carbonylation

Muscle carbonylation of proteins was quantified using the Oxyblot Protein Oxidation Detection Kit (Millipore). Briefly, protein lysates from the superficial *gastrocnemius* muscle of 4-mo-old mice were prepared as for western blots and treated to derivatize carbonyl groups according to the manufacturer’s instructions. Then, protein extracts were separated by SDS-PAGE, transferred to a nitrocellulose membrane and incubated with the provided primary and secondary antibodies. Light emission was recorded using a chemiluminescent detection system (G-Box, Syngene) and quantified by the ImageJ software (v1.47, NIH). Membrane was stained with Coomassie Blue and protein content was normalized to one band.

### Histological staining

*Tibialis anterior* muscles from 7-mo-old mice were snap-frozen in isopentane cooled in liquid nitrogen and then stored at −80 °C. Ten µm-thick transverse sections were stained for Glycerol-3-phosphate dehydrogenase (GPDH), succinate dehydrogenase (SDH) and NADH dehydrogenase (NADH tetrazolium reductase reaction, NADH-TR) activity as previously described (Walmsley et al., 2017). Livers were frozen in OCT embedding medium (Tissue-Tek). Ten µm-thick transverse sections were fixed for 30 min in Baker solution (136 mM CaCl_2_, 4% HCHO), rinsed with water and then stained with Oil Red O (ORO) solution (12 mM ORO in 70% ethanol) for 5 min (liver) or 30 min (muscle). The adipose tissue from gonadal pads was fixed in 10% formalin for 48 h and embedded in paraffin. Four µm-thick transverse sections were stained with hematoxylin-eosin. Images were captured using an Axio Observer Z1 microscope (Zeiss) and analyzed in a blinded manner using the ImageJ software (v1.47, NIH).

### Analysis of the distribution of MHC isoforms

#### Electrophoresis of MHC isoforms

*Tibialis anterior*, superficial *gastrocnemius* and *soleus* muscles from 4-mo-old mice were subjected to an analysis of their constitutive MHC isoforms as previously described (Talmadge and Roy, 1993). Briefly, myofibrils were extracted from small sections of muscles in 7 vol of buffer solution (0.3 M NaCl, 0.1 M NaH_2_PO_4_, 0.05 M Na_2_HPO_4_, 0.01 M Na_4_P_2_O_7_, 1 mM MgCl_2_, 10 mM EDTA and 1.4 mM 2β-mercaptoethanol, pH 6.5). Myofibril samples were denatured with the sodium dodecyl sulfate (SDS) incubation medium according to the method of Laemmli. Electrophoresis was then performed using a Mini Protean II system (Bio-Rad) with separating and stacking gels containing 0.4% SDS and 8% and 0.16% acrylamide-bis (50:1), respectively. Gels were run at a constant voltage (72 V) for 31 h and then silver-stained at 4 °C. Bands corresponding to the distinct MHC isoforms were scanned and quantified using a densitometer GS 800, driven by the Quantify One 4.6.1 software (Bio-Rad).

#### Immunostaining of MHC isoforms

Muscle biopsy samples from mice (7-mo-old) were snap-frozen in isopentane cooled in liquid nitrogen and stored at −80 °C. Transverse-sections (7 μm thickness) were thawed, dried 45 min at RT and then rehydrated in PBS. Sections were permeabilized in PBS supplemented with triton 0.1% and blocked with PBS-1% BSA (Sigma) for 1 h at RT. Sections were rinsed with PBS and then incubated with Fab Fragment (Goat Anti-Mouse IgG H+L, Jackson ImmunoResearch, 1:100) for 30 min at RT before incubation with antibodies. Primary anti-myosin heavy chain type I (Mouse BAD5 IgG2b, DSHB, 1:40), anti-myosin heavy chain type IIA (Mouse SC71 IgG1 DSHB, 1:40), anti-myosin heavy chain type IIB (Mouse BFF3 IgM DSHB, 1:40) and anti-Laminin (Rabbit Sigma L9393, 1:1000) antibodies were incubated overnight at 4 °C and revealed using secondary Alexa 488 goat anti-rabbit (1:500), Alexa 350 Goat anti-mouse IgG2b (1:500), Alexa 555 goat anti-mouse IgG1 (1:500), Alexa 633 Goat anti-mouse IgM (1:500) from Invitrogen. Slides were mounted using Fluorescent mounting medium (Dako) containing DAPI (1.5 μg/ml) (Vectorshield). Images were captured using an Axio Observer Z1 microscope (Zeiss) and analyzed in a blinded manner using the ImageJ software (v1.47, NIH).

### Electron microscopy

After euthanasia of 5-to-6-mo-old mice, muscle tension and position were normalized by pining the entire skinned hindlimb on a paraffin-coated dish, with the metatarsus making an angle of 90 degree with the tibia. The limb was then fixed with 2% glutaraldehyde in 0.1 M sodium cacodylate buffer, pH 7.2, at room temperature for 4 h. Muscles were then cut in small pieces and kept overnight in the same fixation buffer at 4 °C. Samples were contrasted with Oolong Tea Extract 0.5% in cacodylate buffer; postfixed with a solution of 1% osmium tetroxide and 1.5% potassium cyanoferrate; gradually dehydrated in ethanol (30% to 100%) and substituted gradually in a mix of propylene oxyde-epon and embedded in Epon (Delta microscopy). Thin sections (70 nm) were collected onto 200 mesh copper grids, and counterstained with lead citrate. Grids were examined with a Hitachi HT7700 electron microscope operating at 80 kV (Elexience). Images were acquired with a charge-coupled device camera (AMT).

#### Data Analysis

For cristae quantification, 20 images at x 12,000 magnification were analyzed per animal and all visible cristae were assessed using the function plot profile on ImageJ (v1.47, NIH). Mean cristae width corresponded to the maximal diameter of the tubular part and was calculated for each crista of each analyzed mitochondrion. For each animal, a minimum of 100 cristae was measured for the superficialis *gastrocnemius* muscle and 800 cristae for the *soleus* muscle. All images were analyzed in a blinded manner.

### Fecal lipids quantitation

One gram of feces was collected after 8 wk of HFD (7-mo-old mice) and powdered using a tissue grinder. The powder was re-suspended in 5 ml of normal saline solution (NaCl 0.9%) and lipids were extracted in a mixture of chloroform:methanol (2:1). The solution was centrifuged at 1,000 g for 10 min. Lipids collected in the lower phase were air-dried and weighted.

## QUANTIFICATION AND STATISTICAL ANALYSIS

All data are presented as mean ± s.e.m.. Data were analysed using Sigma Stat (Sigma Stat, version 3.0, Systat Software, San Jose, CA, USA). Raw data from each individual experiment were evaluated using an unpaired two-tailed t test with 95% confidence. For datasets that did not pass the D’Agostino and Pearson omnibus normality test (alpha = 0.05), differences were evaluated using a two-tailed unpaired non-parametric Mann-Whitney test with 95% confidence. Multiple pair wise tests were corrected using Holm-Bonferroni method. Differences between groups were considered significant if the *P* value was less than 0.05.

## Notes

https://tinyurl.com/prola-tableS1

## REFERENCES

Acehan, D., Malhotra, A., Xu, Y., Ren, M., Stokes, D.L., and Schlame, M. (2011). Cardiolipin Affects the Supramolecular Organization of ATP Synthase in Mitochondria. Biophys. J. 100, 2184–2192.

Barndorff-Nielsen, O. (1965). Identifiability of mixtures of exponential families. J. Math. Anal. Appl. 12, 115–121.

Barth, P.G., Scholte, H.R., Berden, J.A., Moorsel, J.M.V.D.K.-V., Luyt-Houwen, I.E.M., Veer-Korthof, E.T.V., Harten, J.J.V.D., and Sobotka-Plojhar, M.A. (1983). An X-linked mitochondrial disease affecting cardiac muscle, skeletal muscle and neutrophil leucocytes. J. Neurol. Sci. 62, 327–355.

Bhandari, S., Lee, J.N., Kim, Y.-I., Nam, I.-K., Kim, S.-J., Kim, S.-J., Kwak, S., Oh, G.-S., Kim, H.-J., Yoo, H.J., et al. (2016). The fatty acid chain elongase, Elovl1, is required for kidney and swim bladder development during zebrafish embryogenesis. Organogenesis 12, 78–93.

Blondelle, J., Ohno, Y., Gache, V., Guyot, S., Storck, S., Blanchard-Gutton, N., Barthélémy, I., Walmsley, G., Rahier, A., Gadin, S., et al. (2015). HACD1, a regulator of membrane composition and fluidity, promotes myoblast fusion and skeletal muscle growth. J. Mol. Cell Biol. 7, 429–440.

Blondelle, J., Pais de Barros, J.-P., Pilot-Storck, F., and Tiret, L. (2017). Targeted lipidomic analysis of myoblasts by GC-MS and LC-MS/MS. In Skeletal Muscle Development, (James Ryall), pp. 39–60.

Boström, P., Wu, J., Jedrychowski, M.P., Korde, A., Ye, L., Lo, J.C., Rasbach, K.A., Boström, E.A., Choi, J.H., Long, J.Z., et al. (2012). A PGC1-α-dependent myokine that drives brown-fat-like development of white fat and thermogenesis. Nature 481, 463–468.

Bruss, M.D., Khambatta, C.F., Ruby, M.A., Aggarwal, I., and Hellerstein, M.K. (2010). Calorie restriction increases fatty acid synthesis and whole body fat oxidation rates. Am. J. Physiol. Endocrinol. Metab. 298, E108–116.

Caffin, F., Prola, A., Piquereau, J., Novotova, M., David, D., Garnier, A., Fortin, D., Alavi, M., Veksler, V., Ventura-Clapier, R., et al. (2013). Altered skeletal muscle mitochondrial biogenesis but improved endurance capacity in trained OPA1-deficient mice. J. Physiol. 591, 6017–6037.

Chen, W., and Guéron, M. (1992). The inhibition of bovine heart hexokinase by 2-deoxy-d-glucose-6-phosphate: characterization by 31P NMR and metabolic implications. Biochimie 74, 867–873.

Colman, E. (2007). Dinitrophenol and obesity: An early twentieth-century regulatory dilemma. Regul. Toxicol. Pharmacol. 48, 115–117.

Davidian, M., and Giltinan, D.M. (2003). Nonlinear models for repeated measurement data: An overview and update. J. Agric. Biol. Environ. Stat. 8, 387.

Denic, V., and Weissman, J.S. (2007). A molecular caliper mechanism for determining very long-chain fatty acid length. Cell 130, 663–677.

Derumeaux, G., Ichinose, F., Raher, M.J., Morgan, J.G., Coman, T., Lee, C., Cuesta, J.M., Thibault, H., Bloch, K.D., Picard, M.H., et al. (2008). Myocardial alterations in senescent mice and effect of exercise training: a strain rate imaging study. Circ. Cardiovasc. Imaging 1, 227–234.

Eckel, R.H., Grundy, S.M., and Zimmet, P.Z. (2005). The metabolic syndrome. Lancet Lond. Engl. 365, 1415–1428.

Even, P.C., Mokhtarian, A., and Pele, A. (1994). Practical aspects of indirect calorimetry in laboratory animals. Neurosci. Biobehav. Rev. 18, 435–447.

Ferferieva, V., Van den Bergh, A., Claus, P., Jasaityte, R., La Gerche, A., Rademakers, F., Herijgers, P., and D’hooge, J. (2013). Assessment of strain and strain rate by two-dimensional speckle tracking in mice: comparison with tissue Doppler echocardiography and conductance catheter measurements. Eur. Heart J. Cardiovasc. Imaging 14, 765–773.

Grandier-Vazeille, Xavier, and Guérin, M. (1996). Separation by blue native and colorless native polyacrylamide gel electrophoresis of the oxidative phosphorylation complexes of yeast mitochondria … - PubMed - NCBI. Anal. Biochem. Volume 242, 248–254.

Grundlingh, J., Dargan, P.I., El-Zanfaly, M., and Wood, D.M. (2011). 2,4-Dinitrophenol (DNP): A Weight Loss Agent with Significant Acute Toxicity and Risk of Death. J. Med. Toxicol. 7, 205–212.

Habersetzer, J., Ziani, W., Larrieu, I., Stines-Chaumeil, C., Giraud, M.-F., Brèthes, D., Dautant, A., and Paumard, P. (2013). ATP synthase oligomerization: from the enzyme models to the mitochondrial morphology. Int. J. Biochem. Cell Biol. 45, 99–105.

Hackenbrock, C.R. (1966). Ultrastructural bases for metabolically linked mechanical activity in mitochondria. I. Reversible ultrastructural changes with change in metabolic steady state in isolated liver mitochondria. J. Cell Biol. 30, 269–297.

Haines, T.H., and Dencher, N.A. (2002). Cardiolipin: a proton trap for oxidative phosphorylation. FEBS Lett. 528, 35–39.

Ikeda, M., Kanao, Y., Yamanaka, M., Sakuraba, H., Mizutani, Y., Igarashi, Y., and Kihara, A. (2008). Characterization of four mammalian 3-hydroxyacyl-CoA dehydratases involved in very long-chain fatty acid synthesis. FEBS Lett. 582, 2435–2440.

Ikon, N., and Ryan, R.O. (2017a). Barth Syndrome: Connecting Cardiolipin to Cardiomyopathy. Lipids 52, 99–108.

Ikon, N., and Ryan, R.O. (2017b). Cardiolipin and mitochondrial cristae organization. Biochim. Biophys. Acta BBA - Biomembr. 1859, 1156–1163.

Jastroch, M., Divakaruni, A.S., Mookerjee, S., Treberg, J.R., and Brand, M.D. (2010). Mitochondrial proton and electron leaks. Essays Biochem. 47, 53–67.

Joly-Amado, A., Denis, R.G.P., Castel, J., Lacombe, A., Cansell, C., Rouch, C., Kassis, N., Dairou, J., Cani, P.D., Ventura-Clapier, R., et al. (2012). Hypothalamic AgRP-neurons control peripheral substrate utilization and nutrient partitioning. EMBO J. 31, 4276–4288.

José C. Pinheiro and Douglas M. Bates (2000). Mixed-Effects Models in S and S-PLUS (New York: Springer-Verlag).

Karnovsky, M.J., Kleinfeld, A.M., Hoover, R.L., and Klausner, R.D. (1982). The concept of lipid domains in membranes. J. Cell Biol. 94, 1–6.

Kenneth P. Burnham and David R. Anderson (2002). Model Selection and Multimodel Inference - A Practical | Kenneth P. Burnham | Springer (New York: Springer-Verlag New York).

Khalifat, N., Puff, N., Bonneau, S., Fournier, J.-B., and Angelova, M.I. (2008). Membrane deformation under local pH gradient: mimicking mitochondrial cristae dynamics. Biophys. J. 95, 4924–4933.

Kihara, A. (2012). Very long-chain fatty acids: elongation, physiology and related disorders. J. Biochem. (Tokyo) 152, 387–395.

Kim, S.H., and Plutzky, J. (2016). Brown Fat and Browning for the Treatment of Obesity and Related Metabolic Disorders. Diabetes Metab. J. 40, 12–21.

Klausner, R.D., Kleinfeld, A.M., Hoover, R.L., and Karnovsky, M.J. (1980). Lipid domains in membranes. Evidence derived from structural perturbations induced by free fatty acids and lifetime heterogeneity analysis. J. Biol. Chem. 255, 1286–1295.

Kobayashi, S.D., and Nagiec, M.M. (2003). Ceramide/long-chain base phosphate rheostat in Saccharomyces cerevisiae: regulation of ceramide synthesis by Elo3p and Cka2p. Eukaryot. Cell 2, 284–294.

Kraschnewski, J.L., Boan, J., Esposito, J., Sherwood, N.E., Lehman, E.B., Kephart, D.K., and Sciamanna, C.N. (2010). Long-term weight loss maintenance in the United States. Int. J. Obes. 34, 1644–1654.

Kuznetsov, A.V., Veksler, V., Gellerich, F.N., Saks, V., Margreiter, R., and Kunz, W.S. (2008). Analysis of mitochondrial function in situ in permeabilized muscle fibers, tissues and cells. Nat. Protoc. 3, 965–976.

Lewis, R.N.A.H., and McElhaney, R.N. (2009). The physicochemical properties of cardiolipin bilayers and cardiolipin-containing lipid membranes. Biochim. Biophys. Acta 1788, 2069–2079.

Lizana, L., Bauer, B., and Orwar, O. (2008). Controlling the rates of biochemical reactions and signaling networks by shape and volume changes. Proc. Natl. Acad. Sci. U. S. A. 105, 4099–4104.

Mannella, C.A. (2006). Structure and dynamics of the mitochondrial inner membrane cristae. Biochim. Biophys. Acta 1763, 542–548.

Matsuzaka, T., Shimano, H., Yahagi, N., Kato, T., Atsumi, A., Yamamoto, T., Inoue, N., Ishikawa, M., Okada, S., Ishigaki, N., et al. (2007). Crucial role of a long-chain fatty acid elongase, Elovl6, in obesity-induced insulin resistance. Nat. Med. 13, 1193–1202.

Mayr, J.A., Havlícková, V., Zimmermann, F., Magler, I., Kaplanová, V., Jesina, P., Pecinová, A., Nusková, H., Koch, J., Sperl, W., et al. (2010). Mitochondrial ATP synthase deficiency due to a mutation in the ATP5E gene for the F1 epsilon subunit. Hum. Mol. Genet. 19, 3430–3439.

Mendez, R., and Banerjee, S. (2017). Sonication-Based Basic Protocol for Liposome Synthesis. Methods Mol. Biol. Clifton NJ 1609, 255–260.

Mi, H., Muruganujan, A., Casagrande, J.T., and Thomas, P.D. (2013). Large-scale gene function analysis with the PANTHER classification system. Nat. Protoc. 8, 1551–1566.

Mitchell, P. (1961). Coupling of phosphorylation to electron and hydrogen transfer by a chemi-osmotic type of mechanism. Nature 191, 144–148.

Mouchiroud, L., Eichner, L.J., Shaw, R.J., and Auwerx, J. (2014). Transcriptional coregulators: fine-tuning metabolism. Cell Metab. 20, 26–40.

Moulin, M., Solgadi, A., Veksler, V., Garnier, A., Ventura-Clapier, R., and Chaminade, P. (2015). Sex-specific cardiac cardiolipin remodelling after doxorubicin treatment. Biol. Sex Differ. 6, 20.

Mourier, A., Ruzzenente, B., Brandt, T., Kühlbrandt, W., and Larsson, N.-G. (2014). Loss of LRPPRC causes ATP synthase deficiency. Hum. Mol. Genet. 23, 2580–2592.

Muhammad, E., Reish, O., Ohno, Y., Scheetz, T., DeLuca, A., Searby, C., Regev, M., Benyamini, L., Fellig, Y., Kihara, A., et al. (2013). Congenital myopathy is caused by mutation of HACD1. Hum. Mol. Genet. 22, 5229.

Oemer, G., Lackner, K., Muigg, K., Krumschnabel, G., Watschinger, K., Sailer, S., Lindner, H., Gnaiger, E., Wortmann, S.B., Werner, E.R., et al. (2018). Molecular structural diversity of mitochondrial cardiolipins. Proc. Natl. Acad. Sci. 115, 4158–4163.

Ohno, Y., Suto, S., Yamanaka, M., Mizutani, Y., Mitsutake, S., Igarashi, Y., Sassa, T., and Kihara, A. (2010). ELOVL1 production of C24 acyl-CoAs is linked to C24 sphingolipid synthesis. Proc. Natl. Acad. Sci. U. S. A. 107, 18439–18444.

Paradies, G., Paradies, V., De Benedictis, V., Ruggiero, F.M., and Petrosillo, G. (2014). Functional role of cardiolipin in mitochondrial bioenergetics. Biochim. Biophys. Acta 1837, 408–417.

Pelé, M., Tiret, L., Kessler, J.-L., Blot, S., and Panthier, J.-J. (2005). SINE exonic insertion in the PTPLA gene leads to multiple splicing defects and segregates with the autosomal recessive centronuclear myopathy in dogs. Hum. Mol. Genet. 14, 1417–1427.

Perry, R.J., Zhang, D., Zhang, X.-M., Boyer, J.L., and Shulman, G.I. (2015). Controlled-release mitochondrial protonophore reverses diabetes and steatohepatitis in rats. Science 347, 1253– 1256.

Pinheiro, J.-C., and Bates, D.-M. (2000). Mixed-effects models in S and S-PLUS (Berlin: Springer).

Piquereau, J., Caffin, F., Novotova, M., Prola, A., Garnier, A., Mateo, P., Fortin, D., Huynh, L.H., Nicolas, V., Alavi, M.V., et al. (2012). Down-regulation of OPA1 alters mouse mitochondrial morphology, PTP function, and cardiac adaptation to pressure overload. Cardiovasc. Res. 94, 408–417.

Prats, M., Teissié, J., and Tocanne, J.-F. (1986). Lateral proton conduction at lipid–water interfaces and its implications for the chemiosmotic-coupling hypothesis. Nature 322, 756.

Pulfer, M., and Murphy, R.C. (2003). Electrospray mass spectrometry of phospholipids. Mass Spectrom. Rev. 22, 332–364.

Richter, E.A., and Hargreaves, M. (2013). Exercise, GLUT4, and Skeletal Muscle Glucose Uptake. Physiol. Rev. 93, 993–1017.

Rolfe, D.F., and Brown, G.C. (1997). Cellular energy utilization and molecular origin of standard metabolic rate in mammals. Physiol. Rev. 77, 731–758.

Sanderson, C., and Curtin, R. (2016). Armadillo: a template-based C++ library for linear algebra.

Sassa, T., Ohno, Y., Suzuki, S., Nomura, T., Nishioka, C., Kashiwagi, T., Hirayama, T., Akiyama, M., Taguchi, R., Shimizu, H., et al. (2013). Impaired epidermal permeability barrier in mice lacking elovl1, the gene responsible for very-long-chain fatty acid production. Mol. Cell. Biol. 33, 2787–2796.

Sawai, M., Uchida, Y., Ohno, Y., Miyamoto, M., Nishioka, C., Itohara, S., Sassa, T., and Kihara, A. (2017). The 3-hydroxyacyl-CoA dehydratases HACD1 and HACD2 exhibit functional redundancy and are active in a wide range of fatty acid elongation pathways. J. Biol. Chem. 292, 15538–15551.

Schneider, H., Lemasters, J.J., Höchli, M., and Hackenbrock, C.R. (1980). Liposome-mitochondrial inner membrane fusion. Lateral diffusion of integral electron transfer components. J. Biol. Chem. 255, 3748–3756.

Shrivastava, S., Dutta, D., and Chattopadhyay, A. (2016). Effect of local anesthetics on the organization and dynamics in membranes of varying phase: A fluorescence approach. Chem. Phys. Lipids 198, 21–27.

Song, D.H., Park, J., Maurer, L.L., Lu, W., Philbert, M.A., and Sastry, A.M. (2013). Biophysical significance of the inner mitochondrial membrane structure on the electrochemical potential of mitochondria. Phys. Rev. E Stat. Nonlin. Soft Matter Phys. 88, 062723.

Srikanthan, P., and Karlamangla, A.S. (2011). Relative muscle mass is inversely associated with insulin resistance and prediabetes. Findings from the third National Health and Nutrition Examination Survey. J. Clin. Endocrinol. Metab. 96, 2898–2903.

Stubbs, C.D., Ho, C., and Slater, S.J. (1995). Fluorescence techniques for probing water penetration into lipid bilayers. J. Fluoresc. 5, 19–28.

Talmadge, R.J., and Roy, R.R. (1993). Electrophoretic separation of rat skeletal muscle myosin heavy-chain isoforms. J. Appl. Physiol. Bethesda Md 1985 *75*, 2337–2340.

Teissié, J., Prats, M., Soucaille, P., and Tocanne, J.F. (1985). Evidence for conduction of protons along the interface between water and a polar lipid monolayer. Proc. Natl. Acad. Sci. 82, 3217– 3221.

Tiret, L., Blot, S., Kessler, J.-L., Gaillot, H., Breen, M., and Panthier, J.-J. (2003). The cnm locus, a canine homologue of human autosomal forms of centronuclear myopathy, maps to chromosome 2. Hum. Genet. 113, 297–306.

Toscano, A., Emmanuele, V., Savarese, M., Musumeci, O., Torella, A., Conca, E., Moggio, M., Nigro, V., and Rodolico, C. (2017). Pseudo-dominant inheritance of a novel homozygous HACD1 mutation associated with congenital myopathy: The first caucasian family. Neuromuscul. Disord. 27, S173.

Tseng, Y.-H., Cypess, A.M., and Kahn, C.R. (2010). Cellular bioenergetics as a target for obesity therapy. Nat. Rev. Drug Discov. 9, 465–482.

Vafai, S.B., and Mootha, V.K. (2012). Mitochondrial disorders as windows into an ancient organelle. Nature 491, 374–383.

Vandesompele, J., De Preter, K., Pattyn, F., Poppe, B., Van Roy, N., De Paepe, A., and Speleman, F. (2002). Accurate normalization of real-time quantitative RT-PCR data by geometric averaging of multiple internal control genes. Genome Biol. 3, RESEARCH0034.

Walmsley, G.L., Blot, S., Venner, K., Sewry, C., Laporte, J., Blondelle, J., Barthélémy, I., Maurer, M., Blanchard-Gutton, N., Pilot-Storck, F., et al. (2017). Progressive Structural Defects in Canine Centronuclear Myopathy Indicate a Role for HACD1 in Maintaining Skeletal Muscle Membrane Systems. Am. J. Pathol. 187, 441–456.

Wang, B., Pelletier, J., Massaad, M.J., Herscovics, A., and Shore, G.C. (2004). The yeast split-ubiquitin membrane protein two-hybrid screen identifies BAP31 as a regulator of the turnover of endoplasmic reticulum-associated protein tyrosine phosphatase-like B. Mol. Cell. Biol. 24, 2767– 2778.

Westerterp, K.R. (2010). Physical activity, food intake, and body weight regulation: insights from doubly labeled water studies. Nutr. Rev. 68, 148–154.

Yoshinaga, M.Y., Kellermann, M.Y., Valentine, D.L., and Valentine, R.C. (2016). Phospholipids and glycolipids mediate proton containment and circulation along the surface of energy-transducing membranes. Prog. Lipid Res. 64, 1–15.

Zhang, M., Mileykovskaya, E., and Dowhan, W. (2005). Cardiolipin is essential for organization of complexes III and IV into a supercomplex in intact yeast mitochondria. J. Biol. Chem. 280, 29403– 29408.

Zimmermann, C., Santos, A., Gable, K., Epstein, S., Gururaj, C., Chymkowitch, P., Pultz, D., Rødkær, S.V., Clay, L., Bjørås, M., et al. (2017). TORC1 Inhibits GSK3-Mediated Elo2 Phosphorylation to Regulate Very Long Chain Fatty Acid Synthesis and Autophagy. Cell Rep. 18, 2073–2074.

Zimorski, V., Ku, C., Martin, W.F., and Gould, S.B. (2014). Endosymbiotic theory for organelle origins. Curr. Opin. Microbiol. 22, 38–48.

