## Supplementary material for "Alteration of cardiolipin-dependent mitochondrial coupling in muscle protects against obesity": Supp Tables and Figures

**Table S2: Mitochondrial respiration rates.** Related to figure 3.

**Mitochondrial respiration rates for different complexes in WT or *Hacd1*-KO mice superficial *gastrocnemius* and *soleus* permeabilized muscle fibers.**

| | WT *Gast.* (6) | *Hacd1*-KO *Gast.* (6) | *P* |
|---|---|---|---|
| **Vo** | 2.16 ± 0.29 | 4.10 ± 0.45 | **0.003** |
| **Complex I** | 11.06 ± 0.89 | 9.37 ± 1.03 | 0.235 |
| **Complex I+II** | 16.11 ± 1.56 | 12.26 ± 1.42 | 0.089 |
| **Complex II** | 5.71 ± 0.54 | 3.45 ± 0.44 | **0.006** |
| **Ratio CxII/CxI** | 0.52 ± 0.04 | 0.37 ± 0.05 | **0.020** |
| **ACR (Complex I/Vo)** | 5.73 ± 0.94 | 2.71 ± 0.68 | **0.007** |
| | **WT *Soleus* (6)** | ***Hacd1*-KO *Soleus* (6)** | ***P*** |
| **Vo** | 4.21 ± 0.49 | 7.92 ± 0.55 | <**0.001** |
| **Complex I** | 19.72 ± 2.31 | 16.05 ± 1.39 | 0.263 |
| **Complex I+II** | 25.07 ± 2.78 | 23.46 ± 2.28 | 0.652 |
| **Complex II** | 13.78 ± 2.93 | 15.37 ± 1.62 | 0.534 |
| **Ratio CxII/CxI** | 0.68 ± 0.03 | 0.67 ± 0.05 | 0.878 |
| **ACR (Complex I/Vo)** | 4.83 ± 0.35 | 3.05 ± 0.28 | <**0.001** |
| | **WT Heart (4)** | ***Hacd1*-KO Heart (4)** | ***P*** |
| **Vo** | 5.40 ± 0.30 | 5.62 ± 0.32 | 0.611 |
| **Complex I** | 27.25 ± 1.28 | 29.67 ± 1.83 | 0.299 |
| **Complex I+II** | 37.34 ± 2.76 | 44.59 ± 3.80 | 0.207 |
| **ACR (Complex I/Vo)** | 5.22 ± 0.42 | 5.39 ± 0.38 | 0.782 |

**Mitochondrial substrate utilization in WT or *Hacd1*-KO mice superficial *gastrocnemius* and *soleus* permeabilized muscle fibers.**

| | WT *Gast.* (6) | *Hacd1*-KO *Gast.* (6) | *P* |
|---|---|---|---|
| **Vo** | 0.97 ± 0.16 | 2.07 ± 0.14 | **<0.001** |
| **Palmitoyl-CoA** | 2.27 ± 0.30 | 3.06 ± 0.35 | **0.043** |
| **+ octanoate** | 2.29 ± 0.36 | 3.10 ± 0.30 | **0.026** |
| **+ pyruvate** | 7.37 ± 0.52 | 7.49 ± 0.36 | 0.851 |
| **+ Glutamate/Succinate** | 10.27 ± 0.98 | 11.18 ± 0.62 | 0.425 |
| **ACR (PCoA/Vo)** | 3.07 ± 0.61 | 1.52 ± 0.16 | **0.003** |
| **% fatty acids** | 27.33 ± 4.81 | 28.33 ± 2.47 | 0.845 |
| **% pyruvate** | 59.03 ± 5.06 | 67.90 ± 2.67 | 0.112 |
| **% PCoA/Octanoate** | 97.52 ± 6.21 | 97.71 ± 1.76 | 0.975 |
| | **WT *Soleus* (6)** | ***Hacd1*-KO *Soleus* (6)** | ***P*** |
| **Vo** | 3.32 ± 0.81 | 6.04 ± 0.50 | **<0.001** |
| **Palmitoyl-CoA** | 12.83 ± 1.13 | 15.79 ± 1.03 | **0.049** |
| **+ octanoate** | 14.65 ± 0.75 | 17.89 ± 0.89 | **0.011** |
| **+ pyruvate** | 18.88 ± 1.17 | 20.44 ± 1.82 | 0.583 |
| **+ Glutamate/Succinate** | 24.14 ± 1.54 | 25.14 ± 1.90 | 0.931 |
| **ACR (PCoA/Vo)** | 3.85 ± 0.22 | 2.77 ± 0.26 | **0.004** |
| **% fatty acids** | 61.96 ± 3.50 | 72.69 ± 2.76 | **0.025** |
| **% pyruvate** | 78.48 ± 2.14 | 81.01 ± 2.12 | 0.583 |
| **% PCoA/Octanoate** | 86.03 ± 4.72 | 89.22 ± 5.93 | 0.678 |

**Mitochondrial respiration rates in isolated mitochondria from WT or *Hacd1*-KO mice TA.**

| | WT (4) | *Hacd1*-KO (4) | *P* |
|---|---|---|---|
| **Phosphorylating (State 3)** | 101.55 ± 19.72 | 48.03 ± 7.20 | **0.044** |
| **State 4** | 10.90 ± 0.87 | 12.75 ± 2.01 | 0.432 |
| **RCR** | 9.14 ± 1.24 | 3.12 ± 0.48 | **0.004** |
| **Non-phosphorylating** | 14.34 ± 1.70 | 11.25 ± 1.04 | 0.172 |
| **Uncoupled** | 189.42 ± 16.95 | 160.78 ± 16.48 | 0.153 |
| **Cyt-C stimulation (%)** | 7.80 ± 3.80 | 8.57 ± 3.98 | 0.667 |
| **$VO_2$ (for ATP/O determination)** | 129.87 ± 6.80 | 96.62 ± 10.57 | **0.038** |
| **ATP production (for ATP/O determination)** | 2.02 ± 0.24 | 0.89 ± 0.22 | **0.011** |

**Table S3: Echocardiographic parameters of WT and *Hacd1*-KO mice.** Related to figure 3.

| | **WT** | ***Hacd1*-KO** |
|---|---|---|
| **EF(%)** | 81.2 ± 2.3 | 83 ± 2.2 |
| **FS (%)** | 44.8 ± 2.5 | 46.7 ± 2.7 |
| **HR (bpm)** | 631 ± 9 | 624 ± 17 |
| **LVd (mm)** | 2.73 ± 0.11 | 2.88 ± 0.05 |
| **LVs (mm)** | 1.51 ± 0.10 | 1.54 ± 0.10 |
| **IVSd (mm)** | 0.83 ± 0.04 | 0.78 ± 0.03 |
| **IVSs (mm)** | 1.43 ± 0.05 | 1.44 ± 0.05 |
| **PWd (mm)** | 0.82 ± 0.03 | 0.80 ± 0.04 |
| **PWs (mm)** | 1.36 ± 0.07 | 1.48 ± 0.05 |
| **Strain rate ant (unit/s)** | 25.64 ± 1.40 | 23.25 ± 1.33 |
| **Strain rate post (unit/s)** | 25.91 ± 1.60 | 23.75 ± 1.08 |

**Table S4: Non-linear mixed-effects models of the fluorescence decay of TMA-DPH.** Related to figure 4.

**Non-linear mixed-effects models of the fluorescence decay from the analyses of mitoplasts.**

| | *Dependent variable:* Experimental response (event count/ns) | | | |
|---|---|---|---|---|
| | **A** | **B1** | **B2** | **C** |
| $\theta_1$ | 0.455∗∗∗ (0.153) | 0.505∗∗∗ (0.153) | | |
| $B_1$ | 0.025*** (0.001) | | 0.025*** (0.001) | |
| $\theta_2$ | −2.852∗∗∗ (0.242) | −2.631∗∗∗ (0.257) | | |
| $B_2$ | 0.0002∗ (0.0001) | | 0.0001∗ (0.0001) | |
| $\theta 3$ | −0.697∗∗∗ (0.087) | −0.667∗∗∗ (0.094) | | |
| $B_3$ | 0.016∗∗∗ (0.001) | | 0.016∗∗∗ (0.001) | |
| $B_1$ (inter.) | | 0.029∗∗∗ (0.002) | | 0.029∗∗∗ (0.002) |
| $B_1$ (WT) | | −0.007∗∗∗ (0.002) | | −0.007∗∗∗ (0.002) |
| $B_2$ (inter.) | | 0.001∗∗∗ (0.0002) | | 0.0005∗∗∗ (0.0002) |
| $B_2$ (WT) | | −0.0004 (0.0003) | | −0.0004 (0.0003) |
| $B_3$ (inter.) | | 0.018∗∗∗ (0.001) | | 0.018∗∗∗ (0.001) |
| $B_3$ (WT) | | −0.002 (0.002) | | −0.002 (0.002) |
| $\theta_1$ (inter.) | | | 0.752∗∗∗ (0.160) | 0.846∗∗∗ (0.172) |
| $\theta_1$ (WT) | | | −0.647∗∗∗ (0.226) | −0.708∗∗∗ (0.243) |
| $\theta_2$ (inter.) | | | −2.629∗∗∗ (0.316) | −2.432∗∗∗ (0.336) |
| $\theta_2$ (WT) | | | −0.509 (0.448) | −0.549 (0.476) |
| $\theta_3$ (inter.) | | | −0.727∗∗∗ (0.121) | −0.692∗∗∗ (0.126) |
| $\theta_3$ (WT) | | | 0.040 (0.171) | 0.028 (0.178) |
| Obs. | 57,344 | 57,344 | 57,344 | 57,344 |
| Log Likelihood | −136,223.600 | −136,134.000 | −136,268.900 | −136,157.800 |
| Akaike Inf. Crit. | 272,475.300 | 272,301.900 | 272,571.900 | 272,355.600 |
| Bayesian Inf. Crit. | 272,600.700 | 272,454.200 | 272,724.100 | 272,534.800 |

**Non-linear mixed-effects models of the fluorescence decay from the analyses of whole mitochondria.**

| | *Dependent variable:* Experimental response (event count/ns) | | | |
|---|---|---|---|---|
| | **A** | **B1** | **B2** | **C** |
| $\theta_1$ | 0.371∗∗∗ (0.043) | 0.370∗∗∗ (0.042) | | |
| $B_1$ | 0.019∗∗∗ (0.0004) | | 0.019∗∗∗ (0.0003) | |
| $\theta_2$ | -1.767∗∗∗ (0.030) | −1.764∗∗∗ (0.029) | | |
| $B_2$ | 0.001∗∗∗ (0.0001) | | 0.001∗∗∗ (0.0001) | |
| $\theta 3$ | −0.822∗∗∗ (0.020) | −0.822∗∗∗ (0.020) | | |
| $B_3$ | 0.017∗∗∗ (0.0003) | | 0.017∗∗∗ (0.0003) | |
| $B_1$ (inter.) | | 0.018∗∗∗ (0.0004) | | 0.018∗∗∗ (0.0004) |
| $B_1$ (WT) | | 0.002∗∗ (0.001) | | 0.001∗∗ (0.001) |
| $B_2$ (inter.) | | 0.001∗∗∗ (0.0001) | | 0.001∗∗∗ (0.0001) |
| $B_2$ (WT) | | −0.0003 (0.0002) | | −0.0002 (0.0002) |
| $B_3$ (inter.) | | 0.017∗∗∗ (0.0005) | | 0.016∗∗∗ (0.0005) |
| $B_3$ (WT) | | −0.0001 (0.001) | | 0.0001 (0.001) |
| $\theta_1$ (inter.) | | | 0.350∗∗∗ (0.060) | 0.363∗∗∗ (0.060) |
| $\theta_1$ (WT) | | | 0.033 (0.085) | 0.010 (0.084) |
| $\theta_2$ (inter.) | | | −1.812∗∗∗ (0.041) | −1.808∗∗∗ (0.040) |
| $\theta_2$ (WT) | | | 0.094 (0.058) | 0.083 (0.057) |
| $\theta_3$ (inter.) | | | −0.877∗∗∗ (0.025) | −0.869∗∗∗ (0.025) |
| $\theta_3$ (WT) | | | 0.104∗∗∗ (0.035) | 0.092∗∗∗ (0.035) |
| Obs. | 73,728 | 73,728 | 73,728 | 73,728 |
| Log Likelihood | −162,949.900 | −162,940.500 | −162,954.600 | −162,943.100 |
| Akaike Inf. Crit. | 325,927.700 | 325,915.000 | 325,943.200 | 325,926.100 |
| Bayesian Inf. Crit. | 326,056.600 | 326,071.600 | 326,099.700 | 326,110.300 |

**Table S5: Anatomical parameters of WT and *Hacd1*-KO mice after normal or high fat diet.** Related to figure 6.

| | WT ND | *Hacd1*-KO ND | WT HFD | *Hacd1*-KO HFD |
|---|---|---|---|---|
| **Eviscerated BM (g)** | 29.9 ± 1.4 | 26.3 ± 1.5 | 33.1 ± 2.1 | **25.8 ± 1.3** * |
| **Initial BM (g)** | 29.3 ± 0.9 | **26.3 ± 0.5** ** | 29.6 ± 0.6 | **25.4 ± 0.7** *** |
| **Final BM (g)** | 32.4 ± 1.2 | **28.8 ± 0.4** ** | **44.2 ± 1.3 $$$** | **31.4 ± 1.4** *** |
| **BM gain (g)** | 3.1 ± 0.6 | 2.5 ± 0.3 | **14.6 ± 1.0 $$$** | **6.3 ± 1.0 ***$** |
| **BM gain (%)** | 10.4 ± 1.8 | 9.7 ± 1.1 | **49.5 ± 3.2 $$$** | **25.3 ± 3.9 ***$$** |
| **Tibial length (cm)** | 1.78 ± 0.03 | 1.80 ± 0.04 | 1.74 ± 0.02 | 1.74 ± 0.02 |
| **Heart mass (mg)** | 130.3 ± 4.9 | 120.1 ± 3.6 | **153.7 ± 7.6 $** | **130.7 ± 13.5** ** |
| **Lung mass (mg)** | 185.8 ± 16.4 | 164.9 ± 11.6 | 273.3 ± 27.9 | 221.2 ± 9.9 |
| **Liver mass (mg)** | 1633.7 ± 85.1 | 1309.6 ± 39.0 | 1698.5 ± 118.2 | **1132.7 ± 103.1 $** |
| **Kidney mass (mg)** | 214.9 ± 11.2 | 185.9 ± 7.2 | 203.8 ± 5.8 | 205.2 ± 3.9 |
| ***Tibial anterior* (mg)** | 46.1 ± 1.3 | **37.5 ± 0.9** ** | 47.9 ± 1.8 | **40.8 ± 1.3** *** |
| **Gonadal fat pad (mg)** | 416 ± 82 | 279 ± 90 | **2182 ± 244 $$$** | **1079 ± 244 **$** |
| **Retroperitoneal fat pad (mg)** | 157.2 ± 47.3 | 94.3 ± 16.5 | **927.8 ± 83.0 $$$** | **353.3 ± 91.2** *** |
| **Mesenteric fat pad (mg)** | 567.2 ± 27.5 | 504.0 ± 42.9 | **942.7 ± 121.2 $$** | **452.0 ± 102.7** ** |
| **BAT mass (mg)** | 133.9 ± 16.9 | **79.3 ± 5.8** ** | **283.2 ± 26.2 $$$** | **174.1 ± 26.9 $$$**** |
| **HW/BM** | 4.21 ± 0.11 | **4.83 ± 0.14** * | **3.52 ± 0.15 $** | **3.96 ± 0.22 $$** |
| **Lung /BM** | 5.98 ± 0.44 | 6.61 ± 0.43 | 6.24 ± 0.49 | 6.66 ± 0.27 |
| **Liver/BM** | 52.63 ± 1.39 | 52.53 ± 0.82 | **38.57 ± 2.06 $** | **33.45 ± 1.76 $** |
| **Kidney/BM** | 6.92 ± 0.13 | 7.46 ± 0.26 | **4.70 ± 0.21 $** | 6.24 ± 0.33 |
| **TA/BM** | 1.75 ± 0.03 | **1.52 ± 0.04** ** | **1.11 ± 0.02 $$$** | **1.23 ± 0.00 $$$** |
| **Gonadal fat pad/BM** | 11.85 ± 2.03 | 9.13 ± 0.72 | **53.31 ± 5.04 $$$** | **34.26 ± 5.87 *$** |
| **Retroperitoneal fat pad/BM** | 4.44 ± 1.23 | 3.01 ± 0.37 | 22.81 ± 1.34 **$$$** | **11.13 ± 2.31 ***$$$** |
| **Mesenteric fat pad/BM** | 16.37 ± 0.59 | 16.27 ± 0.37 | **23.13 ± 1.94 $** | **14.31 ± 2.35** * |
| **BAT/BM** | 4.25 ± 0.40 | **3.18 ± 0.21** * | **6.43 ± 0.50 $$** | **5.05 ± 0.57 $$*** |

**Table S6: Sequence of qPCR primers used in this study.**
Related to Methods

| Gene | Forward primer | Reverse primer |
|---|---|---|
| *Acadm* | CCGTTCCCTCTCATCAAAAG | ACACCCATACGCCAACTCTT |
| *Acadvl* | TGGGCCTCTCTAATACCCAGT | TCCCAGGGTAACGCTAACAC |
| *Cd36* | CCACTGTGTACAGACAGTTTTGG | GCTCAAAGATGGCTCCATTG |
| *Catalase* | CCTCGTTCAGGATGTGGTTT | TCTGGTGATATCGTGGGTGA |
| *CoxI* | CACTAATAATCGGAGCCCCA | TTCATCCTGTTCCTGCTCCT |
| *Cpt1* | TCACCTGGGCTACACGGAGA | TCGGGGCTGGTCCTACACTT |
| *Cpt2* | ACCTGCTCGCTCAGGATAAA | AAACGACAGAGTCTCGAGCAG |
| *Fas* | GCAAGTGCAGCCTGAGGGAC | ACAGCCTGGGGTCATCTTTGC |
| *Glut4* | CATAGGAGCTGGTGTGGTCA | GATGGCACAGCCACACAT |
| *Gpx1* | GTTTCCCGTGCAATCAGTTC | TCACTTCGCACTTCTCAAACA |
| *Hacd1-FL* | ATGAAGAGAGCGTGGTGCTT | AAGGCGGCGTATATTGTGAG |
| *Hacd2* | TGCTATAGGGATTGTGCCATC | ACGGATAATTTCCGTGATTGTCC |
| *Hacd3* | GACGTGCAGAACCCTGCTATC | CTTCTGGACTGTGATGTTCACC |
| *Hacd4* | CAGCTCACAGAGAGAGTGATC | GAGTGTTTGACTGAGCCATGTC |
| *Hadha* | GCTTGGGAGGAGGACTTGA | AGCAACACTTCAGGGACACC |
| *Hk2* | GACCACATTGTCCAGTGCAT | TTTGTCCACTTGAGGAGGATG |
| *MyHC-I* | CCATCTCTGACAACGCCTATC | GGATGACCCTCTTAGTGTTGA |
| *MyHC-IIa* | TCAGGCTTCAGGATTTGGTG | GGATCTTGCGGAACTTGGATAG |
| *MyHC-IIb* | TCTGGCTTTCCTATTTTCTGGG | GTGCTCTTCAAGTTGGTCATC |
| *MyHC-IIx* | ATGTTCCTGTGGATGGTCAC | CTCGTTGGTGAAGTTGATGC |
| *Nrf1* | TTACTCTGCTGTGGCTGATGG | CCTCTGATGCTTGCGTCGTCT |
| *Pdk4* | CCTTCACACCTTCACCACAT | AAAGAGGCGGTCAGTAATCC |
| *Pex11a* | TCAAGAGGCTGGAGACCAGT | CGGTTGAGGTTGGCTAATGT |
| *Pex19* | TGTACCCATCCCTGAAGGAG | TGTGCTGCTGCTGGTACTTC |
| *Polr2a* | CAACATGCTGACAGATATGACC | TGATGATCTTCTTCTTGTTGTCTG |
| *Ppara* | AAGTGCCTGTCTGTCGGGAT | GCTTCGTGGATTCTCTTGCC |
| *Ppard* | GGGGACCAGAACACACGCTT | CCGCACACCCGACATTCCAT |
| *Ppargc1a* | CACCAAACCCACAGAGAACAG | GCAGTTCCAGAGAGTTCCACA |
| *Ppargc1b* | TGGAAAGCCCCTGTGAGAGT | TTGTATGGAGGTGTGGTGGG |
| *Rpl32* | GCTGCTGATGTGCAACAAA | GGGATTGGTGACTCTGATGG |
| *Rplpo* | GCGACCTGGAAGTCCAACTA | TTGTCTGCTCCCACAATGAA |
| *Sod2* | AGGAGCAAGGTCGCTTACAG | GTAGTAAGCGTGCTCCCACA |
| *Scd1* | CTACATGACCAGCGCTCTG | CGTACACGTCATTCTGGAACG |
| *Scd2* | GTAATCAGCGCCCTGGGCAT | CATACACGTCATTCTGGAACGC |
| *Srebp-1c* | CTGGAGACATCGCAAACAAGCTG | TGAGGTTCCAAAGCAGACTGCAG |
| *Tfam* | GCTGATGGGTATGGAGAAG | GAGCCGAATCATCCTTTGC |
| *Ucp1* | AGGGAGAGAAACACCTGCC | GCATTCTGACCTTCACGACC |
| *Ucp2* | CAGATGTGGTAAAGGTCCGC | CAGAAGTGAAGTGGCAAGGG |
| *Ucp3* | ATGGTTGGACTTCAGCCCTC | TGGGGGTGTAGAACTGCTTG |

**Supplemental Listing S1 | Final non linear mixed effects models of the fluorescence decay of TMA-DPH on mitoplasts and whole mitochondria.**
Related to Figure 4. *(in the R statistical language).*

```
library(nlme)

# Shared coefficients between Hacd1-KO and WT
model_A <- nlme(
  Decay ~ ...,
  fixed = theta1 + B1 + theta2 + B2 + theta3 + B3 ~ 1,
  random = list(
    File = pdDiag(B1 + B2 + B3 + theta1 + theta2 + theta3 ~ 1)
  ),
  weights = varConstPower(fixed = c(power = 0.5), const = 1, form = ~ fitted(.))
)

# Only Bs (resolving to mixing proportions) allowed to be different
# in Hacd1-KO and WT populations
model_B1 <- update(
  model_A,
  fixed = list(
    theta1 + theta2 + theta3 ~ 1,
    B1 + B2 + B3 ~ Population
  )
)

# Only thetas (resolving to characteristic times) allowed to be different
model_B2 <- update(
  model_A,
  fixed = list(
    theta1 + theta2 + theta3 ~ Population,
    B1 + B2 + B3 ~ 1
  )
)

# All coefficients allowed to be different
model_C <- update(
  model_A,
  fixed = theta1 + B1 + theta2 + B2 + theta3 + B3 ~ Population
)
```

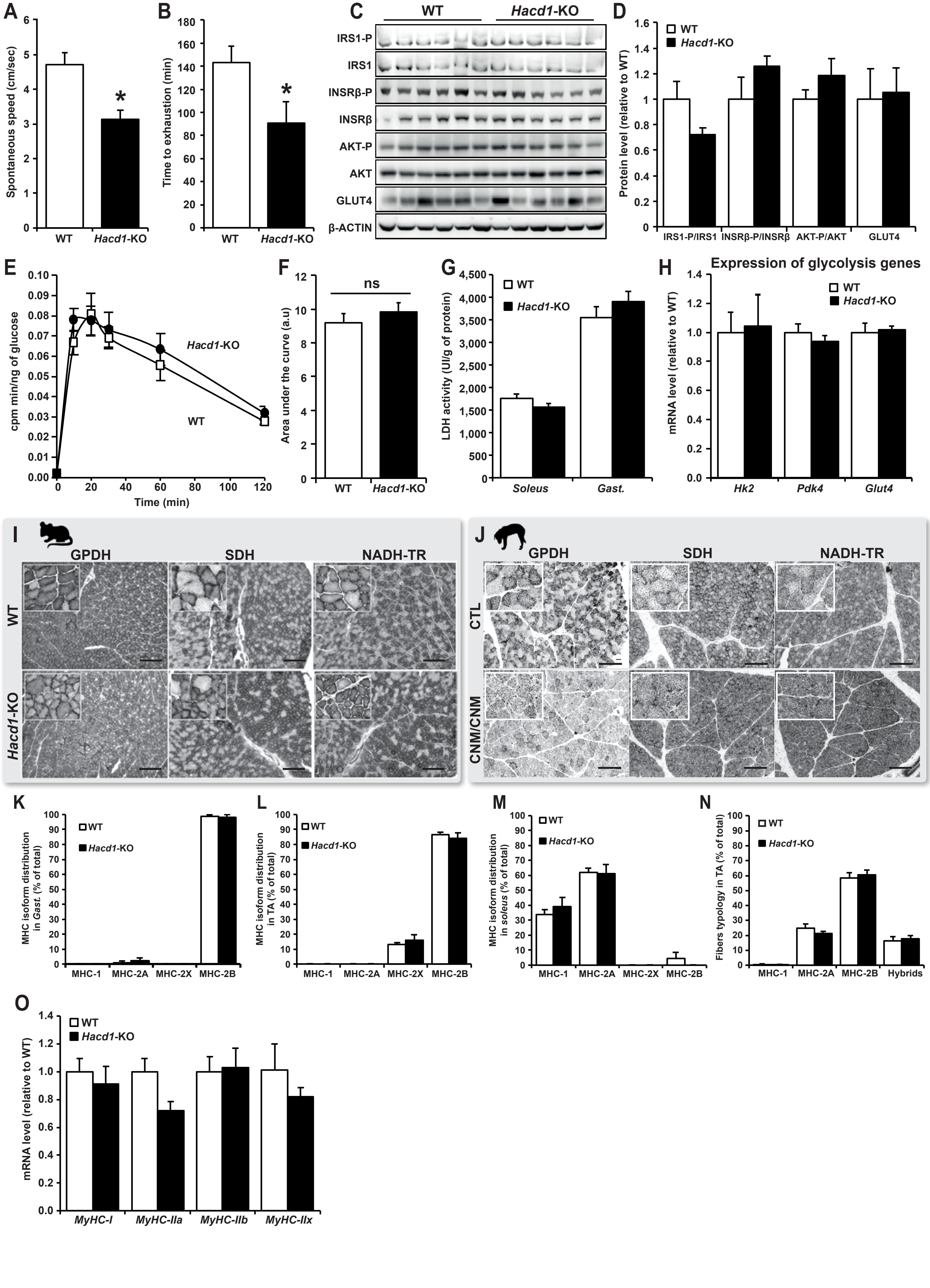


Figure S1 Prola

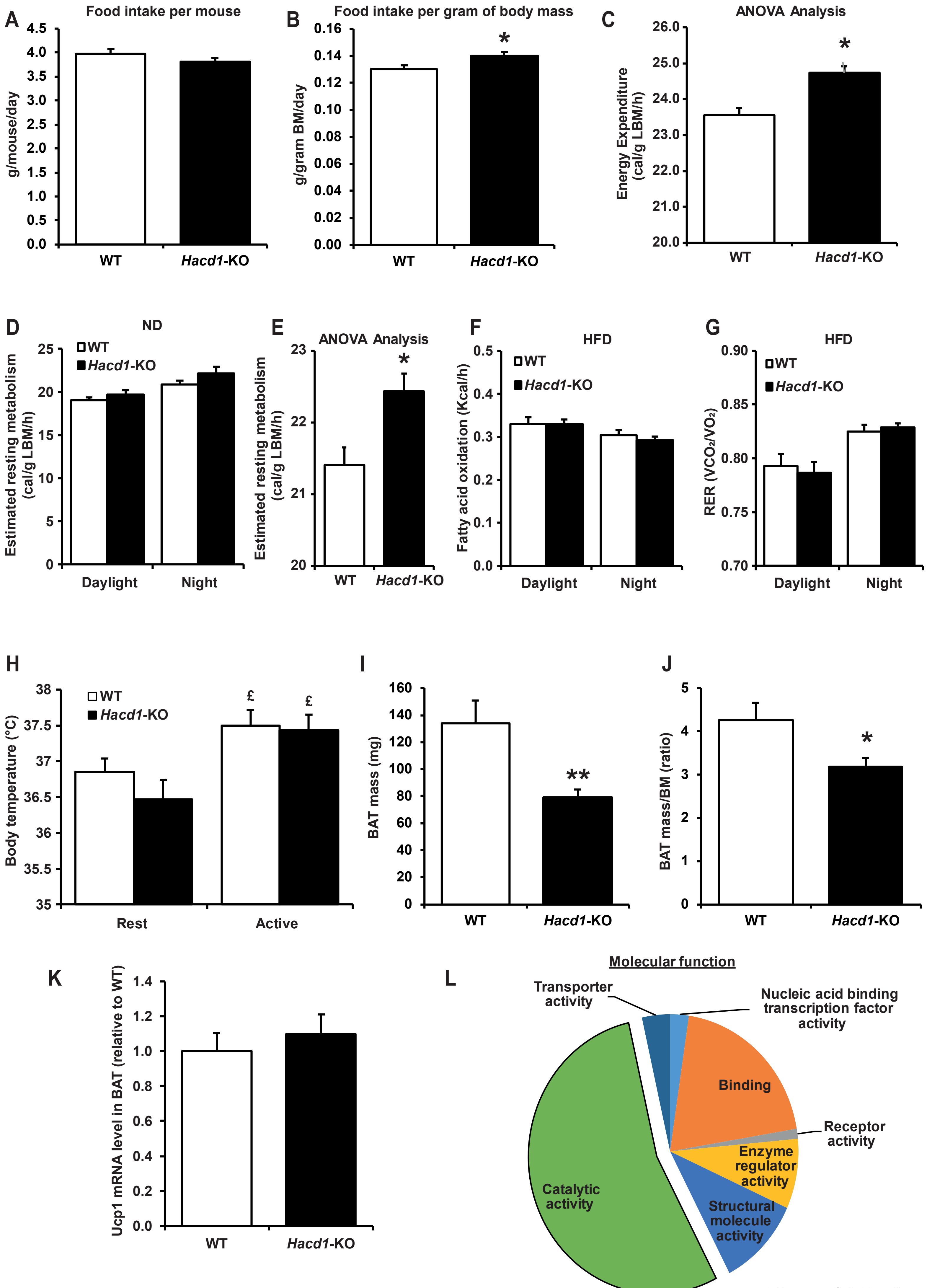


Figure S2 Prola

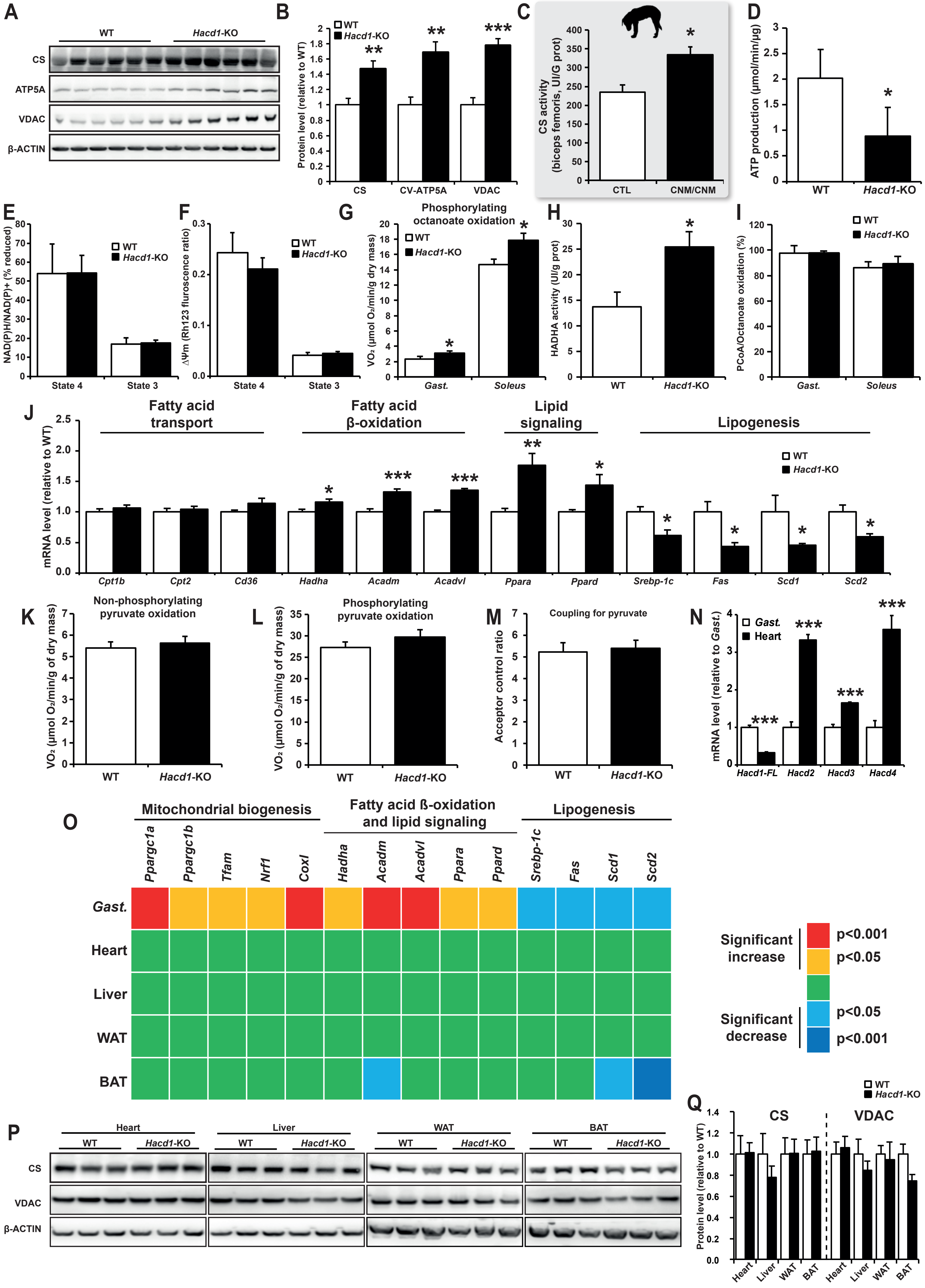


Figure S3 Prola

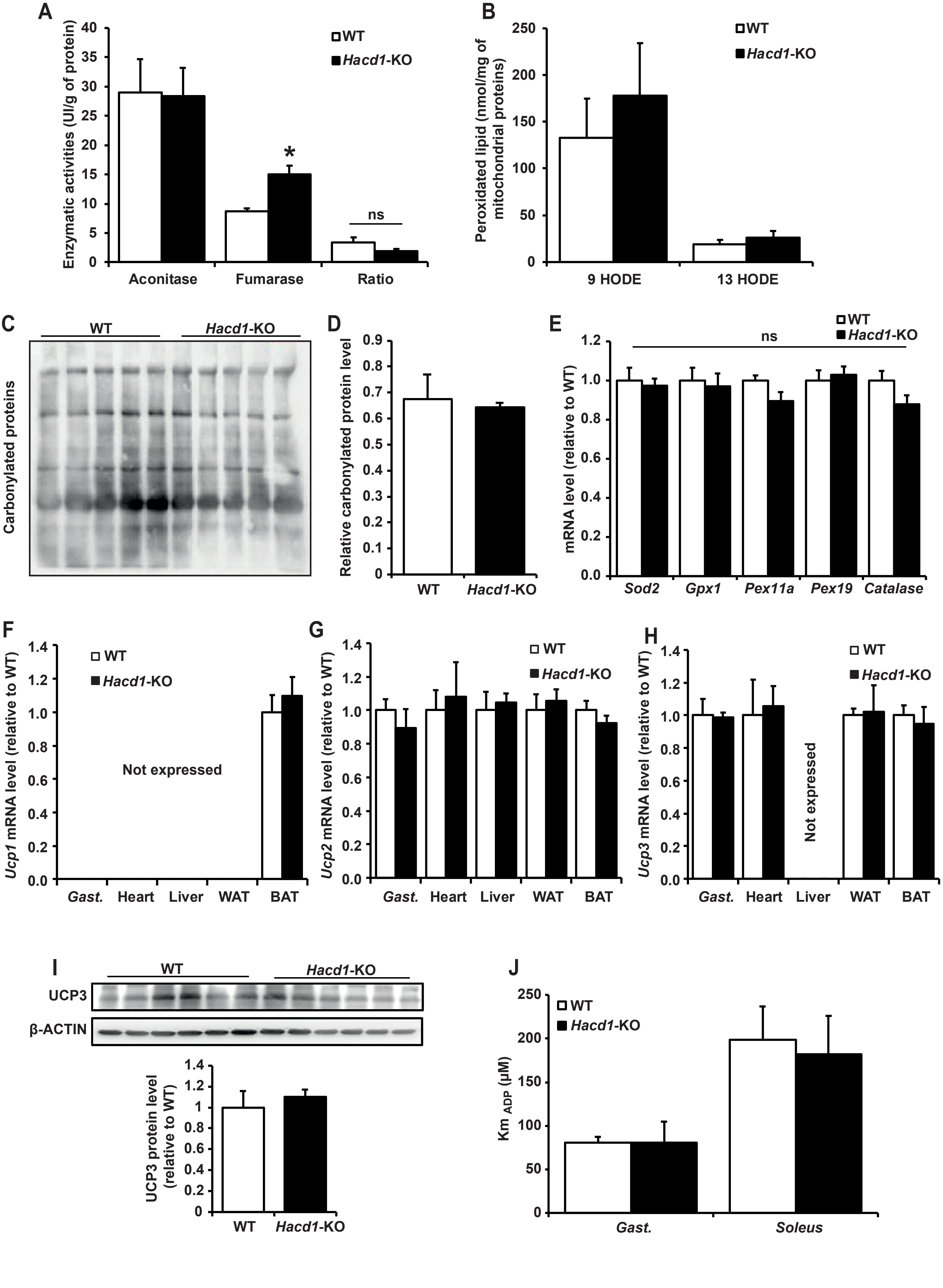


Figure S4 Prola

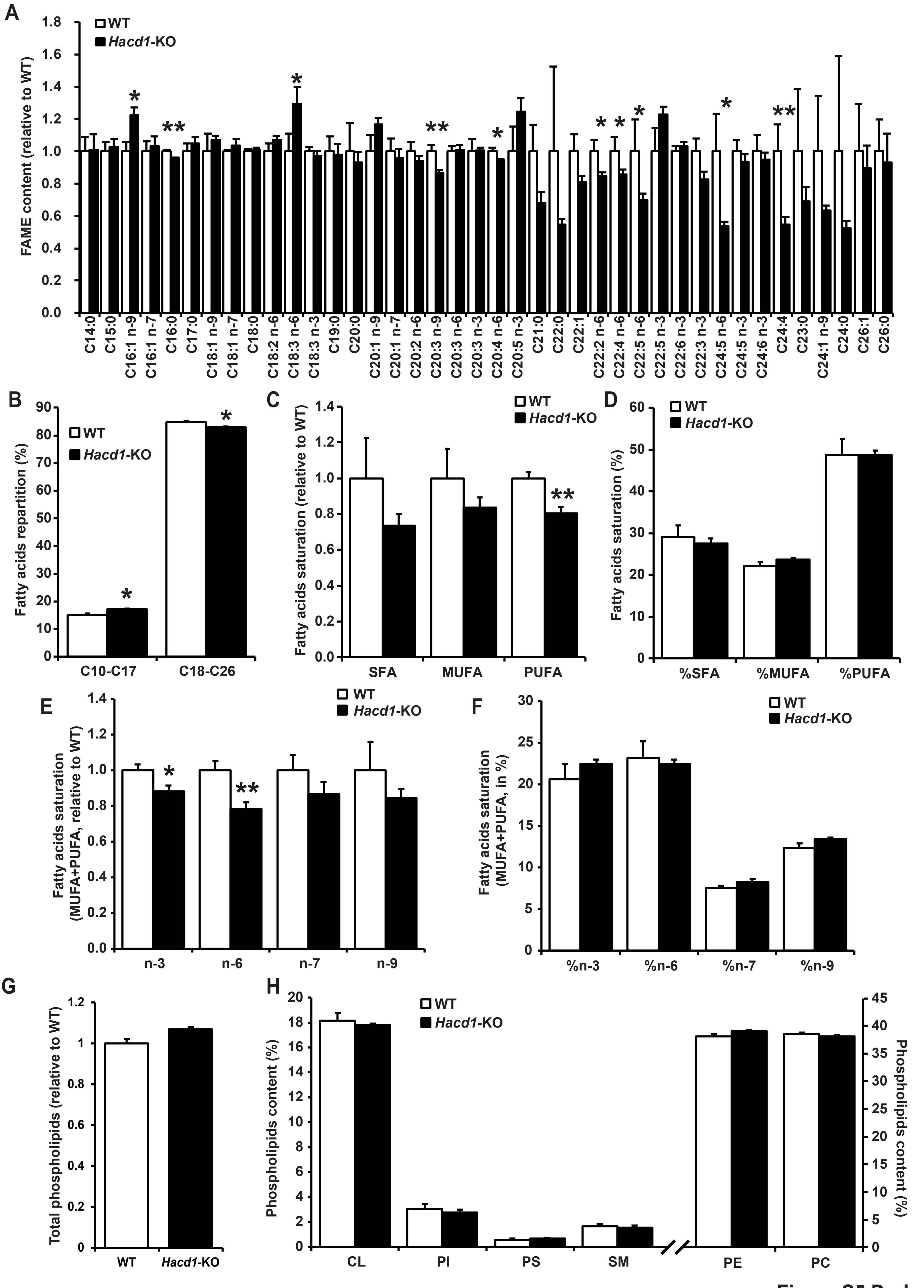


Figure S5 Prola

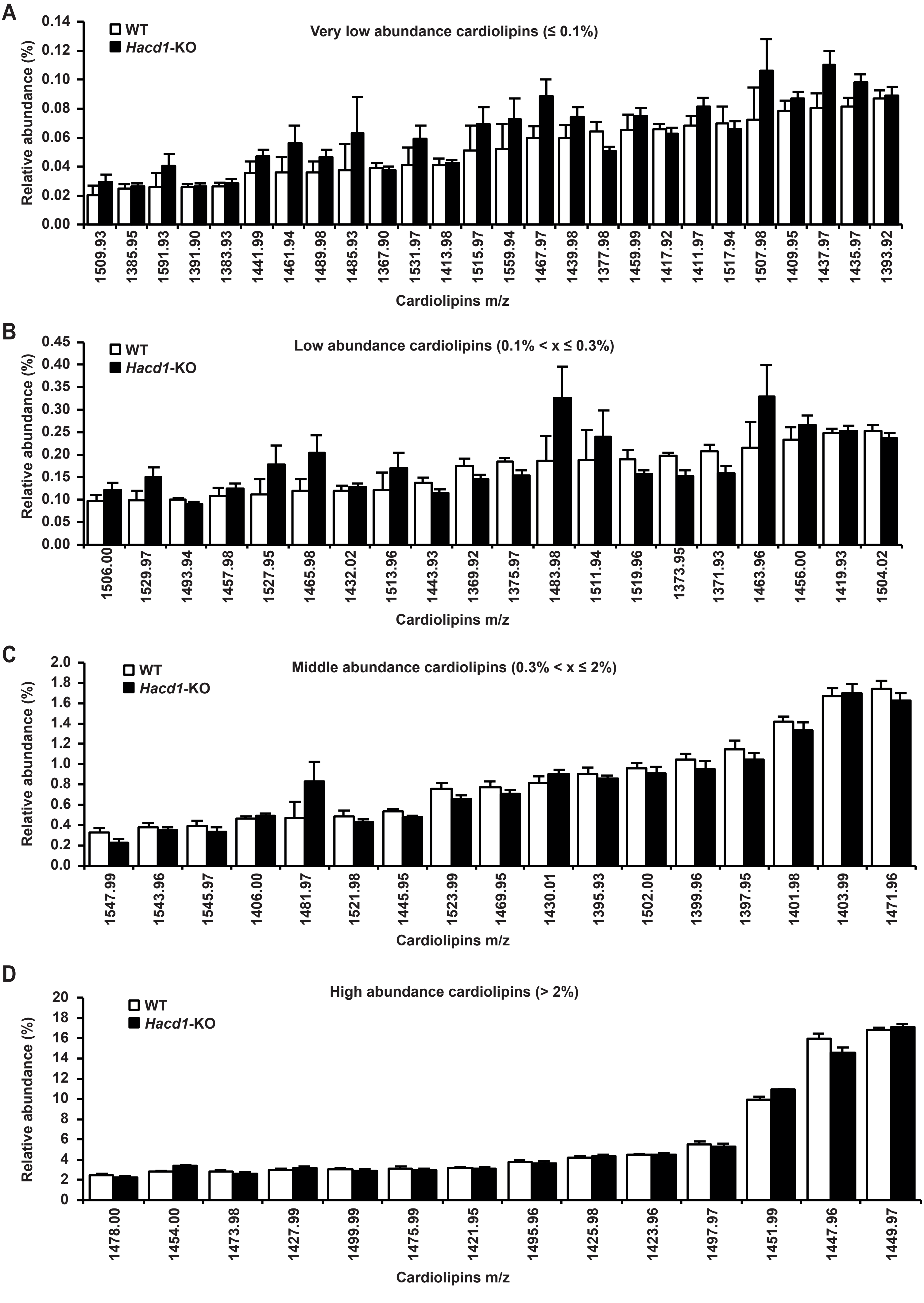


**Figure S6 Prola**

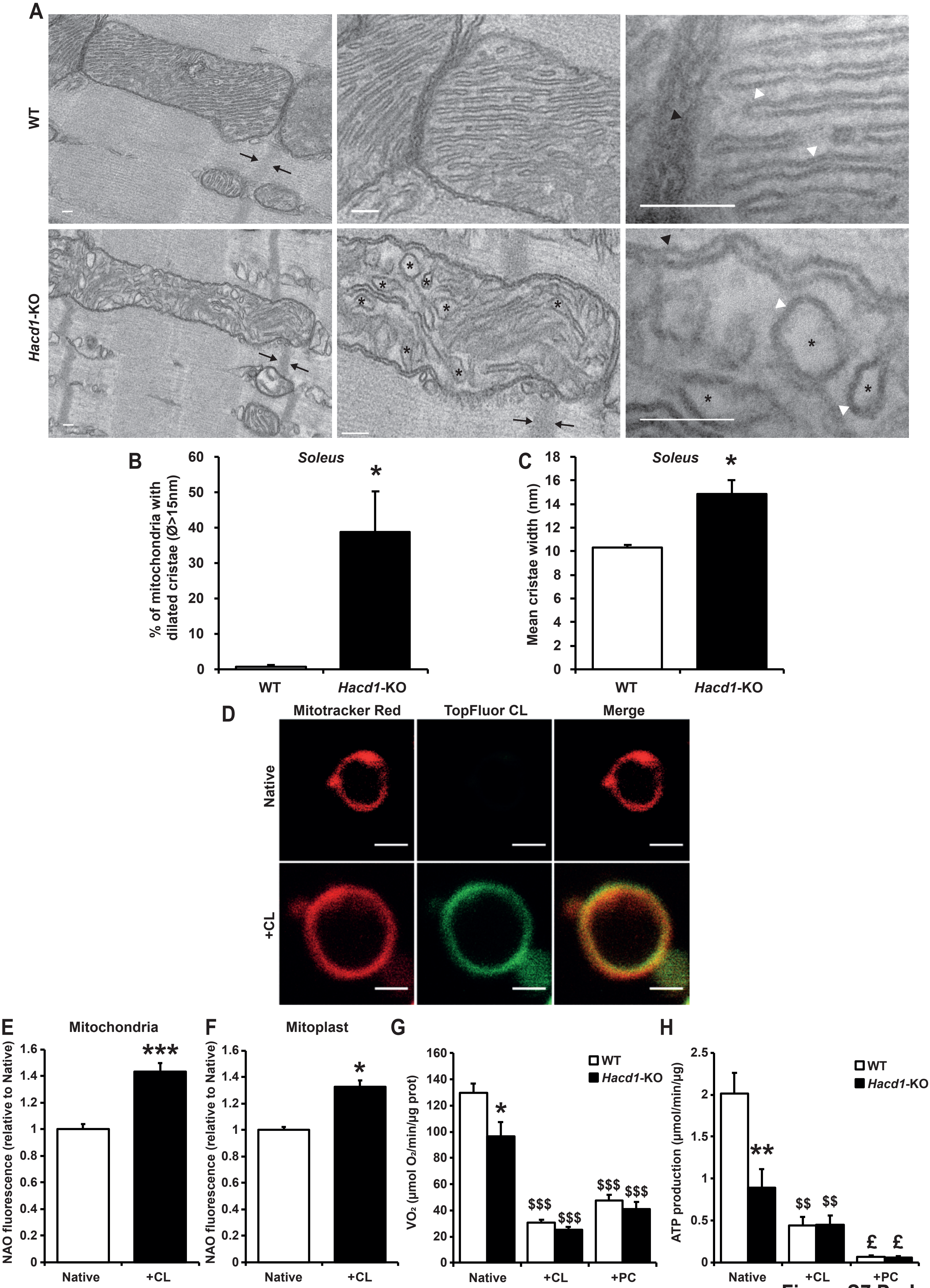


Figure S7 Prola

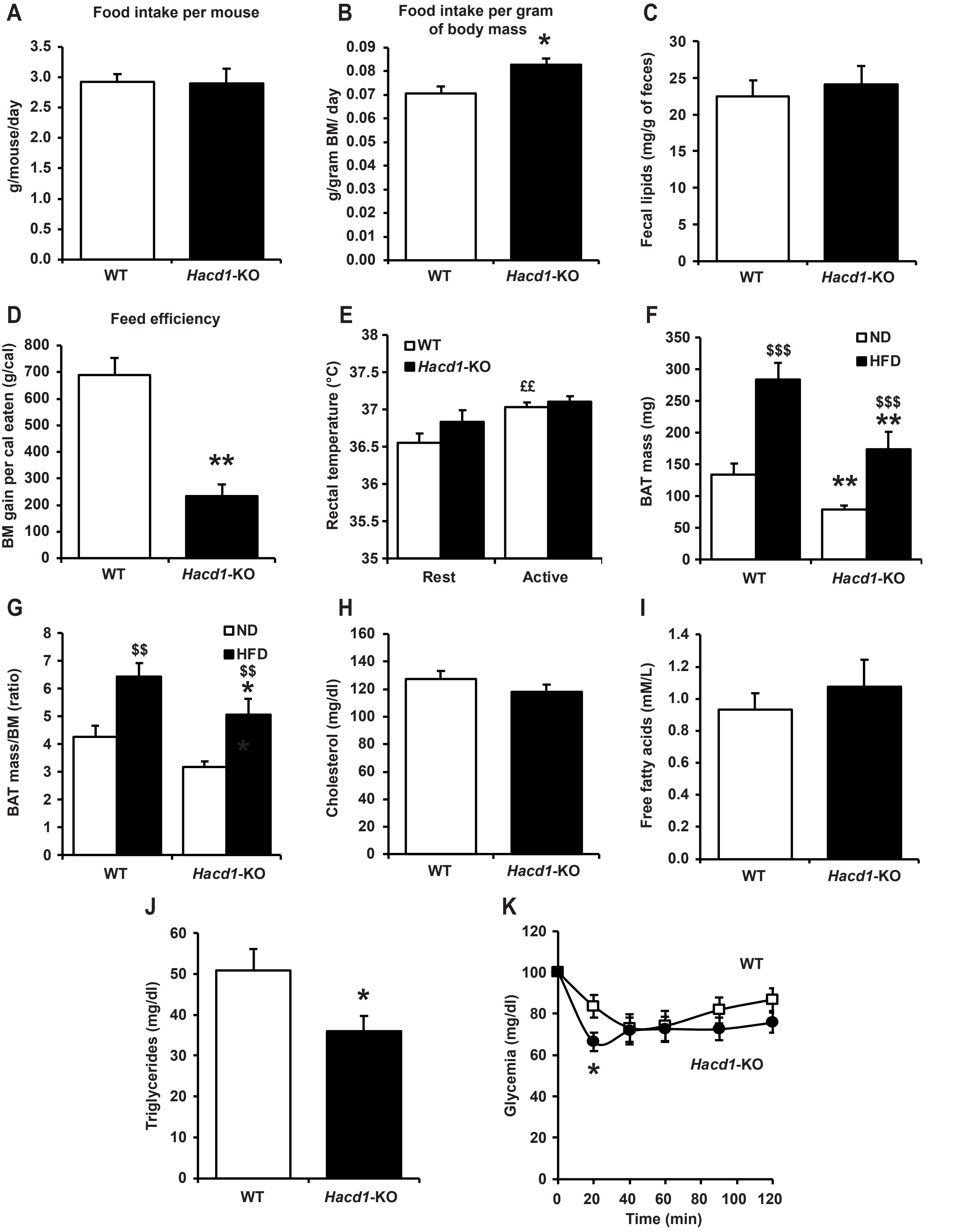


Figure S8 Prola
